# The analytical Flory random coil is a simple-to-use reference model for unfolded and disordered proteins

**DOI:** 10.1101/2023.03.12.531990

**Authors:** Jhullian J. Alston, Garrett M. Ginell, Andrea Soranno, Alex S. Holehouse

## Abstract

Denatured, unfolded, and intrinsically disordered proteins (collectively referred to here as unfolded proteins) can be described using analytical polymer models. These models capture various polymeric properties and can be fit to simulation results or experimental data. However, the model parameters commonly require users’ decisions, making them useful for data interpretation but less clearly applicable as stand-alone reference models. Here we use all-atom simulations of polypeptides in conjunction with polymer scaling theory to parameterize an analytical model of unfolded polypeptides that behave as ideal chains (ν = 0.50). The model, which we call the analytical Flory Random Coil (AFRC), requires only the amino acid sequence as input and provides direct access to probability distributions of global and local conformational order parameters. The model defines a specific reference state to which experimental and computational results can be compared and normalized. As a proof-of-concept, we use the AFRC to identify sequence-specific intramolecular interactions in simulations of disordered proteins. We also use the AFRC to contextualize a curated set of 145 different radii of gyration obtained from previously published small-angle X-ray scattering experiments of disordered proteins. The AFRC is implemented as a stand-alone software package and is also available via a Google colab notebook. In summary, the AFRC provides a simple-to-use reference polymer model that can guide intuition and aid in interpreting experimental or simulation results.

## INTRODUCTION

Proteins are finite-sized heteropolymers, and the application of polymer physics has provided a useful toolkit for understanding protein structure and function^1–9^. In particular, there has been significant interest in unfolded proteins under both native and non-native conditions^2,10–17^. Depending on the experimental techniques employed, a variety of polymeric properties can be measured, including the radius of gyration (R_g_), the hydrodynamic radius (R_h_), the end-to-end distance (R_e_), and the apparent scaling exponent (ν_app_). These and many other parameters can be calculated directly from all-atom simulations, and the synergy of simulation and experiment has provided a powerful approach for constructing large ensembles of unfolded proteins for greater insight into the unfolded state^15,18–28^.

Polymers can be described in terms of scaling laws, expressions that describe how chain dimensions vary as a function of chain length^29–31^. Polymer scaling laws typically have the format D = R_0_N^ν^. Here, D reports on chain dimensions, R_0_ is a prefactor in units of spatial distance, and N is the number of monomers, which in the case of proteins is typically written in terms of the number of amino acids. ν (or, more accurately, ν_app_ when applied to finite-sized heteropolymers like proteins) is the (apparent) Flory scaling exponent. In principle, ν_app_ lies between 0.33 (as is obtained for a perfect spherical globule) and 0.59 (as obtained for a self-avoiding chain). However, for finite-sized polymers, values beyond 0.59 can be obtainable for self-repulsive chains ^32–34^. The applicability of polymer scaling laws to describe real proteins assumes they are sufficiently long to display *bona fide* polymeric behavior and that they are sufficiently self-similar over a certain length scale, analogous to fractals. While this assumption often holds true, it is worth noting that sequence-encoded patterns in specific chemistries and/or secondary structure can lead to deviations from homopolymer-like behavior ^18,35–37^.

To what extent do polymer scaling laws apply to real proteins? For denatured polypeptides, Kohn *et al*. reported the ensemble-average radius of gyration using the scaling expressions R_g_ = 1.927N^0.598 11^. This result provides strong experimental evidence to support a model whereby denaturants unfolded proteins by uniformly weakening intramolecular protein-protein interactions^1^. A value for ν_app_ of 0.598 also agrees with the previously reported value of 0.57 by Wilkins *et al*. and earlier work by Damaschun^1,10,12^. In short, under strongly denaturing conditions, proteins appear to behave as polymers in a good solvent ^1,32,38–41^.

For proteins under native or native-like conditions, the apparent scaling exponents obtained for unfolded polypeptides are more variable. Marsh and Forman-Kay reported an average scaling expression of R_h_ = 2.49N^0.509^, for a set of intrinsically disordered proteins, while Bernadó and Svergun found a similar average relationship in R_g_ = 2.54N^0.52 42,43^. More recently, various means to estimate ν_app_ for individual proteins have enabled values of ν_app_ between 0.42 and 0.60 to be measured for a wide range of unfolded proteins of different lengths and compositions^15,18,23,25,39,40,44–46^. An emerging consensus suggests that ν_app_ depends on the underlying amino acid sequence^2,17,47^. If sequence-encoded chemical biases enable intramolecular interactions, then ν_app_ may be lower than 0.5. Notably, despite clear conceptual limitations, the physics of homopolymers remains a convenient tool through which unfolded proteins can be assessed^15,18,36,37,48^.

Given the variety in scaling exponents for unfolded proteins under native conditions, we felt that a sequence-specific reference model would be helpful for the field. Such a model could provide a touchstone for experimentally measurable polymeric parameters, including intermolecular distances, the radius of gyration, the end-to-end distance, and the hydrodynamic radius. Similarly, such a model would provide a simple reference state with which simulations could be directly compared and used to identify sequence-specific effects. Finally, a standard reference model could offer an easy way to compare unfolded proteins of different lengths to assess if they behave similarly despite different absolute dimensions.

Here, we perform sequence-specific numerical simulations for polypeptides as an ideal chain, so-called Flory Random Coil (FRC) simulations^2,31^. Under these conditions, chain-chain, chain-solvent, and solvent-solvent interactions are all equivalent, no long-range excluded volume contributions are included, and as such, the polypeptide behaves as a Gaussian chain with ν_app_ = 0.5. Because our FRC implementation minimizes finite-chain artifacts, we can parameterize an analytical, sequence-specific model using standard approaches from scaling theory, a model we call the Analytical Flory Random Coil (AFRC). This model enables the calculation of distance distributions for the end-to-end distance and the radius of gyration, as well as a variety of additional parameters that become convenient for the analysis of all-atom simulations and experiments.

The AFRC is not a predictor of unfolded protein dimensions. Those dimensions depend on the complex interplay of chain:chain and chain:solvent interactions, which are themselves determined by sequence-encoded chemistry^49–53^. Instead, the AFRC provides a simple reference state that can aid in interpreting experimental and computational results without needing information other than the protein sequence. The AFRC is implemented in a stand-alone Python package and is also provided as a simple Google Colab notebook. We demonstrate the utility of this model by comparing experimental data and computational results.

The remainder of this paper is outlined as follows. First, we discuss the implementation details of the model, including a comparison against existing polymer models. Next, we analyzed previously published all-atom simulations to demonstrate how the AFRC can identify signatures of sequence-specific intramolecular interactions in disordered ensembles. Finally, we use the AFRC model to re-interpret previously reported small-angle X-ray scattering data of intrinsically disordered proteins.

## METHODS AND RESULTS

### Implementation of a numerical model for sequence-specific ideal chain simulations

We used a Monte Carlo-based approach to construct sequence-specific atomistic ensembles of polypeptides as ideal chains. All-atom simulations with all non-bonded and solvation interactions scaled to zero were performed using a modified version of the CAMPARI Monte Carlo simulation engine using bond lengths and atomic radii defined by the ABSINTH-OPLS forcefield^2,54,55^. We modified CAMPARI to reproduce Flory’s rotational isomeric state approximation^31,56^. In this method, an initial conformation of the polypeptide is randomly generated. Upon each Monte Carlo step, a residue is randomly selected, the backbone dihedrals are rearranged to one of a subset of allowed residue-specific psi/phi values (i.e., specific isomeric states), and the chain is rearranged accordingly (**Fig. 1A, B)**. Allowed phi/psi values are selected from a database of residue-specific allowed values as determined by all-atom simulations of peptide units, with the associated Ramachandran maps shown in **Fig. S1**. Importantly, the Monte Carlo moves in these simulations approach are rejection-free. That is, only allowed phi/psi angles are proposed, and no consideration of steric overlap in the resulting conformation is given. The ensemble generated by these simulations is referred to as the Flory Random Coil (FRC, **Fig. 1C**) and has been used as a convenient reference frame for comparing simulations of disordered and unfolded polypeptides for over a decade (as reviewed by Mao *et al*.^*2*^)^15,57–61^.

**Figure 1:**
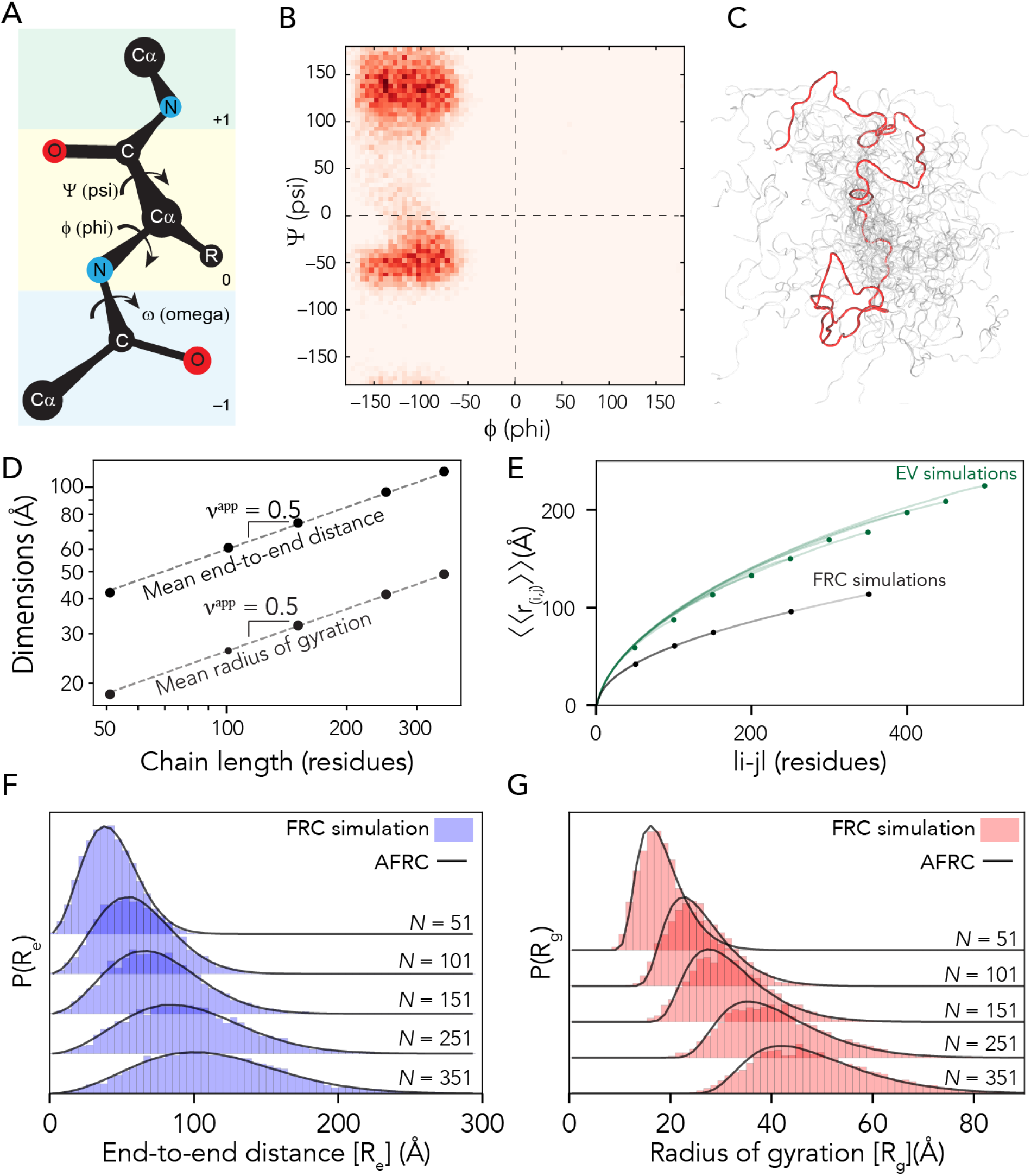
The AFRC is a pre-parameterized polymer model based on residue-specific polypeptide behavior. **A**. Schematic of the amino acid dihedral angles. **B**. Ramachandran map for alanine used to select acceptable backbone conformations for the FRC simulations. All twenty amino acids are shown in **Fig. S1. C**. Graphical rendering of an FRC ensemble for a 100-residue homopolymer. **D**. Flory Random Coil (FRC) simulations performed using a modified version of the ABSINTH implicit model and CAMPARI simulation engine yield ensembles that scale as ideal chains (i.e., R_e_ and R_g_ scale with the number of monomers to the power of 0.5). **E**. Internal scaling profiles for FRC simulations and Excluded Volume (EV) simulations for poly-alanine chains of varying lengths (filled circles demark the end of profiles for different polymer lengths). Internal scaling profiles map the average distance between all pairs of residues |i-j| apart in sequence space, where i and j define two residues. This double average reports on the fact we average over both all pairs of residues that are |i–j| apart and do so over all possible configurations. EV simulations show a characteristic tapering (“dangling end” effect) for large values of |i–j|. All FRC simulation profiles superimpose on top of one another, reflecting the absence of finite chain effects. **F**. Histograms of end-to-end distances (blue) taken from FRC simulations vs. corresponding probability density profiles generated by the Analytical FRC (AFRC) model (black line) show excellent agreement. **G**. Histograms of radii of gyration (red) taken from FRC simulations vs. corresponding probability density profiles generated by the AFRC model (black line) also show excellent agreement.

FRC simulations enable the construction of ensembles where each amino acid exists in a locally allowed configuration, yet no through-space interactions occur. This has two important implications for the construction of an ideal chain model. Firstly, each monomer has no preference for chain:chain vs. chain:solvent interactions (each monomer is “agnostic” to its surroundings). As a result, both internal and global dimensions show scaling behavior with an apparent scaling exponent (ν_app_) of 0.5 (**Fig. 1D**), analogous to that of a polymer in a theta solvent. Secondly, terminal residues sample conformational space in the same way as residues internal to the chain (**Fig. S2**). This means that end-effects that emerge finite-chain effects are not experienced in terms of end effects (**Fig. 1E**). This is in contrast to finite-sized self-avoiding chains, in which internal scaling profiles reveal a noticeable and predictable “dangling end” finite-chain effect (**Fig 1E, Fig. S2**). In summary, FRC simulations enable us to generate ensembles at all-atom resolution that are nearly fully approximations of ideal chains, reproducing the behavior of a hypothetical “ideal” polypeptide.

### Constructing an analytical description of the Flory Random Coil

Our FRC ensembles enable the calculation of a range of polymeric properties, including inter-residue distances, inter-residue contact probabilities, the hydrodynamic radius, or the radius of gyration. Comparing these properties with experiments or simulations is often convenient, offering a standard reference frame for normalization and biophysical context^2,15,17,36,37^. However, performing and analyzing all-atom simulations with CAMPARI necessitates a level of computational sophistication that may make these calculations inaccessible to many scientists. To address this, we next sought to develop a set of closed-form analytical expressions to reproduce these properties and implement them as an easy-to-use package available both locally and – importantly – via a simple web interface (Google colab notebook).

FRC simulations generate ensembles that – by definition – reproduce the statistics expected for an ideal chain. As mentioned, polymer scaling behavior generally takes the form;

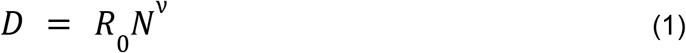

For an ideal chain, ν_app_ should not depend on the amino acid sequence (as all chains should scale with ν_app_ = 0.5). However, the prefactor R_0_ can and will show sequence dependence. As such, computing polymeric properties from sequences necessitates a means to calculate sequence-specific prefactor values. Prefactor values were parameterized using homopolymer simulations of each amino acid (see supplementary information). The inter-residue distance prefactor A_0_ was parameterized by fitting internal scaling profiles using equation (2);

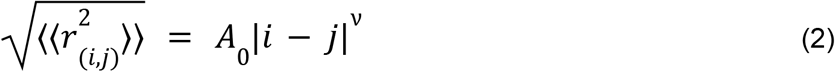

In equation 2, |i–j| is the number of residues between residues at position i and j, the left-hand-side reports on the root-mean-square (RMS) distance between residues i and j in the chain, ν is the scaling exponent (in our case this is equal to 0.5), and A_0_ is a prefactor for which we can directly solve for. The double angle brackets around the RMS distance reflect the fact we are averaging over all pairs of residues that are |i–j| apart and doing so for all chain configurations. Plotting |i–j| vs. the RMSD generates the internal scaling profile shown in **Fig. 1E**. By fitting homopolymers of the 20 amino acids, a set of residue-specific A_0_ prefactors was determined, as listed in **Supplementary Table 1**.

For our homopolymers, we can calculate the root-mean-squared end-to-end distances using equation (3);

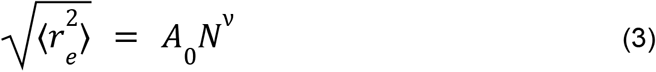

From this, we can then use the standard function for P(r) of a Gaussian chain to calculate the end-to-end distance distribution;

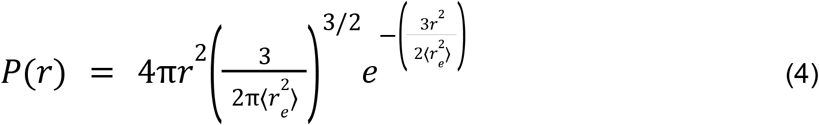

After determining residue-specific A_0_, a comparison of analytical and numerical simulation distributions show excellent agreement when homopolymer end-to-end distance distributions are compared between FRC simulations and the AFRC-derived values (**Fig. 1f**).

We next took a similar route to define the radius of gyration (R_g_) distribution. While no closed-form solution for the distribution of the radius of gyration exists, Lhuillier previously defined a closed-form approximation for this distribution for a fractal chain^62^;

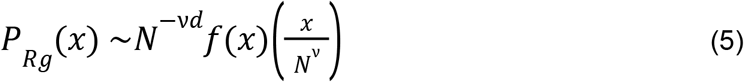

Where;

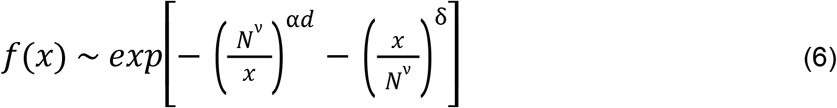

And the variables α and δ are defined as:

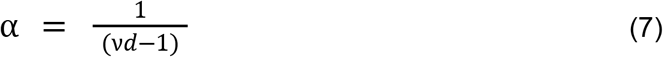

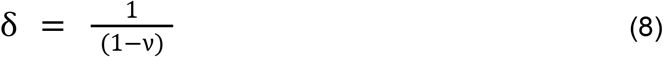

Here, x represents the distance in some arbitrary units (written as such to avoid confusion with r, which represents the distance in Angstroms [Å]), N and ν again represent the total number of residues and the scaling exponent (0.5.), while d is the dimensionality (d=3). This allows us to calculate α and δ exactly, given ν is fixed at 0.5. Consequently, we can recast equation 5 into units of Å using a sequence-specific normalization factor (X_0_);

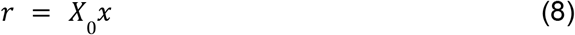

To calculate X_0_, we fit numerically-generated P(R_g_) distributions from homopolymer simulations with a series of analytically generated distributions to identify the best-fitting amino acid-specific X_0_ values. These prefactors are listed in **Supplementary Table 1**. As with the end-to-end distances, a comparison of numerically-generated P(R_g_) with analytically-generated P(R_g_) values are in extremely good agreement (**Fig. 1g**). Comparing ensemble average end-to-end distance and radii of gyration for homopolymers of all 20 amino acids in lengths from 50 to 350 amino acids revealed a Pearson correlation coefficient of 0.999 and a root mean square error (RMSE) of 0.8 Å and 0.3 Å for the end-to-end distance and radius of gyration, respectively (**Fig. S2**).

With analytical expressions for computing the end-to-end distance and radius of gyration probability distributions in hand, we can calculate additional polymeric properties. Given the fractal nature of the Flory Random Coil and the absence of end effects, we can calculate all possible inter-residue distances and, correspondingly, contact frequencies between pairs of residues (**Fig. 2a, b**). Similarly, using either the Kirkwood-Riseman equation or a recently derived empirical relationship, we can compute an approximation for the ensemble-average hydrodynamic radius^63–65^. In summary, the AFRC offers an analytic approach for calculating sequence-specific ensemble properties for unfolded homopolymers.

**Figure 2.**
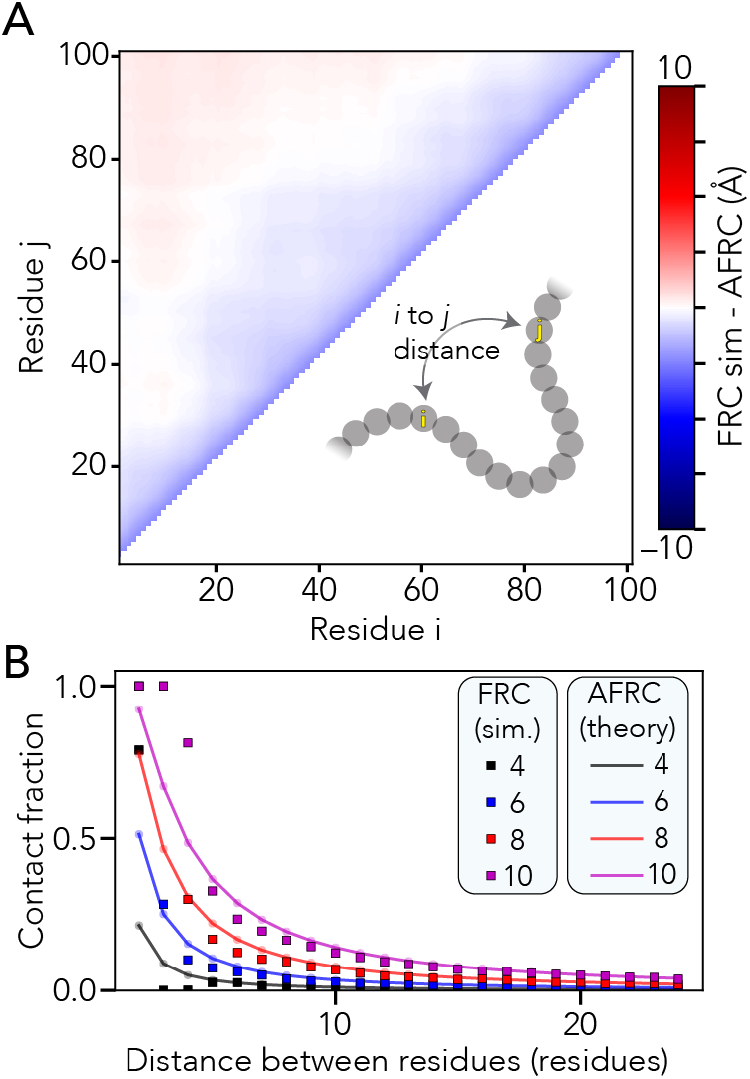
The AFRC enables the calculation of intra-residue distance distributions and expected distance-dependent contact fractions. **A**. We compared all-possible mean inter-residue distances obtained from FRC simulations with predictions from the AFRC. The maximum deviation across the entire chain is around 2.5 Å, with 92% of all distances having a deviation of less than 1 Å. **B**. Using the inter-residue distance, we can calculate the average fraction of an ensemble in which two residues are in contact (i.e., within some threshold distance). Here, we assess how that fractional contact varies with the contact threshold (different lines) and distance between the two residues. The AFRC does a somewhat poor job of estimating contact fractions for pairs of residues separated by 1,2 or 3 amino acids due to the discrete nature of the FRC simulations vis the continuous nature of the Gaussian chain distribution. However, the agreement is excellent above a sequence separation of three or more amino acids, suggesting that the AFRC offers a reasonable route to normalize expected contact frequencies.

### Generalization to heteropolymers

Our parameterization has thus far focused exclusively on homopolymer sequences. However, Flory’s rotational isomeric state approach requires complete independence of each amino residue^31,56^. Consequently, we expected the prefactor associated with a given heteropolymer to reflect a weighted average of prefactors taken from homopolymers, where the sequence composition determines the weights.

To test this expectation, we compared numerical simulations with AFRC predictions for a set of different polypeptide sequences finding excellent agreement in both end-to-end distances and radii of gyration (**Fig. 3a, b** and **Fig. S3**). Similarly, given the absence of end-effects, our analytical end-to-end distance expression works equally well for intramolecular distances in addition to the end-to-end distance. To assess this, we compared internal scaling profiles between FRC simulations and AFRC predictions (**Fig. 3c**). These profiles compare the ensemble average distance between each possible inter-residue distance and offer a convenient means to assess both short and long-range intramolecular distances. We performed FRC simulations for 320 different polypeptide sequences ranging in length from 10 to 500 amino acids with a systematic variation in amino acid composition. Across all internal scaling profile comparisons between FRC and AFRC simulations, the overall average RMSE was 0.5 Å, with almost all (92%) of individual comparisons revealing an RMSE under 1 Å (**Fig. 3D**). Similarly, the Pearson’s correlation coefficient between internal scaling profiles for FRC vs. AFRC for all ten-residue chains was 0.9993, which was the worst correlation across all lengths (**Fig. S4**). In summary, the AFRC faithfully reproduces homo- and hetero-polymeric dimensions for polypeptides under the FRC assumptions.

**Fig. 3.**
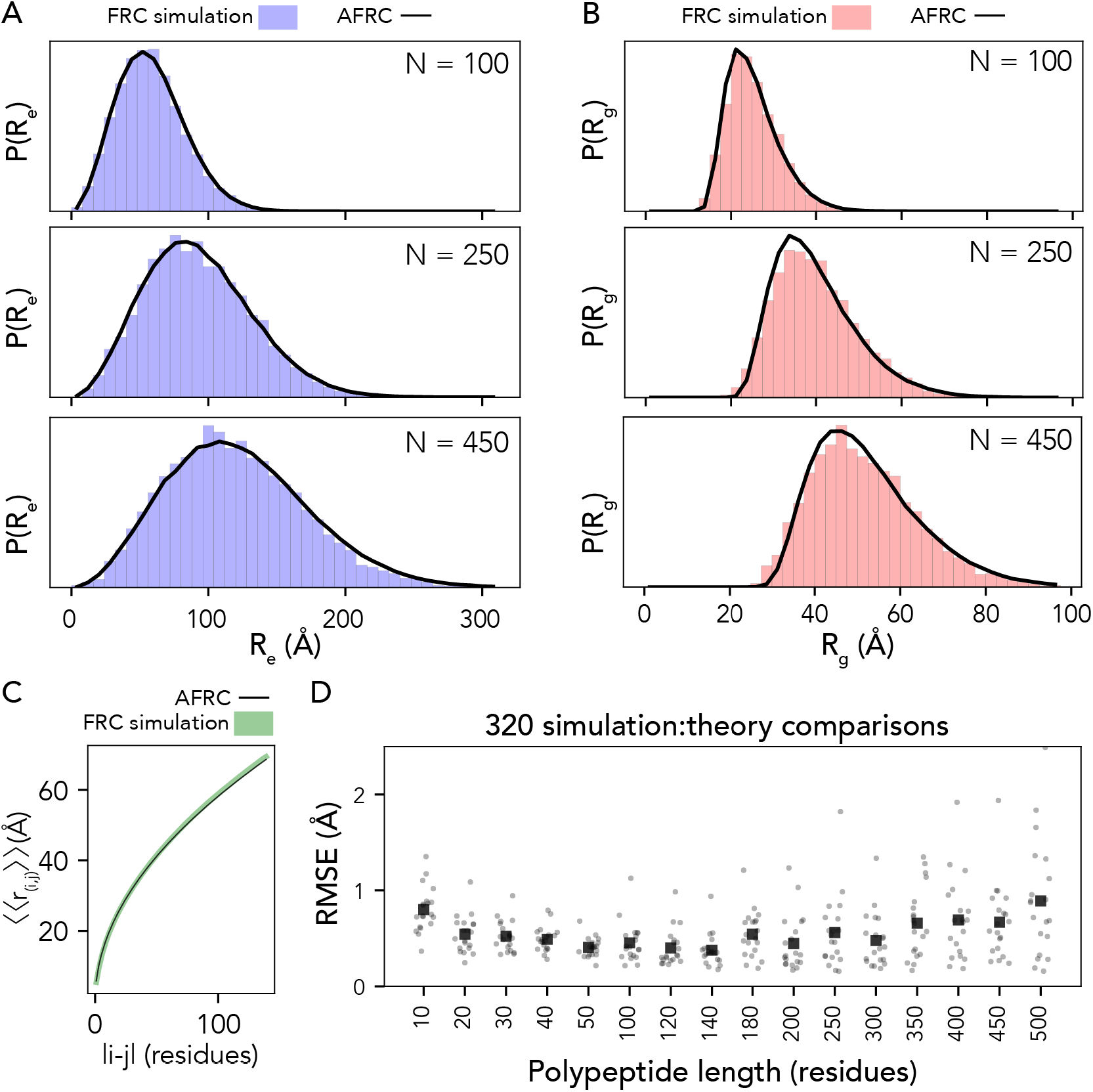
The AFRC generalizes to arbitrary heteropolymeric sequences with the same precision and accuracy as it does for homopolymeric sequences. **A**. Representative examples of randomly polypeptide heteropolymers of lengths 100, 250, and 450, comparing the AFRC-derived end-to-end distance distribution (black curve) with the empirically-determined end-to-end distance histogram from FRC simulations (blue bars). **B**. The same three polymers, as shown in A, now compare the AFRC-derived radius of gyration distance distribution (black curve) with the empirically-determined radius of gyration histogram from FRC simulations (blue bars). **C** Comparison of AFRC vs. FRC simulation-derived internal scaling profiles for a 150-amino acid random heteropolymer. The deviation between FRC and AFRC for these profiles offers a measure of agreement across all length scales. **D** Comparison of root-mean-square error (RMSE) obtained from internal scaling profile comparisons (i.e., as shown in C) for 320 different heteropolymers straddling 10 to 500 amino acids in length. In all cases, the agreement with theory and simulations is excellent.

### Comparison with existing polymer models

For completeness, we compared the end-to-end distance distributions obtained from several other polymer models used throughout the literature for describing unfolded and disordered polypeptides. Previously-used polymer models offer a means to analytically fit experimental or computational results and benefit from taking one (or more) parameters that define the model’s behavior. While the AFRC does not enable fitting to experimental or simulated data, it only requires an amino acid sequence as input. With this in mind, the AFRC serves a fundamentally different purpose than commonly used models.

We wondered if dimensions obtained from the AFRC would be comparable with dimensions obtained from other polymer models when using parameters used previously in the literature. We compared distributions obtained from the worm-like chain (WLC), the self-avoiding walk (SAW) model, and a recently-developed ν-dependent self-avoiding walk (SAW-ν)^23,66^. For the WLC model, we used a persistence length of 3.0 Å and an amino acid size of 3.8 Å (such that the contour length, l_c_, is defined as N×3.8^66^). For the SAW model, we used a scaling prefactor of 5.5 Å (i.e., assuming < R_e_> = 5.5N^0.598^)^23^,^32^,^66^. Finally, for SAW-ν, we computed distributions using a prefactor of 5.5 Å and using several different ν values^6^,^23^. These values were chosen because previous studies have used them to describe intrinsically disordered proteins.

**Fig. 4A** shows comparisons of the AFRC distance distribution obtained for a 100-mer polyalanine (A_100_) vs. the WLC and SAW (top) and vs. ν-dependent distributions (bottom). The AFRC is slightly more expanded than the WLC model using the parameters provided, although the persistence length can, of course, be varied to explore more compact (lower *l*_*p*_) or more extended (higher *l*_*p*_) distributions (**Fig. S6A**). The AFRC is substantially more compact than the SAW model. The comparison with the SAW model is important, as with a prefactor of 5.5 Å the SAW model describes a polypeptide as a self-avoiding random coil (ν=0.588), whereas the AFRC describes a polypeptide as an ideal chain (ν = 0.5), such that we should expect the SAW to be more expanded than the AFRC. Finally, in comparing the AFRC with the SAW-ν model, we find that the AFRC distribution falls almost completely top of the ν = 0.50 distribution. This indicates that both models arrive at nearly identical distance distributions despite being developed independently. This result is both confirmatory and convenient, as it means the AFRC and SAW-ν models can be used to analyze the same data without concern for model incompatibility.

**Fig. 4.**
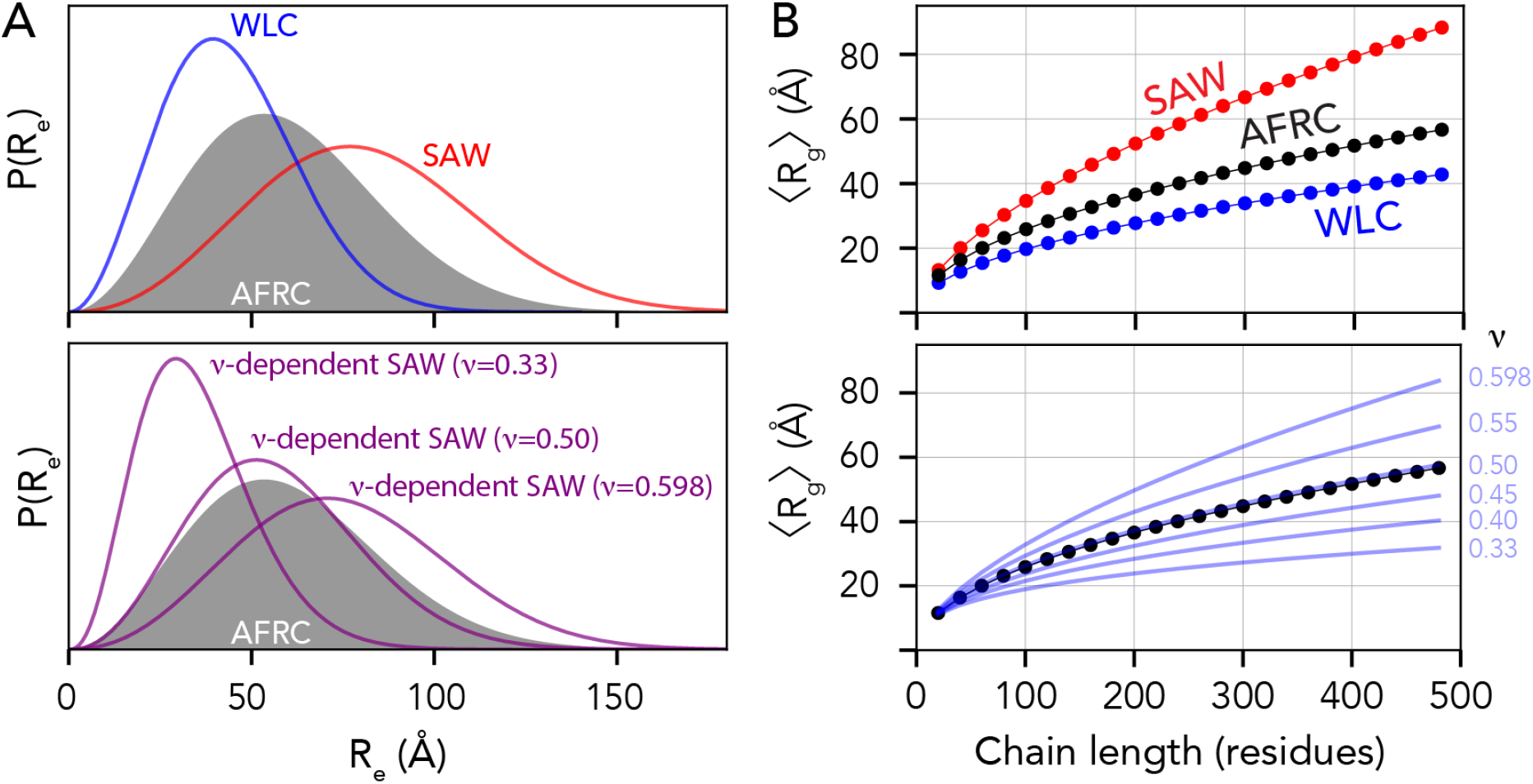
The AFRC is complementary to existing polymer models. **(A)** Comparison of end-to-end distance distributions for several other analytical models, including the Wormlike Chain (WLC), the self-avoiding walk (SAW), and the ν-dependent SAW model (SAW-ν). The AFRC behaves like a ν-dependent SAW with a scaling exponent of 0.5. **(B)** Comparisons of ensemble-average radii of gyration as a function of chain length for the same sets of polymer models. The AFRC behaves as expected and again is consistent with a ν-dependent SAW with a scaling exponent of 0.5.

We emphasize that this comparison with the existing polymer model is not presented to imply the AFRC is better than existing models but to highlight their compatibility. One can tune input parameters for all three models to arrive at qualitatively matching end-to-end distributions (**Fig. S6B**). The major difference between these three models and the AFRC is simply that the AFRC requires only amino acid sequence as input, making it a convenient reference point. For completeness, all four models are implemented in our Google colab notebook.

We also compared ensemble-average radii of gyration obtained from the various models with those obtained from the AFRC. While the WLC, SAW, and SAW-ν models do not provide approximate closed-form solutions for the radius of gyration distribution, they do enable an estimate of the ensemble-average radius of gyration to be calculated^23,66^. Using the same model parameters as was used in **Fig. 4A**, the AFRC falls between the SAW and the WLC. Moreover, the AFRC radii of gyration scale almost 1:1 with the SAW-ν derived radii as a function of chain length when ν = 0.50. As such, we conclude that the AFRC is consistent with existing polymer models yet benefits from being both parameter-free (for the user) and offering full distributions for the radius of gyration and intramolecular distance distributions per-residue contact fractions, convenient properties for normalization in simulations and experiment.

### Comparison with all-atom simulations

Our work thus far has focussed on developing and testing the robustness of the AFRC. Having done this, we next sought to ask how similar (or dissimilar) distributions obtained from the AFRC are compared to all-atom simulations. We used simulations generated via all-atom molecular dynamics with the Amber99-disp forcefield and all-atom Monte Carlo simulations with the ABSINTH-OPLS forcefield^25,55,67–71^. Specifically, we examined nine different fully disordered proteins: The unfolded Drosophila Drk N-terminal SH2 domain (DrkN, 59 residues)^67,72,73^, the ACTR domain of p160 (ACTR, 71 residues)^39,40,67,74^, a C-terminal disordered subregion of the yeast transcription factor Ash1 (Ash1, 83 residues)^68^, the N-terminal disordered regions of p53 (p53, 91 residues)^71,75^, the C-terminal IDR of p27 (p26, 107 residues)^70^, the intrinsically disordered intracellular domain of the notch receptor (Notch, 132 residues)^69^, the C-terminal disordered domain of the measles virus nucleoprotein (Ntail, 132 residues)^67,76^, the C-terminal low-complexity domain of hnRNPA1 (A1-LCD, 137 residues)^25^, and full-length alpha-synuclein (asyn, 140 residues)^67,77,78^. We compared distributions for the end-to-end distance and radius of gyration for our all-atom simulations with analogous distributions generated by the AFRC (**Fig. 5**). These comparisons revealed that while the general shape of the distributions recovered from simulations was not dissimilar from the AFRC-derived end-to-end distance and radius of gyration distributions, the width and mean were often different. This is hardly surprising, given that the global dimensions of an unfolded protein depend on the underlying amino acid sequence. The ratio of the mean end-to-end distance divided by the AFRC-derived mean end-to-end distance (or the corresponding ratio for the radius of gyration) was found to range between 0.7 and 1.4. In some cases, the end-to-end distance ratio or radius of gyration ratio varied within the same protein. For example, for the 132-residue intracellular-domain IDR from Notch (Notch), the end-to-end distance ratio was 0.8 (i.e., smaller than predicted by the AFRC), while the radius of gyration ratio was 1.0. Similarly, in alpha-synuclein (Asyn), the corresponding ratios were 0.7 and 0.9, again reporting a smaller end-to-end distance than radius of gyration. As suggested previously, discrepancies in end-to-end distance vs. radius of gyration vs. expectations from homopolymer models are diagnostic of sequence-encoded conformational biases^18,35,36,79^. We also used the AFRC to calculate scaling maps. Scaling maps are non-redundant matrices of inter-residue distances obtained from simulations and normalized by the expected inter-residue distances obtained by the AFRC (**Fig. 6**)^68^. We compared these scaling maps (top left triangle of each panel) against absolute distances (bottom right triangle). This comparison highlights the advantage that using a reference polymer model offers. Long-range sequence-specific conformational biases are much more readily visualized as deviations from an expected polymer model. Moreover, the same dynamic range of values can be used for chains of different lengths, normalizing the units from Å to a unitless ratio.

**Fig. 5.**
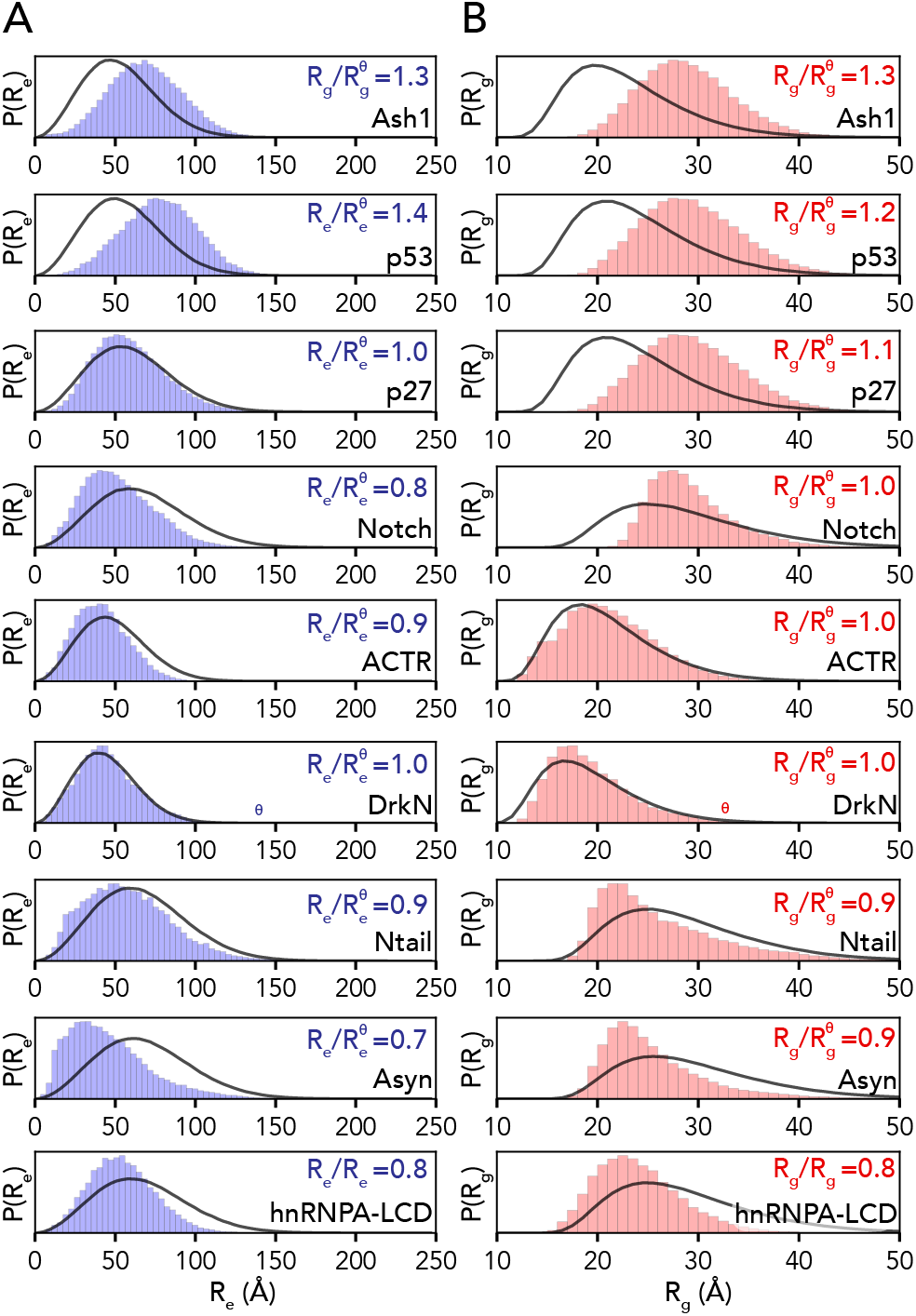
AFRC-derived distance distributions enable simulations to be qualitatively compared against a null model. **A**. Comparison of the AFRC-derived end-to-end distance distributions (black line) with the simulation-derived end-to-end distribution (blue bars) for all-atom simulations of nine different disordered proteins. **B**. Comparison of the AFRC-derived radius of gyration distributions (black line) with the simulation-derived radius of gyration distribution (red bars) for all-atom simulations of nine different disordered proteins.

**Fig. 6.**
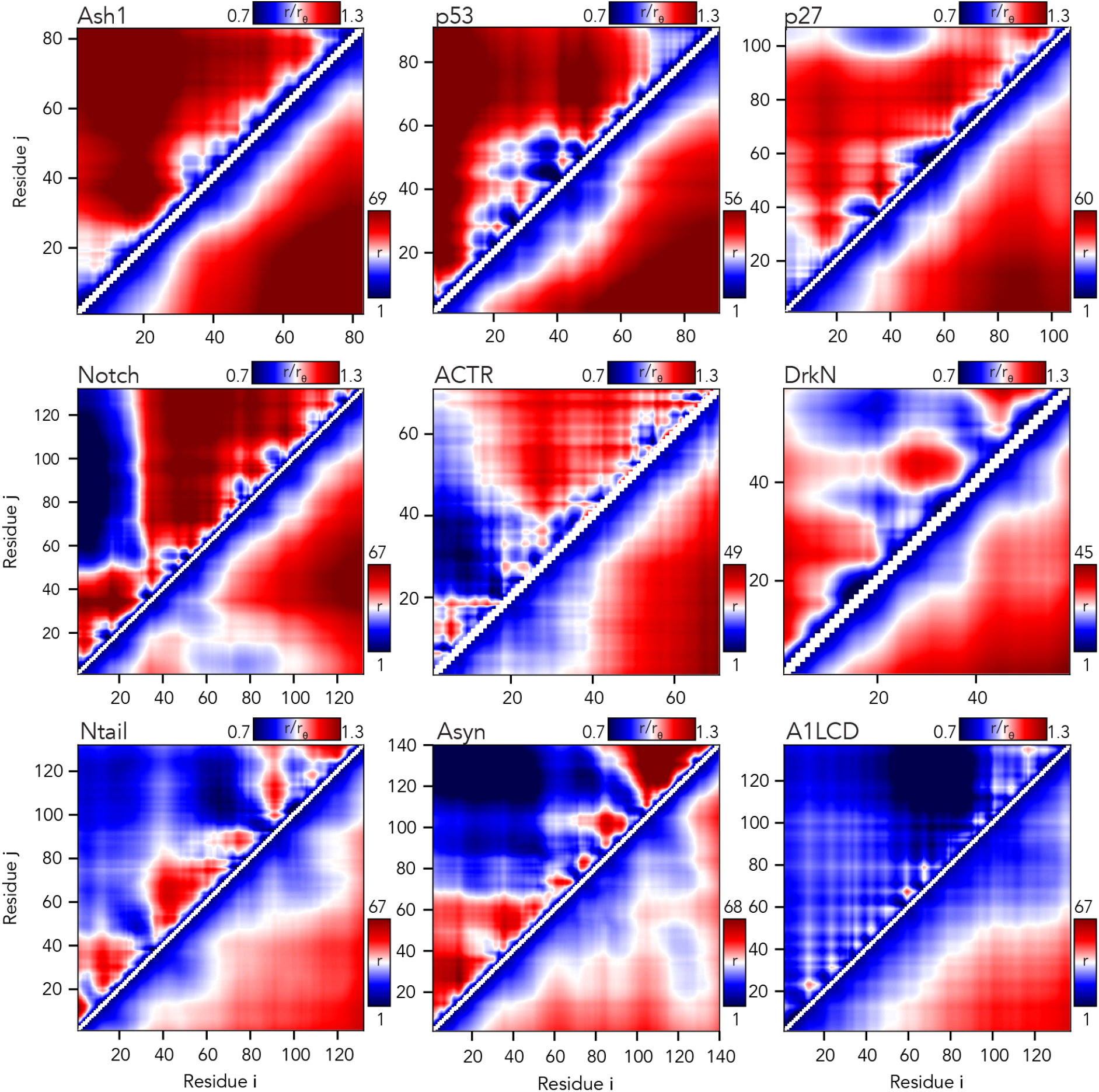
The AFRC enables a consistent normalization of intra-chain distances to identify specific sub-regions that are closer or further apart than expected. Inter-residue scaling maps (top left) and distance maps (bottom right) reveal the nuance of intramolecular interactions. Scaling maps (top left) report the average distance between each pair of residues (i,j) divided by the distance expected for an AFRC-derived distance map, providing a unitless parameter that varies between 0.7 and 1.3 in these simulations. Distance maps (bottom right) report the absolute distance between each pair of residues in angstroms. While distance maps provide a measure of absolute distance in real space, scaling maps provide a cleaner, normalized route to identify deviations from expected polymer behavior, offering a convenient means to identify sequence-specific effects. For example, in Notch and alpha-synuclein, scaling maps clearly identify end-to-end distances as close than expected. Scaling maps also offer a much sharper resolution for residue-specific effects - for example, in p53, residues embedded in the hydrophobic transactivation domains are clearly identified as engaging in transient intramolecular interactions, leading to sharp deviations from expected AFRC distances.

Returning to the notch simulations, both types of intramolecular distance analysis clearly illustrate a strong long-range interaction between the N-terminal residues 1-30 and the remainder of the sequence. The long-range interaction between chain ends influences the end-to-end distance much more substantially than it does the radius of gyration (**Fig. 6**). Similarly, in alpha-synuclein, we observed long-range interactions between the negatively charged C-terminus and the positively-charged residues 20-50, leading to a reduction in the end-to-end distance. In short, the AFRC provides a convenient approach to enable direct interrogation of sequence-to-ensemble relationships in all-atom simulations.

Finally, we calculated per-residue contact scores for each residue in our nine proteins (**Fig. 7**). These contact scores sum the length-normalized fraction of the simulation in which each residue is in contact with any other residue in the sequence^25^. While this collapses information on residue-specificity into a single number, it integrates information from the typically-sparse contact maps for IDR ensembles to identify residues that may have an outside contribution towards short (<6 Å) range molecular interactions. We and others have previously used this approach to identify “stickers” - regions or residues in IDRs that have an outsized contribution to intra- and inter-molecular interactions^25,61,80,81^.

**Fig. 7.**
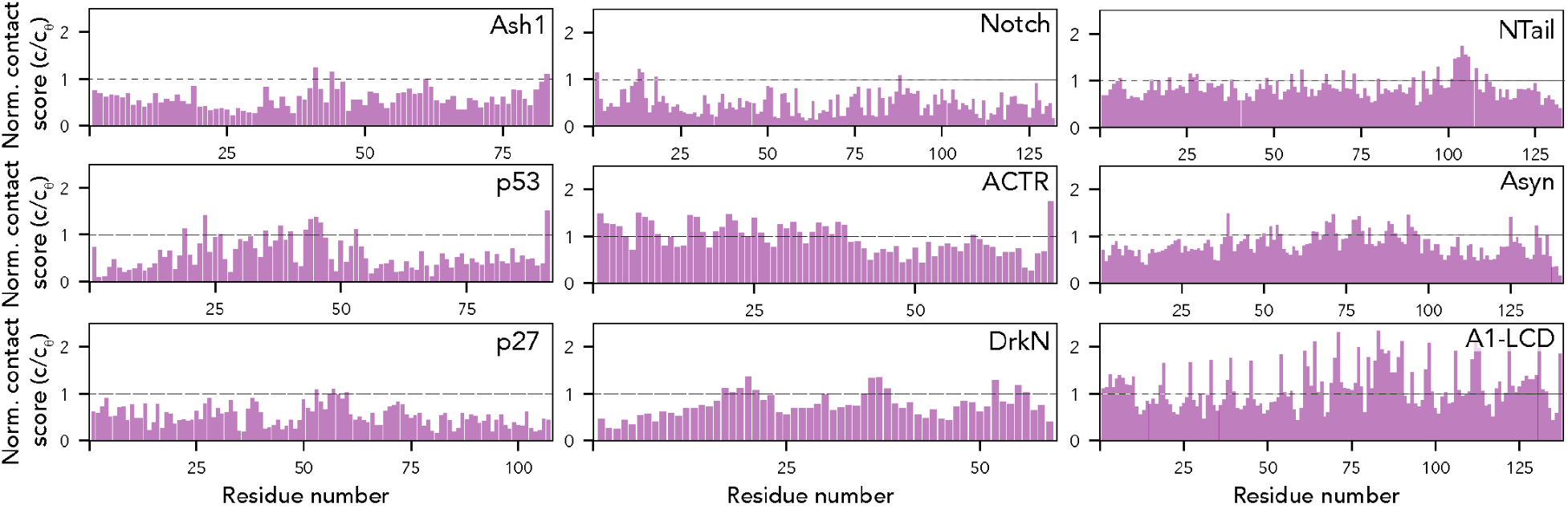
The AFRC enables an expected contract fraction to be calculated, such that normalized contact frequencies can be easily calculated for simulations. Across the nine different simulated disordered proteins, we computed the contact fraction (i.e., the fraction of simulations each residue is in contact with any other residue) and divided this value by the expected contact fraction from the AFRC model. This analysis revealed subregions within IDRs that contribute extensively to intramolecular interactions, mirroring finer-grain conclusions obtained in **Fig. 6**.

In some proteins, specific residues or subregions were identified as contact hotspots. This includes the aliphatic residues in ACTR, and hydrophobic residues in the p53 transactivation domains, in line with recent work identifying aliphatic residues as driving intramolecular interactions^61,82^. Most visually noticeable, aromatic residues in the A1-LCD appear as spikes that uniformly punctuate the sequence, highlighting their previously-identified role as evenly-spaced stickers^25^. Intriguingly, in alpha-synuclein, several regions in the aggregation-prone non-amyloid core (NAC) region (residues 61-95) appear as contact score spikes, potentially highlighting the ability of intramolecular interactions to guide regions or residues that may mediate inter-molecular interaction.

### Comparison with SAXS-derived radii of gyration

Having compared AFRC-derived parameters with all-atom simulations, we next sought to determine if AFRC-derived polymeric properties compared reasonably with experimentally-measured values. As a reminder, the AFRC is not a predictor of IDR behavior; instead, it offers a null model against which IDR dimensions can be compared. To perform a comparison with experimentally derived data, we curated a dataset of 145 examples of radii of gyration measured by small-angle X-ray scattering (SAXS) of disordered proteins. We choose to use SAXS data because SAXS-derived radii of gyration offer a label-free, model-free means to determine the overall dimensions of a disordered protein. That said, SAXS-derived measurements are not without their caveats (see discussion), and where possible, we re-analyzed primary scattering data to ensure all radii of gyration reported here are faithful and accurate. To assess our SAXS-derived radii of gyration, we calculated expected dimensions for denatured proteins, folded globular domains, or AFRC chains by fitting scaling laws with the form R_g_ = R_0_N^ν^ against different polymer models. We used a denatured-state polymer model (ν = 0.59, R_0_ = 1.98, as defined by Kohn *et al*.) and a folded globular domain model (ν = 0.33, R_0_ = 2.86, as obtained from PDBSELECT25 originally plotted by Holehouse & Pappu)<u>^11,48,83^</u>. We also calculated the AFRC-derived radii of gyration for all 145 chains and fitted a polymer scaling model to the resulting data where the only free parameter was R_0_ (ν = 0.50, R_0_ = 2.50). This analysis showed that the majority of the 145 proteins have a radius of gyration above that of the AFRC-derived radius of gyration (see discussion), with some even exceeding the expected radius of gyration of a denatured protein (**Fig. 8A**). Based on these data, we determined an empirical upper and lower bound for the biologically accessible radii of gyration given a chain length (see discussion). This threshold suggests that, for a sequence of a given length, there is a wide range of possible IDR dimensions accessible (**Fig. 8B, Fig. S5**).

**Fig 8.**
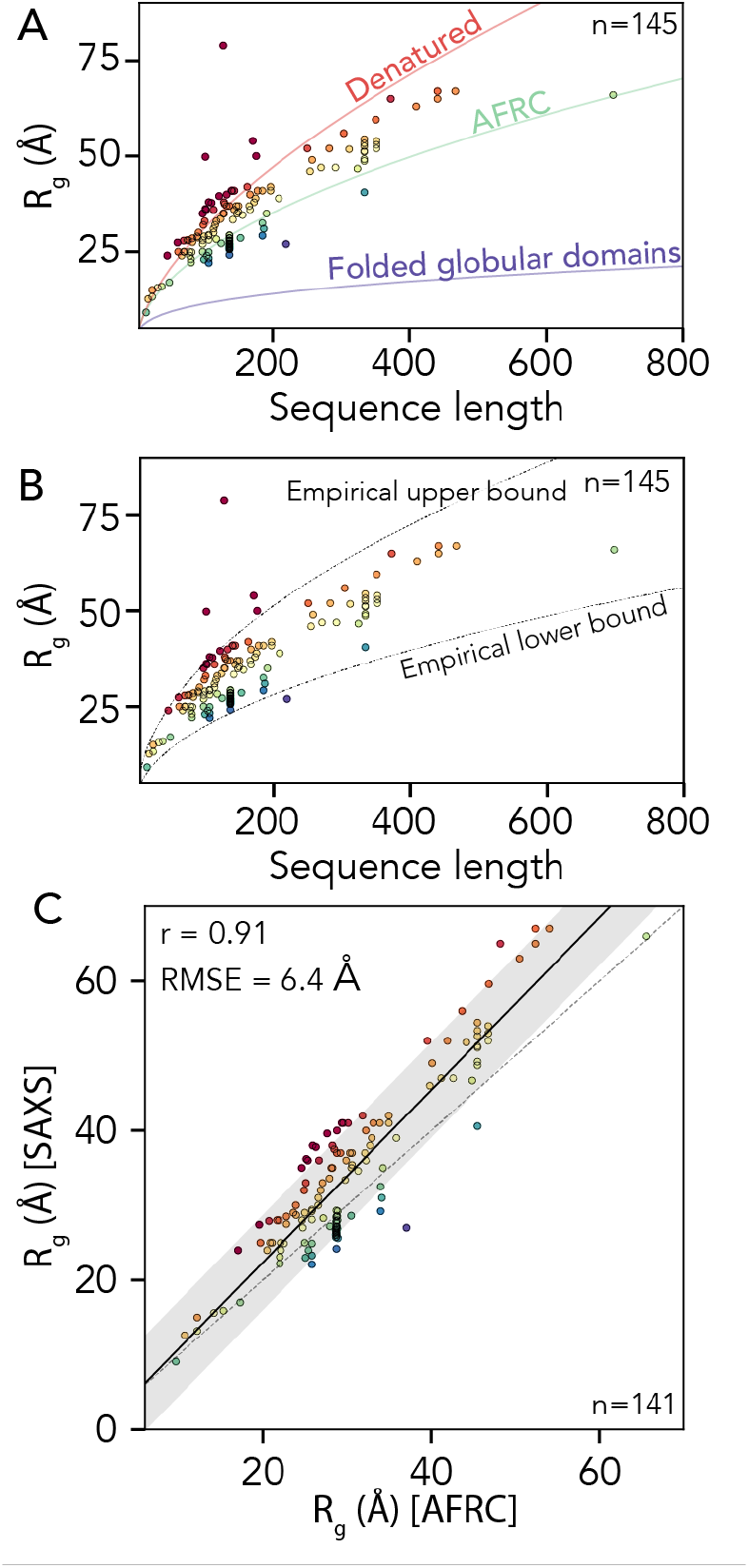
Comparison of AFRC-derived radii of gyration with experimentally-measured values. **A**. We compared 145 experimentally-measured radii of gyration against three empirical polymer scaling models that capture the three classes of polymer scaling (ν = 0.33 [globular domains], ν = 0.5 [AFRC], and ν = 0.59 [denatured state]). Individual points are colored by their normalized radius of gyration (SAXS-derived radius of gyration divided by AFRC-derived radius of gyration). **B**. The same data as in panel A with the empirically defined upper and lower bound. As with panel A, individual points are colored by their normalized radius of gyration. **C**. Comparison of SAXS-derived radii of gyration and AFRC-derived radii of gyration, as with panels A and B, individual points are colored by their normalized radius of gyration.

Finally, we wondered how well the AFRC-derived radii of gyration would correlate with experimentally-measured values. Based on the upper and lower bounds shown in **Fig. 8B**, we excluded four radii of gyration that appear to be spuriously large, leaving 141 data points. For these 141 points, we calculated the Pearson correlation coefficient (r) and the RMSE between the experimentally-measured radii of gyration and the AFRC-derived radii of gyration. This analysis yielded a correlation coefficient of 0.91 and an RMSE of 6.4 Å (**Fig. 8C**). To our surprise, these metrics outperform several established coarse-grained models for assessing intrinsically disordered proteins, as reported recently<u>^84^</u>. We again emphasize that the AFRC is not a predictor of IDR dimensions. However, we tentatively suggest that this result demonstrates that a reasonably good correlation between amino acid sequence and global dimensions can be obtained solely by recognizing that disordered proteins are flexible polymers. With this in mind, we conclude that the AFRC provides a convenient and easily-accessible control for experimentalists measuring the global dimensions of disordered proteins.

### Reference implementation and distribution

Computational and theoretical tools are only as useful as they are usable. To facilitate the adoption of the AFRC as a convenient reference ensemble, we provide the AFRC as a stand-alone Python package distributed through PyPI (**pip install afrc**). We also implemented the additional polymer modes described in **Fig. 4** with a consistent programmatic interface, making it relatively straightforward to apply these models to analyze and interpret computational and experimental data. Finally, to further facilitate access, we provide an easy-to-use Google colab notebook for calculating expected parameters for easy comparison with experiments and simulations. All information surrounding access to the AFRC model is provided at https://github.com/idptools/afrc.

## DISCUSSION & CONCLUSION

In this work, we have developed and presented the Analytical Flory Random Coil (AFRC) as a simple-to-use reference model for comparing against simulations and experiments of unfolded and disordered proteins. We demonstrated that the AFRC behaves as a truly ideal chain and faithfully reproduces homo- and hetero-polymeric inter-residue and radius of gyration distributions obtained from explicit numerical simulations. We also compared the AFRC against several previously-established analytical polymer models, showing that ensemble-average or distribution data obtained from the AFRC are interoperable with existing models. Finally, we illustrated how the AFRC could be used as a null model for comparing data obtained from simulations and from experiments.

The AFRC differs from established polymer models in two key ways. While existing models define functional forms for polymeric properties, they do not prescribe specific length scales or parameters for those models. This is not a weakness - it simply reflects how analytical models work. However, the need to provide ‘appropriate’ parameters to ensure these models recapitulate behaviors expected for polypeptides places the burden on selecting and/or justifying those parameters on the user. The AFRC combines several existing analytical models (the Gaussian chain and the Lhuillier approximation for the radius of gyration distribution) with specific parameters obtained from numerical simulations to provide a “parameter-free” polymer model defined by its reference implementation (as opposed to the mathematical form of the underlying distributions). We place parameter free in quotation marks because the freedom from parameters is at the user level - the model itself is explicitly parameterized to reproduce polypeptides dimensions. However, from the user’s perspective, no information is needed other than the amino acid sequence.

Although the AFRC was explicitly parameterized to recapitulate numerical FRC simulations, sequence-specific effects do not generally have a major impact on the resulting dimensions. For example, **Fig. S6** illustrates the radius of gyration or end-to-end distance obtained for varying lengths of poly-alanine and poly-glycine. This behavior is not a weakness of the model - it *is* the model. This relatively modest sequence dependence reflects the fact that for an ideal chain, both the second and third virial coefficients are set to zero (*i*.*e*., the integral of Mayer f-function should equal 0)^85^. As such, the AFRC does not enable explicitly excluded volume contributions to the chain’s dimensions from sidechain volume, although this is captured implicitly based on the allowed isomeric states (compare glycine to alanine in **Fig. S1**). In summary, the AFRC does not offer any new physics, but it does encapsulate previously derived physical models along with numerically-derived sequence-specific parameters to make it easy to construct null models explicitly for comparison with polypeptides.

In comparing AFRC-derived polymeric properties with those obtained from all-atom simulations, we recapitulate sequence-to-ensemble features identified previously ^25,28,67,69^. When comparing the normalized radii of gyration (R_g_^Sim^/R_g_^AFRC^), we noticed the lower and upper bounds obtained here appear to be approximately 0.8 and 1.4, respectively. To assess if this trend held true for experimentally derived radii of gyration, we calculated the normalized radii of gyration for the 141 values reported in **Fig. 8C**, recapitulating a similar range (0.8 to 1.46). Based on these values, we defined an empirical boundary condition for the anticipated range in which we would expect to see a disordered chain’s radius of gyration as between 0.8R_g_ ^AFRC^ and 1.45R_g_ ^AFRC^ (**Fig. 8B**). We emphasize this is not a hard threshold. However, it offers a convenient rule-of-thumb, such that measured radii of gyration can be compared against this value to assess if a potentially spurious radius of gyration has been obtained (either from simulations or experiments). Such a spurious value does not necessarily imply a problem, but may warrant further investigation to explain its physical origins.

Our comparison with experimental data focussed on radii of gyration obtained from SAXS experiments. We chose this route given the wealth of data available and the label-free and model-free nature in which SAXS data are collected and analyzed. Given the AFRC offers the expected dimensions for a polypeptide behaving qualitatively as if it is in a theta solvent, it may be tempting to conclude from these data that the vast majority of disordered proteins are found in a good solvent environment (**Fig. 9A**). The solvent environment reflects the mean-field interaction between a protein and its environment. In the good solvent regime, protein:solvent interactions are favored, while in the poor solvent regime protein:protein interactions are favored ^2,6,44,48^. However, it is worth bearing in mind that SAXS experiments generally require relatively high concentrations of protein to obtain reasonable signal-to-noise^43^. Recent advances in size exclusion chromatography (SEC) coupled SAXS have enabled the collection of scattering data for otherwise aggregation-prone proteins with great success^86^. However, there is still a major acquisition bias in the technical need of these experiments to work with high concentrations of soluble proteins when integrated over all existing measured data. By definition, such highly soluble proteins experience a good solvent environment. Given this acquisition bias, we remain agnostic as to whether these results can be used to extrapolate to the solution behavior of all IDRs.

**Fig 9.**
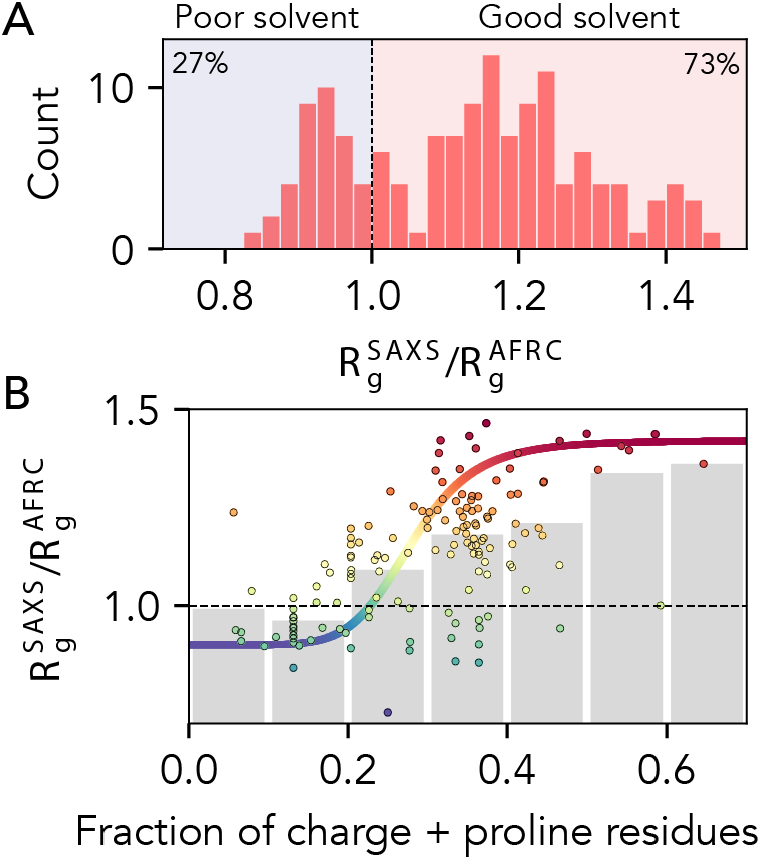
AFRC-normalized radii of gyration from experimentally-measured proteins. **A**. Histogram showing the normalized radii of gyration for 141 different experimentally-measured sequences. **B**. Comparison of normalized radii of gyration for 141 different experimentally-measured sequences against the fraction of charge and proline residues in those sequences. Individual points are colored by their normalized radius of gyration. Grey bars reflect the average radius of gyrations obtained by binning sequences with the corresponding fraction of charge and proline residues. The colored sigmoidal curve is included to guide the eye across the transition region, suggesting that – on average – the midpoint of this transition is at a fraction of charged and proline residues of ∼0.25. The Pearson correlation coefficient (r) for the fraction of charged and proline residues vs. normalized radius of gyration is 0.58).

Prior work has implicated the presence of charged and proline residues as mediating IDR chain expansion ^33,34,42,46,49,57,68,87–90^. We took advantage of the fact that the AFRC enables a length normalization of experimental radii of gyration and assessed the normalized radius of gyration vs. the fraction of charged and proline residue (**Fig. 9B**). Our data support this conclusion as a first approximation, but also clearly demonstrate that while this trend is true on average, there is variance in this relationship. Notably, for IDRs with a fraction of charged and proline residues between 0.2 and 0.4, the full range of possible IDR dimensions are accessible. The transition from (on average) more compact to (on average) more expanded chains occurs around a fraction of proline and charged residues of around 0.25 – 0.30, in qualitative agreement with prior work exploring the fraction of charge residues required to drive chain expansion ^33,34,42^. However, we emphasize that there is massive variability observed on a per-sequence basis. In summary, while the presence of charged and proline residues clearly influences IDR dimensions, complex patterns of intramolecular interactions can further tune this behavior ^2,17,28^.

In summary, the AFRC offers a convenient, analytical approach to obtain a well-defined reference state for comparing and contrasting simulations and experiments of unfolded and disordered proteins. It can be easily integrated into complex analysis pipelines, or used for one-off analysis via a Google Colab notebook without requiring any computational expertise at all.

## ACKNOWLEDGEMENTS

We thank members of the Pappu lab and Holehouse lab for many useful discussions over the years. We are indebted to Dr. Nick Lyle for the original implementation of the CAMPARI-based FRC engine. We thank Dr. Erik Martin for bringing the work of Lhuillier to our attention. Funding for this work was provided by the National Institute on Allergic and Infectious Diseases with R01AI163142 to A.S.H. and A.S., by the National Science Foundation with 2128068 to A.S.H., by the Longer Life Foundation, an RGA/Washington University in St. Louis Collaboration to A.S.H., and by the National Cancer Institute with F99CA264413 to J.J.A. We also thank members of the Water and Life Interface Institute (WALII), supported by NSF DBI grant #2213983, for helpful discussions.

## Notes

https://afrc.readthedocs.io/

### Competing Interest Statement

The authors have declared no competing interest.

### Summary of Updates

Fixed author middle initials....

