## Supplementary Information for "The analytical Flory random coil is a simple-to-use reference model for unfolded and disordered proteins"

### 1. SUPPLEMENTARY METHODS

##### *1.1 Flory Random Coil (FRC) simulations, excluded volume (EV) simulations and quantification of finite size effects*

Flory Random Coil (FRC) Monte Carlo simulations were run using a customized version of CAMPARI (V1). Simulations were run in a simulation droplet with a radius of 500 Å for 25 x $10^6$ steps with 50 x $10^3$ steps discarded as equilibration. Conformers were saved every 5 x $10^3$ steps, generating 5 x $10^3$ independent conformations. Because FRC simulations are rejection free, these ensembles are sufficiently well-sampled and enable calibration for FRC fitting parameters (**Table S1**).

Homopolymeric FRC simulations were run for length of 51, 101, 151, 251 and 351 residues for all twenty amino acids (*i.e.* 100 independent sequences in total). Heteropolymeric simulations were run for lengths 10, 20, 30, 40, 50, 100, 120, 140, 180, 200, 250, 300, 350, 400, 450, 500 (*i.e.* 320 independent sequences in total). For each length series, twenty separate simulations were run where, for each sequence, one of the twenty amino acids is enriched (30% of the sequence) while the remaining residues are randomly selected. All FRC simulations were analyzed using SOURSOP[1].

Excluded volume (EV) simulations were run using CAMPARI (V2). In EV simulations, the underlying energy function for the ABSINTH forcefield is altered such that solvation, attractive Lennard-Jones, and polar (charge) interactions are set to zero, as has been described previously[2]. EV simulations were used solely to compare finite-size effects for ensembles constructed for real chains.  Excluded volume (EV) Monte Carlo simulations were run for homopolymers of 50, 100, 150, 200, 250, 300, 350, 400, 450, and 500 residue poly-alanine chains as a reference model to quantify finite-size effects. Simulations were run in a simulation droplet with a radius of 500 Å for 21 x $10^6$ steps, with 1 x $10^6$ steps discarded as equilibration. It is worth noting that given chains are generated in a random non-overlapping starting configuration and the only criterion for move acceptance or rejection is steric overlap, strictly speaking, no equilibration is needed as the chain begins the simulation “equilibrated” in the context of the underlying Hamiltonian. Conformers were saved every 2 x $10^4$ steps, generating 1 x $10^3$ independent conformations, a sufficiently large ensemble for our purposes of calculating internal scaling profiles, although we suggest these ensembles would not be large enough for other types of quantification (**Fig. 1E**).

For quantifying dangle end effects of internal vs. external inter-residue distances (**Fig. S1D**), we ran extensive additional simulations of an $A_{151}$ homopolymer (to match FRC simulations). For these simulations, ten independent replicas were run for 8.05 x $10^7$ steps, with the first 5 x $10^5$ discarded as equilibration. Conformers were saved every 2 x $10^4$ steps. These simulations generated an ensemble of 4 x $10^4$ conformations, enabling a robust assessment of finite-size effects.

We assessed finite-size effects for FRC simulations in several ways, comparing against excluded volume (EV) simulations as a real-chain reference model. First, we compared internal scaling profiles. For real chains, residues at or near the ends have a great volume of space they can explore than residues internal to chain due to excluded volume of the chain. This manifests for internal scaling profiles whereby super-imposing a series of homopolymers of different lengths reveals the distance between residue 1 and n when 1 and *n* are the first and terminal residues is shorter than residue 1 and n when n is an internal residue (**Fig. 1E**). In contrast, because FRC simulations lack any excluded volume contribution, there is no difference between internal and external residues, such that all inter-residue distances of the same residue spacing are equivalent, regardless of where in the chain the two residues lie. This is even more clearly shown by calculating the normalized distance for different inter-residue spacing as a function of starting residue (**Fig. S2C, D**).

Second, we calculated the Flory characteristic ratio as;

$$C_n = \frac{\langle R^2 \rangle}{n l^2} \tag{1}$$

Where $n$ is the number of residues, $l$ is the monomer size, and $\langle R^2 \rangle$ is the ensemble-average squared end-to-end distance (or inter-residue distance)[3]. Given both the FRC and AFRC models describe ideal chains, we can empirically define $l$ as using the standard ideal chain relationship;

$$l = \sqrt{\frac{\langle R^2 \rangle}{n}} \tag{2}$$

in the limit of $n$ tending to $\infty$[3].

By defining $l$ empirically from our FRC simulations or AFRC model, finite size effects emerge upon plotting $n$ vs. $C_n$ (**Fig. S2E,F**). In FRC simulations, $C_n$ is less than 1 for shorter chains. This is expected in that the rotational isomeric state means local chain geometry is not truly ideal but instead limited to the inter-residue vector path defined by the Ramachandran isomeric states. In contrast, the AFRC is a true ideal chain model, such that the Flory characteristic ratio is always 1 regardless of $n$. This difference between the AFRC and FRC models manifests as a very slight (1-2 Å) difference in intramolecular distances visible in **Fig. 2A**.

##### *1.2 All-atom simulations*

All-atom simulations were analyzed as described previously, and all the all-atom trajectories can be obtained as described previously[1]. Specifically, all-atom simulations included both Monte Carlo and molecular dynamics simulations. Monte Carlo simulations include those of Ash1[4], p53[5], p27[6], the notch intracellular domain[7], the hnRNPA1 low complexity domain[8]. Molecular dynamics simulations include alpha-synuclein, DrkN, ACTR and NTail [9].

##### *1.3 SAXS data*

Experimental SAXS data includes 145 separate radius of gyration values. All values and associated references are included in table S4. In addition, all data are tabulated at the main GitHub directory for this paper (https://github.com/holehouse-lab/supportingdata/tree/master/2023/alston_ginell_2023) and available as an Excel spreadsheet and Pandas-compatible CSV file.

##### *1.4 Amino acid sequence analysis*

Sequence analysis to calculate the fraction of charged residues and proline residues was done using localCIDER [10] and sparrow (https://github.com/idptools/sparrow).

##### *1.5 AFRC implementation*

The AFRC is implemented as a stand-alone Python package. All code is open-sourced and available at https://github.com/idptools/afrc. All documentation is available at https://afrc.readthedocs.io/. The package itself can be downloaded from https://pypi.org/project/afrc and installed using the command `pip install afrc`. A Google colab notebook that implements the AFRC along with the other three analytical models described in this work are linked from https://github.com/idptools/afrc.

The afrc package uses numpy and scipy, and in addition to the AFRC implements the Worm-like chain (WLC), the self-avoiding random walk (SAW), and the v-dependent self-avoiding random walk (SAW-v) [11,12].

##### *1.6 Figures and analysis in this paper*

Jupyter notebooks to recreate all figures in this paper are available at https://github.com/holehouse-lab/supportingdata/tree/master/2023/alston_ginell_2023.

#### 2. SUPPLEMENTARY FIGURES

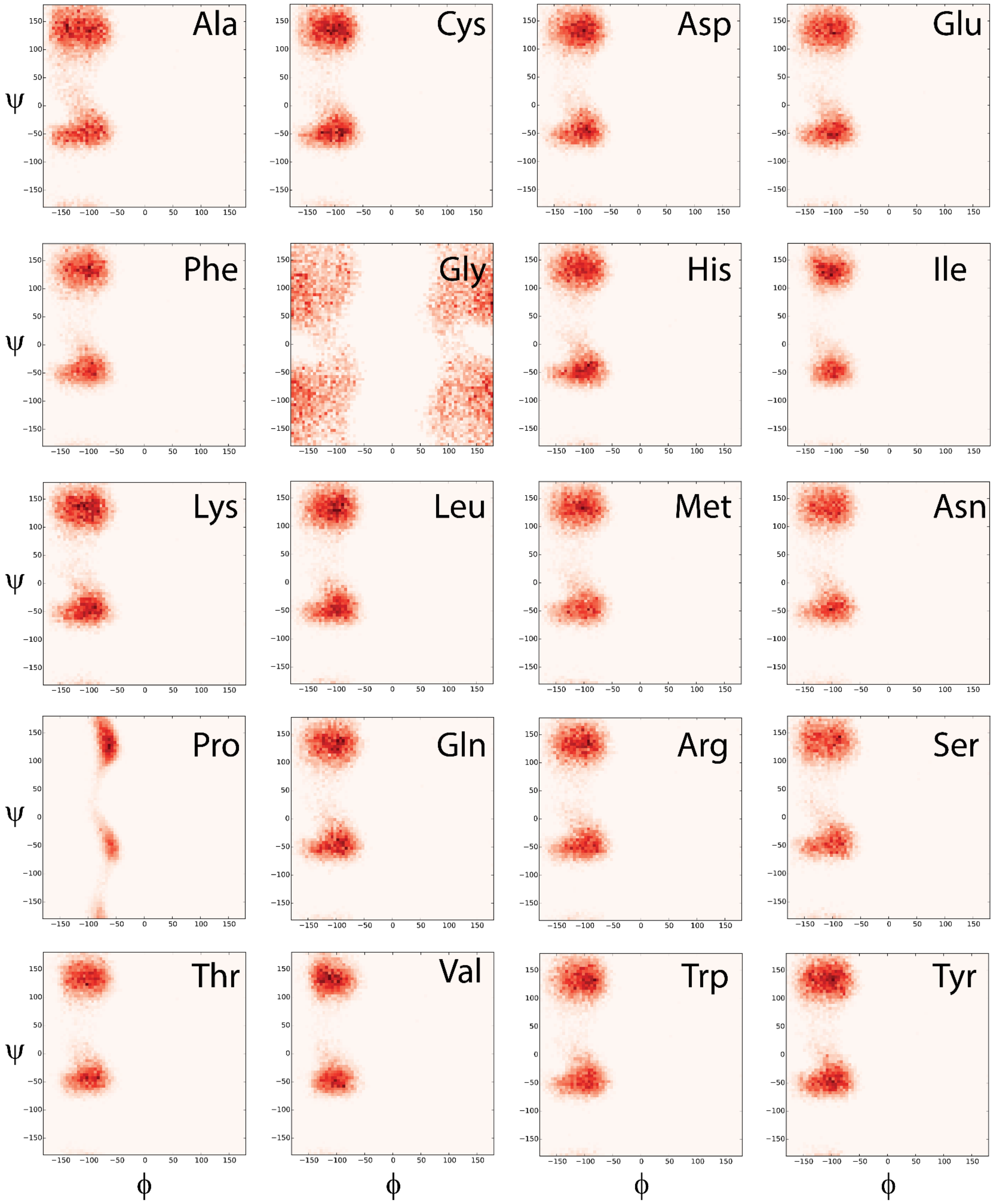


**Fig. S1 Residue-specific Ramachandran maps used for FRC simulations.** Ramachandran maps for all twenty amino acids performed as excluded volume simulations define the allowed isomeric states and are used by FRC simulations to construct the FRC ensembles.

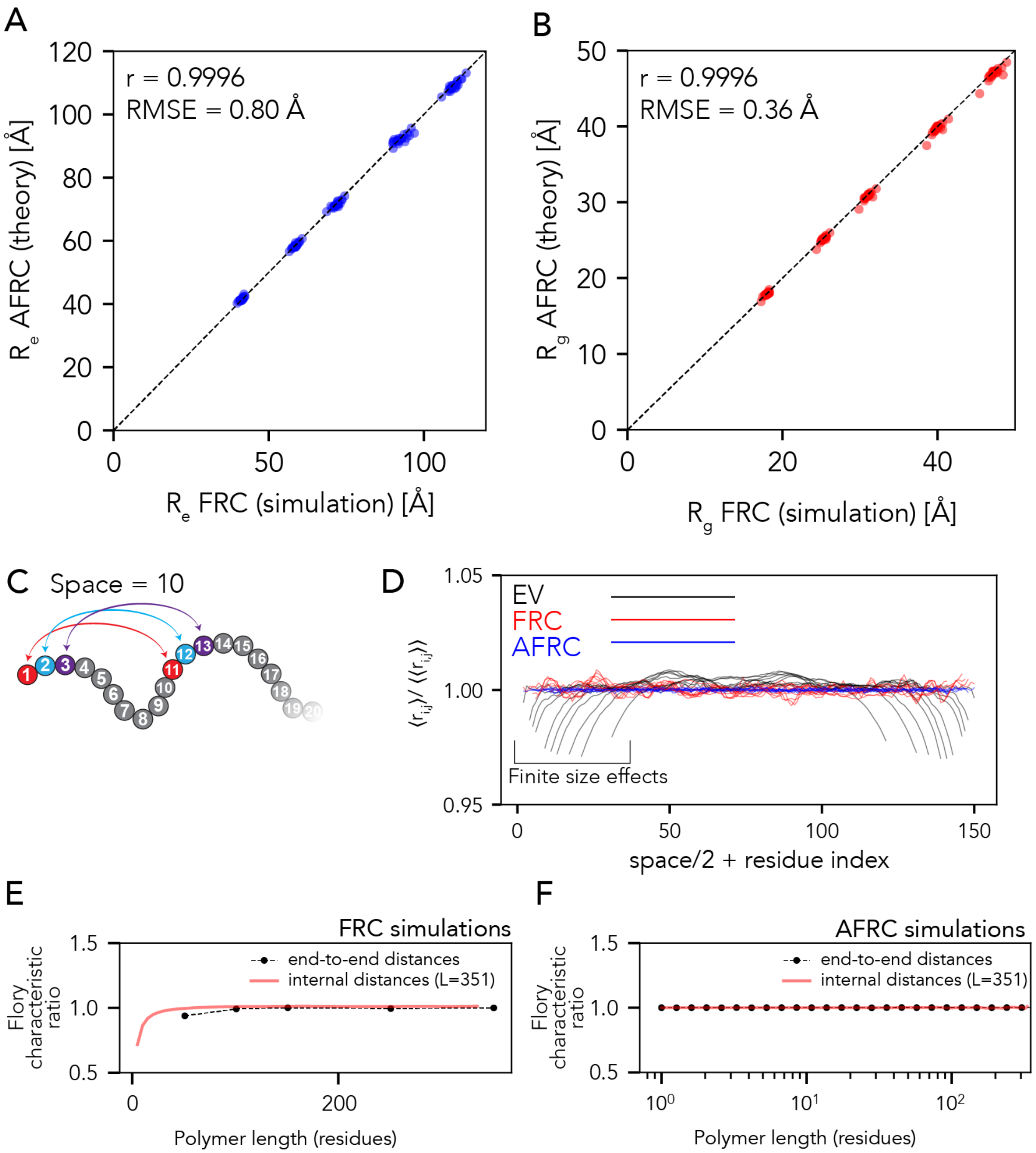


**Fig. S2 Comparison between global dimensions from simulations vs. AFRC. A.** The correlation between the end-to-end distance ($R_e$) obtained from FRC simulations and AFRC analysis is shown. The comparisons here are for ensemble-average values for homopolymers derived from the twenty different amino acids for lengths of 51, 101, 151, 251, and 351 residues.

**B.** The correlation between radius of gyration ($R_g$) values obtained from FRC simulations and AFRC analysis. Again, the comparisons here are for ensemble-average values for homopolymers derived from the twenty different amino acids for lengths of 51, 101, 151, 251 and 351 residues. **C.** Schematic of the approach taken in panel D. **D.** For a 151-residue homopolymer, we calculated the average distance between all pairs of residues that are a fixed spacing apart for EV and FRC simulations and for the AFRC model. The inter-residue spacing used were 2, 6, 8, 10, 16, 20, 24, 32, 40, and 60 residues, and each spacing yields a different line. For example, for a spacing of 6 residues, we calculated the average distance between the following pairs of residues $\langle r_{1,7} \rangle$, $\langle r_{2,8} \rangle$, ..., $\langle r_{145,151} \rangle$. Note the angle brackets here denote the ensemble-average distance. Each line represents the profile revealed by the set of inter-residue distances. For every point along the line, the y-axis position reports on the average distance normalized by the overall average distance for all residues of a given spacing. In contrast, the x-axis position is the location of the first residue of the two in a pair, to which half of the inter-residue spacing is added. For example, if we examined positions for $\langle r_{1,7} \rangle$, $\langle r_{2,8} \rangle$, ..., $\langle r_{145,151} \rangle$ then the corresponding x-axis positions would be ($1 + 0.5\times6 = 4$, $2 + 0.5\times6 = 5$, …, $145 + 0.5{*}\times6 = 148$). We take this approach such that the middle of the x-axis in the figure always corresponds to the central position in the polymer. For EV simulations, when one of the two residues in a pair falls near the end of the chain, we see a suppression of the inter-residue distances compared to the same inter-residue distance when both positions are internal to the chain. This is the expected result and reflects the fact that internal residues are 'repelled' by steric overlap with other residues, whereas end residues are less constrained. For FRC simulations and AFRC models, no such end effects are observed, reflecting the finite-size end effects do not influence ideal chains. **E**. We also calculated the Flory characteristic ratio ($C_n$) for chains of different lengths (black circles) and for intramolecular distances (red lines) for FRC simulations. The characteristic ratio enables correlations in chain dimensions to be assessed, and for FRC simulations, we see the expected deviation from 1 at shorter chain lengths (see supplemental methods). While these deviations are expected finite-size effects, their impact when comparing inter-residue distances is minimal (**Fig. 2**).

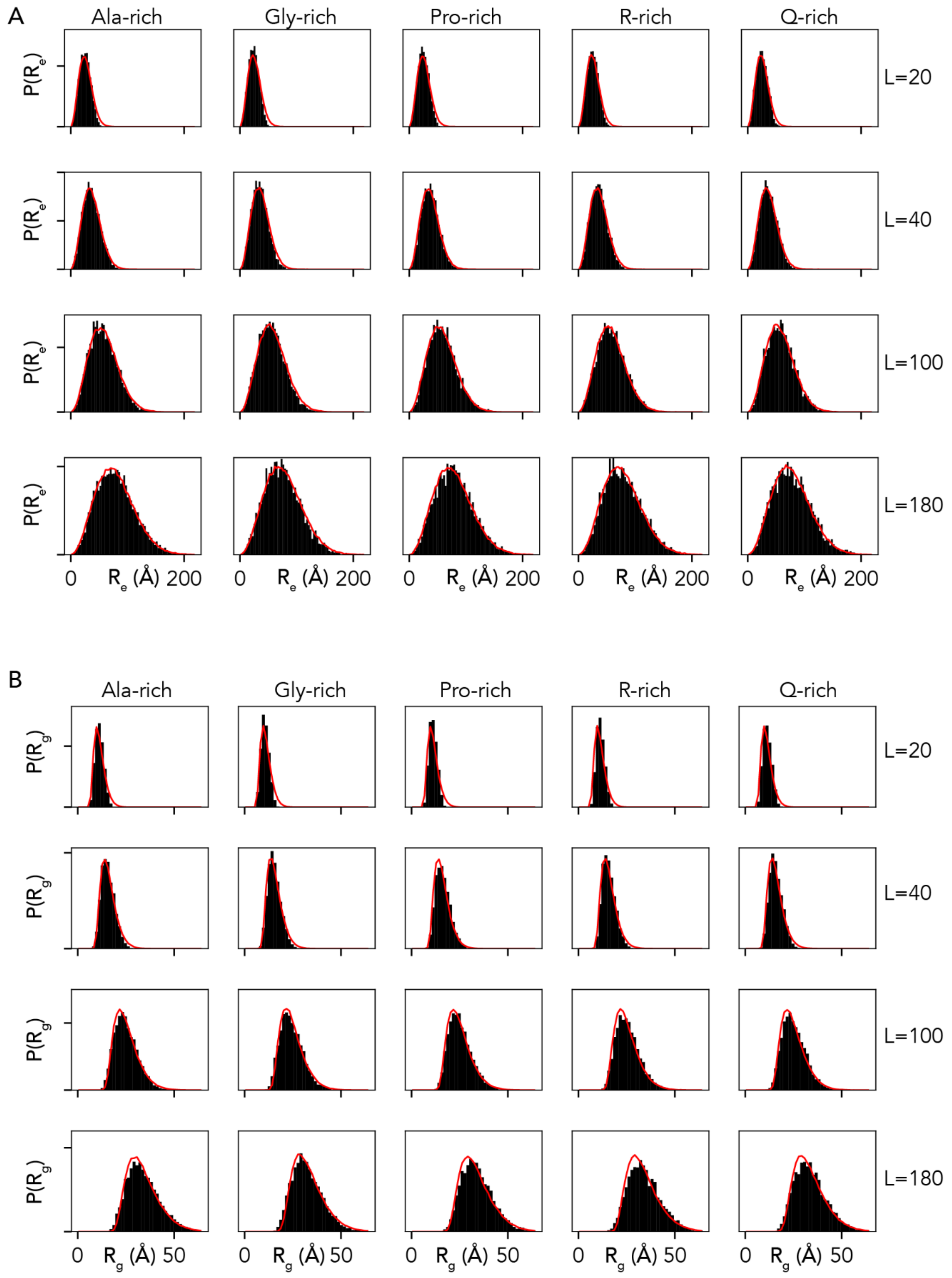


**Fig. S3 Comparison of end-to-end distance distributions and radii of gyration distributions for select heteropolymers of variable composition and length. A.** Comparison of end-to-end distance distributions. Empirical distributions obtained from simulations are shown in black, while predictions of the distribution from the AFRC are shown as red lines. **B.** Comparison of radii of gyration distributions. Empirical distributions obtained from simulations shown in black, while predictions of the distribution from the AFRC are shown as red lines.

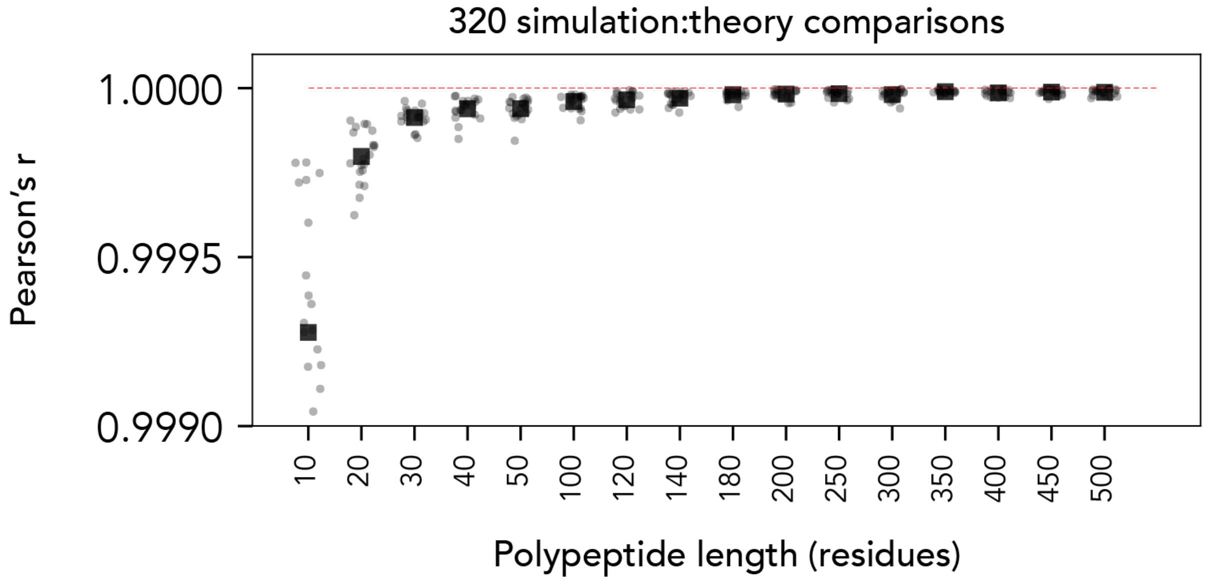


**Fig. S4. Correlation between internal scaling profiles for random heteropolymers from FRC simulations vs. AFRC-derived internal scaling profiles.** For each length (10,20,30, …, 500) 20 different heteropolymers, were generated where each heteropolymer is enriched (30%) in one of the twenty amino acids while the remaining residues are randomly selected. This yields 320 different internal scaling comparisons (16 lengths with 20 amino acids).

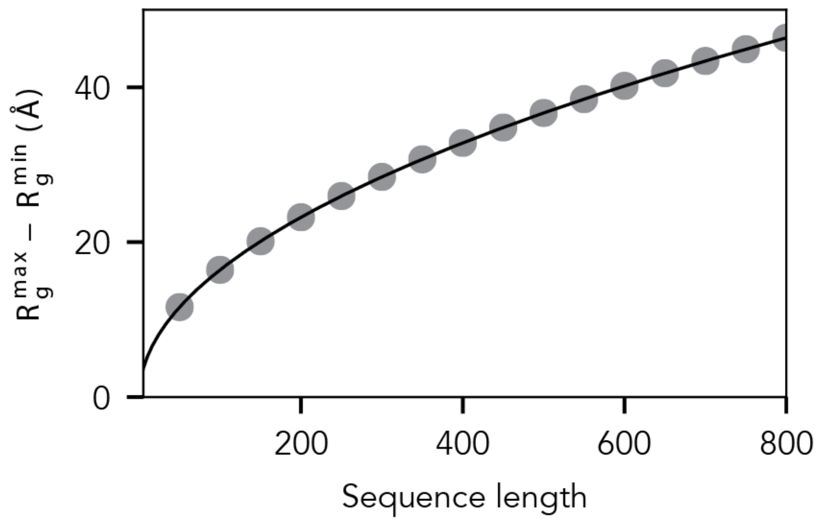


**Fig. S5.** Difference in radii of gyration based on empirical min and max values reveals the length-dependent variation in expected accessible radii of gyration values.

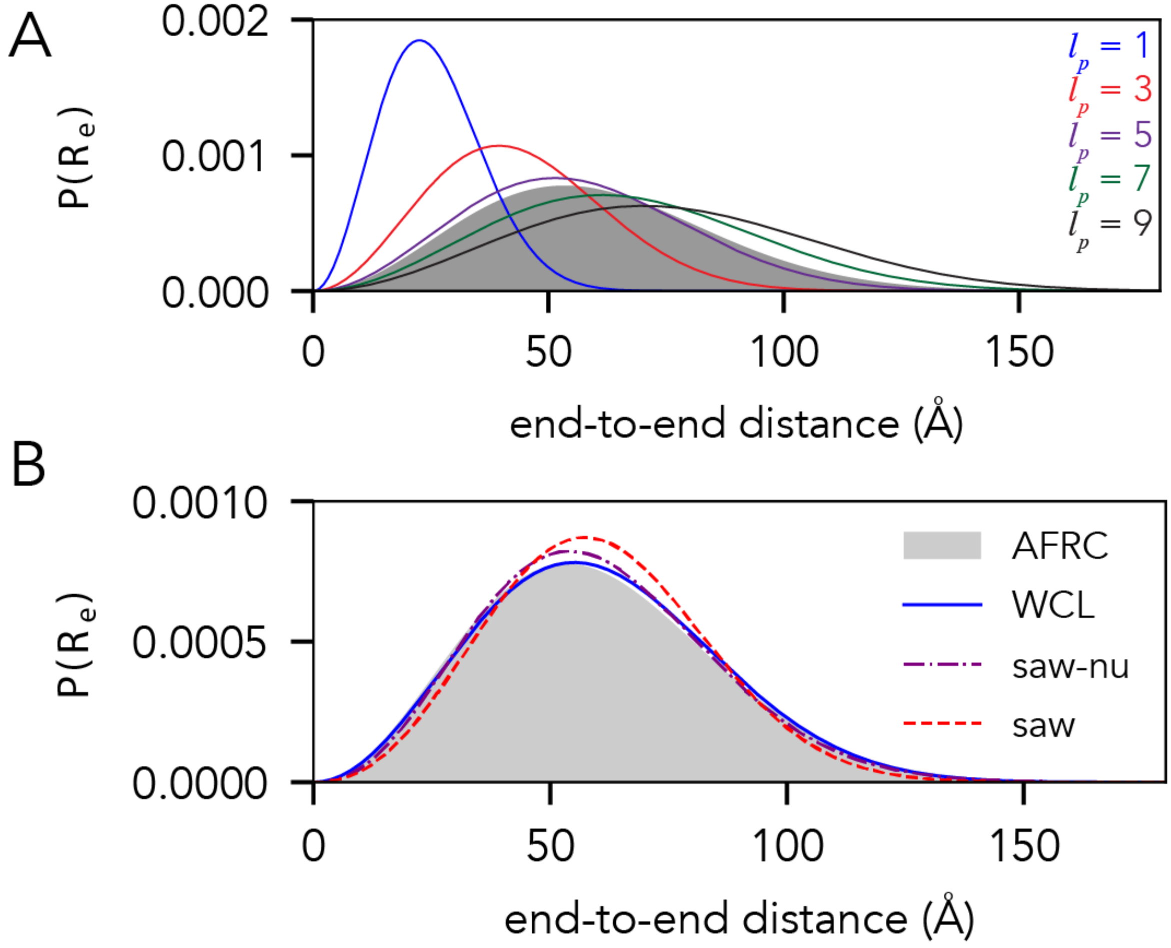


**Fig. S6.** Comparison of the end-to-end distance distributions for the AFRC with existing polymer models. **A.** Comparison of the AFRC model (grey shaded area) for 100-residue polyalanine chain ($A_{100}$) with Worm-Like chain (WLC)-derived distributions, where the WLC monomer size is fixed at 3.8 Å, and the persistence length varies from 1 Å to 9 Å. **B.** Comparison of AFRC, WLC, SAW-ν, and SAW models in which model input parameters were selected to reproduce the AFRC end-to-end distance distribution for an $A_{100}$ chain. The WLC model uses an amino acid size of 3.8 Å and a persistence length of 5.7 Å. The SAW-ν model uses a prefactor of 5.8 Å and a ν of 0.5. The SAW model uses a prefactor of 4.1 Å.

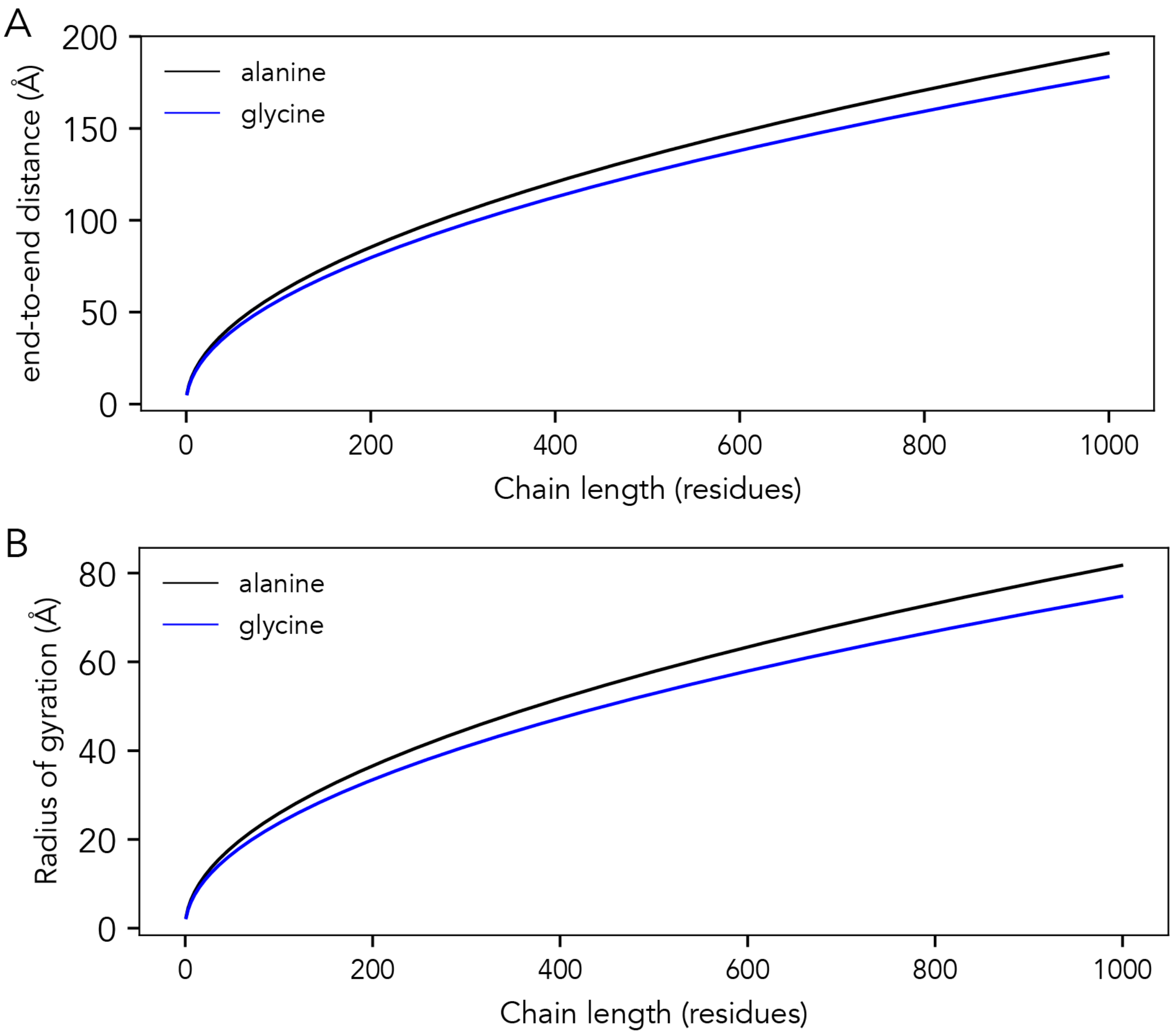


**Fig. S7.** Comparison of chain dimensions obtained from the AFRC model for poly-alanine vs. poly-glycine, examining end-to-end distance (**A**) and radius of gyration (**B**).

#### 3. SUPPLEMENTARY TABLES

| **Amino acid** | **$R_{ij}$ RMS (Å)** | **$R_{ij}$ (Å)** | **$X_0$ (Å$^{-1}$)** |
|---|---|---|---|
| A | 6.5463 | 6.0381 | 0.5405 |
| C | 6.2676 | 5.7826 | 0.5635 |
| D | 6.3994 | 5.911 | 0.5567 |
| E | 6.2649 | 5.768 | 0.5613 |
| F | 6.2519 | 5.7612 | 0.5571 |
| G | 6.1045 | 5.6324 | 0.5911 |

| | | | |
|---|---|---|---|
| H | 6.2156 | 5.7262 | 0.5645 |
| I | 6.4353 | 5.9361 | 0.5483 |
| K | 6.306 | 5.8272 | 0.5533 |
| L | 6.2636 | 5.7801 | 0.5605 |
| M | 6.3813 | 5.8894 | 0.5501 |
| N | 6.2652 | 5.773 | 0.5598 |
| P | 6.4323 | 5.9388 | 0.5599 |
| Q | 6.2547 | 5.7719 | 0.5617 |
| R | 6.279 | 5.7921 | 0.5531 |
| S | 6.3161 | 5.8364 | 0.5553 |
| T | 6.1995 | 5.7242 | 0.5695 |
| V | 6.3204 | 5.8409 | 0.5571 |
| W | 6.3 | 5.814 | 0.5539 |
| Y | 6.3188 | 5.8266 | 0.5543 |

**Table S1** Model parameters obtained by fitting against FRC simulations.

| Name | Sequence |
|---|---|
| **Ash1** | GASASSSPSP STPTKSGKMR SRSSSPVRPK AYTPSPRSPN YHRFALDSPP QSPRRSSNSS ITKKGSRRSS GSSPTRHTTR VCV |
| **p53** | MEEPQSDPSV EPPLSQETFS DLWKLLPENN VLSPLPSQAM DDLMLSPDDI EQWFTEDPGP DEAPRMPEAA PPVAPAPAAP TPAAPAPAPS W |
| **p27** | GSHMKGACKV PAQESQDVSG SRPAAPLIGA PANSEDTHLV DPKTDPSDSQ TGLAEQCAGI RKRPATDDSS TQNKRANRTE ENVSDGSPNA GSVEQTPKKP GLRRRQT |
| **Notch** | MARKRRRQHG QLWFPEGFKV SEASKKKRRE PLGEDSVGLK PLKNASDGAL MDDNQNEWGD EDLETKKFRF EEPVVLPDLD DQTDHRQWTQ QHLDAADLRM SAMAPTPPQG EVDADCMDVN VRGPDGFTPL LE |
| **ACTR** | GTQNRPLLRN SLDDLVGPPS NLEGQSDERA LLDQLHTLLS NTDATGLEEI DRALGIPELV NQGQALEPKQ D |
| **drkN** | MEAIAKHDFS ATADDELSFR KTQILKILNM EDDSNWYRAE LDGKEGLIPS NYIEMKNHD |
| **Ntail** | MHHHHHHTTE DKISRAVGPR QAQVSFLHGD QSENELPRLG GKEDRRVKQS RGEARESYRE TGPSRASDAR AAHLPTGTPL DIDTASESSQ DPQDSRRSAD ALLRLQAMAG ISEEQGSDTD TPIVYNDRNL LD |
| **asyn** | MDVFMKGLSK AKEGVVAAAE KTKQGVAEAA GKTKEGVLYV GSKTKEGVVH GVATVAEKTK EQVTNVGGAV VTGVTAVAQK TVEGAGSIAA ATGFVKKDQL GKNEEGAPQE GILEDMPVDP DNEAYEMPSE EGYQDYEPEA |
| **A1-LCD** | GSMASASSSQ RGRSGSGNFG GGRGGGFGGN DNFGRGGNFS GRGGFGGSRG GGGYGGSGDG YNGFGNDGSN FGGGGSYNDF GNYNNQSSNF GPMKGGNFGG RSSGPYGGGG QYFAKPRNQG GYGGSSSSSS YGSGRRF |

**Table S2. Sequences from simulations**. Full sequences used from all-atom simulations. Amino acids are colored by chemical type as per localCIDER[10].

| Name | N | $R_g$ (Å) | $R_g/R_g^{\theta}$ | $R_e$ (Å) | $R_e/R_e^{\theta}$ | $\nu^{app}$ [(a)] | Quality of $\nu^{app}$ fit [(b)] |
|---|---|---|---|---|---|---|---|
| **Ash1** | 83 | 28.9 | 1.27 | 68.95 | 1.30 | 0.61 | GOOD |
| **p53** | 91 | 29.4 | 1.23 | 77.73 | 1.39 | 0.66 | GOOD |
| **p27** | 107 | 28.3 | 1.09 | 59.15 | 0.98 | 0.49 | POOR |
| **Notch** | 132 | 29.3 | 1.02 | 52.16 | 0.78 | 0.34 | POOR |
| **ACTR** | 71 | 21.1 | 1.01 | 41.45 | 0.85 | 0.50 | GOOD |
| **drkN** | 59 | 19.3 | 1.00 | 45.26 | 1.01 | 0.43 | GOOD |
| **Ntail** | 132 | 26.3 | 0.92 | 58.11 | 0.87 | 0.39 | POOR |
| **asyn** | 140 | 25.6 | 0.87 | 46.47 | 0.67 | 0.23 | POOR |
| **A1-LCD** | 137 | 24.1 | 0.84 | 54.37 | 0.81 | 0.47 | GOOD |

[a] Estimated $\nu^{app}$ based on linear fitting of the internal scaling regime using SOURSOP.

[b] Quality of fit based on the reduced chi-squared from the fit.

**Table S3:** Simulation and AFRC-derived parameters for all-atom simulations.

**Table S4**: SAXS sequences and values (note table caption comes before table as table is 36 pages long).

| Protein name | $R_g$ (Å) | $R_g$ error (Å) | Amino acid sequence | Reference |
|---|---|---|---|---|
| Nucleoporin Nup49 (N49) | 15.9 | 1.3 | GCQTSRGLFGNNNTNNINNSSSGMNNASAGLFGSKPCA | Fuertes, et al. PNAS (2017) 114, E6342–E6351. |
| Heh2 (NLS) | 24 | 3 | ACETNKRKREQISTDNEAKMQIQEEKSPKKKRKKRSSKANKPPECA | Fuertes, et al. PNAS (2017) 114, E6342–E6351. |
| VSV Protein Phosphoprotein P | 24 | 1 | HHHHHELMDNLTKVREYLKSYSRLDQAVGEIDEIEAQRAEKSNYELFQEDGVEEHTKPSYFQAADDS | Leyrat, C., Jensen, M.R., Ribeiro, E.A., Gérard, F.C.A., Ruigrok, R.W.H., Blackledge, M., and Jamin, M. (2011). The N0-binding region of the vesicular stomatitis virus phosphoprotein is globally disordered but contains transient α-helices. Protein Sci. 20, 542–556. |
| LS | 27.9 | 1 | SPPGKPQGPPQQEGNKPQGPPPPGKPQGPPPAGGNPQQPQAPPAGKPQGPPPPPQGGRPPRPAQGQQPPQ | Boze, H., Marlin, T., Durand, D., Pérez, J., Vernhet, A., Canon, F., Sarni-Manchado, P., Cheynier, V., and Cabane, B. (2010). Proline-rich salivary proteins have extended conformations. Biophys. J. 99, 656–665. |
| Nup153_NUS | 24.9 | 1.3 | GCPSASPAFGANQTPTFGQSQGASQPNPPGFGSISSSTALFPTGSQPAPPTFGTVSSSSQPPVFGQQPSQSAFGSGTTPNCA | Fuertes, et al. PNAS (2017) 114, E6342–E6351. |

| | | | | |
|---|---|---|---|---|
| Sic1 | 30 | 4 | GSMTPSTPPRSRGTRYLAQPSGNTSSSALMQGQKTPQKPSQNLVPVTPSTTKSFKNAPLLAPPNSNMGMTSPFNGLTSPQRSPFPKSSVKRT | Gomes G-NW, Krzeminski M, Namini A, Martin EW, Mittag T, Head-Gordon T, et al. Conformational Ensembles of an Intrinsically Disordered Protein Consistent with NMR, SAXS, and Single-Molecule FRET. J Am Chem Soc. 2020;142: 15697–15710. |
| chloroplastic calvin cycle protein | 23 | | HHHHHHHHHHSSGHIEGRHMSGQPAVDLNKKVQDAVKEAEDACAKGTSADCAVAWDTVEELSAAVSHKKDAVKADVTLTDPLEAFCKDAPDADECRVYED | Launay H, Barré P, Puppo C, Zhang Y, Maneville S, Gontero B, Receveur-Bréchot V, J Mol Biol 430(8):1218-1234 (2018) |
| Antitermination protein N (from lambda phage) | 38 | 3.5 | MDAQTRRRERRAEKQAQWKAANPLLVGVSAKPVNRPILSLNRKPKSRVESALNPIDLTVLAEYHKQIESNLQRIERKNQRTWYSKPGERGITCSGRQKIKGKSIPLI | Johansen, D., Trewhella, J., and Goldenberg, D.P. (2011). Fractal dimension of an intrinsically disordered protein: small-angle X-ray scattering and computational study of the bacteriophage λ N protein. Protein Sci. 20, 1955–1970. |
| Nup153_NUL | 30 | 3 | GCGFKGFDTSSSSSNSAASSSFKFGVSSSSSGPSQTLTSTGNFKFGDQGGFKIGVSSDSGSINPMSEGFKFSKPIGDFKFGVSSESKPEEVKKDSKNDNFKFGLSSGLSNPVCA | Fuertes, et al. PNAS (2017) 114, E6342–E6351. |
| DARPP-32 (aka Protein phosphatase 1 regulatory subunit 1B) | 28.28 | | MDPKDRKKIQFSVPAPPSQLDPRQVEMIRRRRPTPALLFRVSEHSSPEEESSPHQRTSGEGHHPKSKRPNPCAYTPPSLKAVQRIAESHLQTISNLSENQASEEEDELGELRELGYPQ | Marsh, J.A., Dancheck, B., Ragusa, M.J., Allaire, M., Forman-Kay, J.D., and Peti, W. (2010). Structural diversity in free and bound states of intrinsically disordered protein phosphatase 1 regulators. Structure 18, 1094–1103. |

| | | | | |
|---|---|---|---|---|
| II-1 | 41 | | GKPVGRRPQGGNQPQRPPPPPGKPQGPPPQGGNQSQGPPPPPGKPEGRPPQGRNQSQGPPPHPGKPERPPPQGGNQSQGTPPPPGKPERPPPQGGNQSHRPPPPPGKPERPPPQGGNQSRGPPPHRGKPEGPPPQEGNKSR | Boze, H., Marlin, T., Durand, D., Pérez, J., Vernhet, A., Canon, F., Sarni-Manchado, P., Cheynier, V., and Cabane, B. (2010). Proline-rich salivary proteins have extended conformations. Biophys. J. 99, 656–665. |
| Fhua | 33.4 | | ESAWGPAATIAARQSATGTKTDTPIQKVPQSISVVTAEEMALHQPKSVKEALSYTPGVSVGTRGASNTYDHLIIRGFAAEGQSQNNYLNGLKLQGNFYNDAVIDPYMLERAEIMRGPVSVLYGKSSPGGLLNMVSKRPTTEP | Riback, J.A., Bowman, M.A., Zmyslowski, A.M., Knoverek, C.R., Jumper, J.M., Hinshaw, J.R., Kaye, E.B., Freed, K.F., Clark, P.L., and Sosnick, T.R. (2017). Innovative scattering analysis shows that hydrophobic disordered proteins are expanded in water. Science 358, 238–241. |
| N98 | 28.6 | 1.3 | GCFNKSFGTPFGGGTGGFGTTSTFGQNTGFGTTSGGAFGTSAFGSSNNTGGLFGNSQTKPGGLFGTSSFSQPATSTSTGFGFGTSTGTANTLFGTASTGTSLFSSQNNAFAQNKPTGFGNFGTSTSSGGLFGTTNTTSNPFGSTSGSLFGPCA | Fuertes, et al. PNAS (2017) 114, E6342–E6351. |
| Protein Phosphatase Inhibitor 2 | 34.6 | | PIKGILKNKTSTTSSMVASAEQPRGNVDEELSKKSQKWDEMNILATYHPADKDYGLMKIDEPSTPYHSMMGDDEDACSDTEATEAMAPDILARKLAAAEGLEPKYRIQEQESSGEEDSDLSPEEREKKRQFEMKRKLHYNEGLNIKLARQLISKDL | Marsh, J.A., Dancheck, B., Ragusa, M.J., Allaire, M., Forman-Kay, J.D., and Peti, W. (2010). Structural diversity in free and bound states of intrinsically disordered protein phosphatase 1 regulators. Structure 18, 1094–1103. |
| Nsp1 | 41 | 3 | GCNFNTPQQNKTPFSFGTANNNSNTTNQNSSTGAGAFGTGQSTFGFNNSAPNNTNNANSSITPAFGSNNTGNTAFGNSNPTSNVFGSNNSTTNTFGSNSAGTSLFGSSSAQQTKSNGTAGGNTFGSSSLFN | Fuertes, et al. PNAS (2017) 114, E6342–E6351. |

| | | | | |
|---|---|---|---|---|
| | | | NSTNSNTTKPAFGGLNFGGGNNTTPSSTGNANTSNNLFGATANANCA | |
| IBB | 32 | 2 | GCTNENANTPAARLHRFKNKGKDSTEMRRRRIEVNVELRKAKKDDQMLKRRNVSSFPDDATSPLQENRNNQGTVNWSVDDIVKGINSSNVENQLQATCA | Fuertes, et al. PNAS (2017) 114, E6342–E6351. |
| Ash1 | 28.5 | 3.4 | GASASSSPSPSTPTKSGKMRSRSSSPVRPKAYTPSPRSPNYHRFALDSPPQSPRRSSNSSITKKGSRRSSGSSPTRHTTRVCV | Martin, E.W., Holehouse, A.S., Grace, C.R., Hughes, A., Pappu, R.V., and Mittag, T. (2016). Sequence Determinants of the Conformational Properties of an Intrinsically Disordered Protein Prior to and upon Multisite Phosphorylation. J. Am. Chem. Soc. 138, 15323–15335. |
| pAsh1 | 27.5 | 1.2 | GASASSSPSPSTPTKSGKMRSRSSSPVRPKAYTPSPRSPNYHRFALDSPPQSPRRSSNSSITKKGSRRSSGSSPTRHTTRVCV | Martin, E.W., Holehouse, A.S., Grace, C.R., Hughes, A., Pappu, R.V., and Mittag, T. (2016). Sequence Determinants of the Conformational Properties of an Intrinsically Disordered Protein Prior to and upon Multisite Phosphorylation. J. Am. Chem. Soc. 138, 15323–15335. |
| PIR domain (GRB14) | 27 | | YGMQLYQNYMHPYQGRSGCSSQSISPMRSISENSLVAMDFSGQKSRVIENPTEALSVAVEEGLAWRKKGCLRLGTHGSPTASSQSSATNMAIHRSQPW | Moncoq, K., Broutin, I., Craescu, C.T., Vachette, P., Ducruix, A., and Durand, D. (2004). SAXS study of the PIR domain from the Grb14 molecular adaptor: a natively unfolded protein with a transient structure primer? Biophys. J. 87, 4056–4064. |

| | | | | |
|---|---|---|---|---|
| RpII215_gibbs | 28 | 0.7 | YSPGNAYSPSSSNYSPNSPSYSPTSPSYSPSSPSYSPTSPCYSPTSPSYSPTSPNYTPVTPSYSPTSPNYSASPQ | Gibbs, E.B., Lu, F., Portz, B., Fisher, M.J., Medellin, B.P., Laremore, T.N., Zhang, Y.J., Gilmour, D.S., and Showalter, S.A. (2017). Phosphorylation induces sequence-specific conformational switches in the RNA polymerase II C-terminal domain. Nat. Commun. 8, 15233. |
| RpII215_portz | 51.8 | | SPSYSPTSPNYTASSPGGASPNYSPSSPNYSPTSPLYASPRYASTTPNFNPQSTGYSPSSSGYSPTSPVYSPTVQFQSSPSFAGSGSNIYSPGNAYSPSSSNYSPNSPSYSPTSPSYSPSSPSYSPTSPCYSPTSPSYSPTSPNYTPVTPSYSPTSPNYSASPQYSPASPAYSQTGVKYSPTSPTYSPPSPSYDGSPGSPQYTPGSPQYSPASPKYSPTSPLYSPSSPQHSPSNQYSPTGSTYSATSPRYSPNMSIYSPSSTKYSPTSPTYTPTARNYSPTSPMYSPTAPSHYSPTSPAYSPSSPTFEESED | Portz, B., Lu, F., Gibbs, E.B., Mayfield, J.E., Rachel Mehaffey, M., Zhang, Y.J., Brodbelt, J.S., Showalter, S.A., and Gilmour, D.S. (2017). Structural heterogeneity in the intrinsically disordered RNA polymerase II C-terminal domain. Nat. Commun. 8, 15231. |
| ACTR | 25 | | GPSGTQNRPLLRNSLDDLVGPPSNLEGQSDERALLDQLHTLLSNTDATGLEEIDRALGIPELVNQGQALEPKQDSGGPR | Borgia, A., Zheng, W., Buholzer, K., Borgia, M.B., Schüler, A., Hofmann, H., Soranno, A., Nettels, D., Gast, K., Grishaev, A., et al. (2016). Consistent View of Polypeptide Chain Expansion in Chemical Denaturants from Multiple Experimental Methods. J. Am. Chem. Soc. 138, 11714–11726. |
| Msh6 | 56 | 2 | MAPATPKTSKTAHFENGSTSSQKKMKQSSLLSFFSKQVPSGTPSKKVQKPTPATLENTATDKITKNPQGGKTGKLFVDVDEDNDLTIAEETVSTVRSDIMHSQEPQSDTMLNSNTTEPKSTTTDEDLSSSQSRRNHKRRVNYAESDDDDSDTTFTAKRKKGKVVDSESDEDEYLPDKNDGDEDDDIADDKEDIKGELAEDSGDDDDLISLAETTSKKKFSYNTSHSSSPFTRNIS | Shell, S.S., Putnam, C.D., and Kolodner, R.D. (2007). The N terminus of Saccharomyces cerevisiae Msh6 is an unstructured tether to PCNA. Mol. Cell 26, 565–578. |

| | | | | |
|---|---|---|---|---|
| | | | RDNSKKKSRPNQAPSRSYNPSHSQPSATSKSSKFNKQNEERYQWLVDERDAQRRPKSDPEYDPRTLYIP | |
| AN16 | 50 | 2 | AQTPSSQYGAPAQTPSSQYGAPAQTPSSQYGAPAQTPSSQYGAPAQTPSSQYGAPAQTPSSQYGAPAQTPSSQYGAPAQTPSSQYGAPAQTPSSQYGAPAQTPSSQYGAPAQTPSSQYGAPAQTPSSQYGAPAQTPSSQYGAPAQTPSSQYGAPAQTPSSQYGAPAQTPSSQYGAP | Nairn, K.M., Lyons, R.E., Mulder, R.J., Mudie, S.T., Cookson, D.J., Lesieur, E., Kim, M., Lau, D., Scholes, F.H., and Elvin, C.M. (2008). A synthetic resilin is largely unstructured. Biophys. J. 95, 3358–3365. |
| HrpO | 35 | | MEDTLEDDPQRAALEQVISLLTPVRQHRQASAERAHRHAQVELKSMLDHLSKIRASLDQERDNHKRRREGLSQEHLEKTISPNDIDRWHEKEKHMLDRLACIRQDVQQQQLRVAEQQALLEQKRLQAKASQRAVEKLACMEETLNEEG | Gazi, A.D., Bastaki, M., Charova, S.N., Gkougkoulia, E.A., Kapellios, E.A., Panopoulos, N.J., and Kokkinidis, M. (2008). Evidence for a Coiled-coil Interaction Mode of Disordered Proteins from Bacterial Type III Secretion Systems. J. Biol. Chem. 283, 34062–34068. |
| alpha-syn | 41 | 1 | MDVFMKGLSKAKEGVVAAAEKTKQGVAEAAGKTKEGVLYVGSKTKEGVVHGVATVAEKTKEQVTNVGGAVVTGVTAVAQKTVEGAGSIAAATGFVKKDQLGKNEEGAPQEGILEDMPVDPDNEAYEMPSEEGYQDYEPEA | Uversky, V.N., Li, J., Souillac, P., Millett, I.S., Doniach, S., Jakes, R., Goedert, M., and Fink, A.L. (2002). Biophysical properties of the synucleins and their propensities to fibrillate: inhibition of alpha-synuclein assembly by beta- and gamma-synucleins. J. Biol. Chem. 277, 11970–11978. |
| NTail | 27.2 | 0.5 | TTEDKISRAVGPRQAQVSFLHGDQSENELPRLGGKEDRRVKQSRGEARESYRETGPSRASDARAAHLPTGTPLDIDTASESSQDPQDSRRSADALLRLQAMAGISEEQGSDTDTPIVYNDRNLLD | Longhi, S., Receveur-Bréchot, V., Karlin, D., Johansson, K., Darbon, H., Bhella, D., Yeo, R., Finet, S., and Canard, B. (2003). The C-terminal domain of the measles virus nucleoprotein is intrinsically |

| | | | | |
|---|---|---|---|---|
| | | | | disordered and folds upon binding to the C-terminal moiety of the phosphoprotein. J. Biol. Chem. 278, 18638–18648. |
| ERM | 39.6 | 0.7 | MDGFYDQQVPFMVPGKSRSEECRGRPVIDRKRKFLDTDLAHDSEELFQDLSQLQEAWLAEAQVPDDEQFVPDFQSDNLVLHAPPPTKIKRELHSPSSELSSCSHEQALGANYGEKCLYNYCA | Lens, Z., Dewitte, F., Monté, D., Baert, J.-L., Bompard, C., Sénéchal, M., Van Lint, C., de Launoit, Y., Villeret, V., and Verger, A. (2010). Solution structure of the N-terminal transactivation domain of ERM modified by SUMO-1. Biochem. Biophys. Res. Commun. 399, 104–110. |
| Neuroligin-3 | 33 | 3 | YRKDKRRQEPLRQPSPQRGAGAPELGAAPEEELAALQLGPTHHECEAGPPHDTLRLTALPDYTLTLRRSPDDIPLMTPNTITMIPNSLVGLQTLHPYNTFAAGFNSTGLPHSHSTTRV | Paz, A., Zeev-Ben-Mordehai, T., Lundqvist, M., Sherman, E., Mylonas, E., Weiner, L., Haran, G., Svergun, D.I., Mulder, F.A.A., Sussman, J.L., et al. (2008). Biophysical characterization of the unstructured cytoplasmic domain of the human neuronal adhesion protein neuroligin 3. Biophys. J. 95, 1928–1944. |
| Prothymosin alpha | 37.8 | 0.9 | MSDAAVDTSSEITTKDLKEKKEVVEEAENGRDAPANGNAENEENGEQEADNEVDEEEEEGGEEEEEEEEGDGEEEDGDEDEEAESATGKRAAEDDEDDDVDTKKQKTDEDD | Uversky, V.N., Gillespie, J.R., Millett, I.S., Khodyakova, A.V., Vasiliev, A.M., Chernovskaya, T.V., Vasilenko, R.N., Kozlovskaya, G.D., Dolgikh, D.A., Fink, A.L., et al. (1999). Natively Unfolded Human Prothymosin α Adopts Partially Folded Collapsed Conformation at Acidic pH. Biochemistry 38, 15009–15016. |

| | | | | |
|---|---|---|---|---|
| Fez1 | 36 | 1 | QIQEEEETLQDEEVWDALTDNYIPSLSEDWRDPNIEALNGNCSDTEIHEKEEEEFNEKSENDSGINEEPLLTADQVIEEIEEMMQNSPDPEEEEEVLEEEDGG | Alborghetti, M.R., Furlan, A.S., Silva, J.C., Paes Leme, A.F., Torriani, I.C.L., and Kobarg, J. (2010). Human FEZ1 Protein Forms a Disulfide Bond Mediated Dimer: Implications for Cargo Transport. J. Proteome Res. 9, 4595–4603. |
| HIV-TAT | 33 | 1.05 | MEPVDPRLEPWKHPGSQPRTACTNCYCKKCCFHCQVCFIRKALGISYGRKKRRQRRRAPQDSETHQVSPPKQPASQPRGDPTGPKESKKKVERETETHPVN | Foucault, M., Mayol, K., Receveur-Bréchot, V., Bussat, M.-C., Klinguer-Hamour, C., Verrier, B., Beck, A., Haser, R., Gouet, P., and Guillon, C. (2010). UV and X-ray structural studies of a 101-residue long Tat protein from a HIV-1 primary isolate and of its mutated, detoxified, vaccine candidate. Proteins 78, 1441–1456. |
| p531-91 | 28.7 | 0.3 | MEEPQSDPSVEPPLSQETFSDLWKLLPENNVLSPLPSQAMDDLMLSPDDIEQWFTEDPGPDEAPRMPEAAPPVAPAPAAPTPAAPAPAPSW | Wells, M., Tidow, H., Rutherford, T.J., Markwick, P., Jensen, M.R., Mylonas, E., Svergun, D.I., Blackledge, M., and Fersht, A.R. (2008). Structure of tumor suppressor p53 and its intrinsically disordered N-terminal transactivation domain. Proc. Natl. Acad. Sci. U. S. A. 105, 5762–5767. |
| Tau - ht40 | 65 | 3 | MAEPRQEFEVMEDHAGTYGLGDRKDQGGYTMHQDQEGDTDAGLKESPLQTPTEDGSEEPGSETSDAKSTPTAEDVTAPLVDEGAPGKQAAAQPHTEIPEGTTAEEAGIGDTPSLEDEAAGHVTQARMVSKSKDGTGSDDKKAKGADGKTKIATPRGAAPPGQKGQANATRIPAKTPPAPKTPPSSGEPPKSGDRSG | E. Mylonas, A. Hascher, P. Bernado´, M. Blackledge, E. Mandelkow and D. I. Svergun, Biochemistry, 2008, 47, 10345–10353. |

| | | | | |
|---|---|---|---|---|
| | | | YSSPGSPGTPGSRSRTPSLPTPPTREPKKVAVV RTPPKSPSSAKSRLQTAPVPMPDLKNVKSKIGST ENLKHQPGGGKVQIINKKLDLSNVQSKCGSKDNI KHVPGGGSVQIVYKPVDLSKVTSKCGSLGNIHH KPGGGQVEVKSEKLDFKDRVQSKIGSLDNITHVP GGGNKKIETHKLTFRENAKAKTDHGAEIVYKSPV VSGDTSPRHLSNVSSTGSIDMVDSPQLATLADE VSASLAKQGL | |
| Tau - K32 | 42 | 3 | SSPGSPGTPGSRSRTPSLPTPPTREPKKVAVVR TPPKSPSSAKSRLQTAPVPMPDLKNVKSKIGSTE NLKHQPGGGKVQIINKKLDLSNVQSKCGSKDNIK HVPGGGSVQIVYKPVDLSKVTSKCGSLGNIHHK PGGGQVEVKSEKLDFKDRVQSKIGSLDNITHVP GGGNKKIETHKLTFRENAKAKTDHGAEIVY | E. Mylonas, A. Hascher, P. Bernado´, M. Blackledge, E. Mandelkow and D. I. Svergun, Biochemistry, 2008, 47, 10345–10353. |
| Tau - K16 | 39 | 3 | SSPGSPGTPGSRSRTPSLPTPPTREPKKVAVVR TPPKSPSSAKSRLQTAPVPMPDLKNVKSKIGSTE NLKHQPGGGKVQIINKKLDLSNVQSKCGSKDNIK HVPGGGSVQIVYKPVDLSKVTSKCGSLGNIHHK PGGGQVEVKSEKLDFKDRVQSKIGSLDNITHVP GGGNKKIE | E. Mylonas, A. Hascher, P. Bernado´, M. Blackledge, E. Mandelkow and D. I. Svergun, Biochemistry, 2008, 47, 10345–10353. |
| Tau - K18 | 38 | 3 | QTAPVPMPDLKNVKSKIGSTENLKHQPGGGKVQ IINKKLDLSNVQSKCGSKDNIKHVPGGGSVQIVY KPVDLSKVTSKCGSLGNIHHKPGGGQVEVKSEK LDFKDRVQSKIGSLDNITHVPGGGNKKIE | E. Mylonas, A. Hascher, P. Bernado´, M. Blackledge, E. Mandelkow and D. I. Svergun, Biochemistry, 2008, 47, 10345–10353. |
| Tau - ht23 | 53 | 3 | MAEPRQEFEVMEDHAGTYGLGDRKDQGGYTM HQDQEGDTDAGLKAEEAGIGDTPSLEDEAAGHV TQARMVSKSKDGTGSDDKKAKGADGKTKIATPR GAAPPGQKGQANATRIPAKTPPAPKTPPSSGEP PKSGDRSGYSSPGSPGTPGSRSRTPSLPTPPTR EPKKVAVVRTPPKSPSSAKSRLQTAPVPMPDLK NVKSKIGSTENLKHQPGGGKVQIVYKPVDLSKVT SKCGSLGNIHHKPGGGQVEVKSEKLDFKDRVQS KIGSLDNITHVPGGGNKKIETHKLTFRENAKAKTD HGAEIVYKSPVVSGDTSPRHLSNVSSTGSIDMVD SPQLATLADEVSASLAKQGL | E. Mylonas, A. Hascher, P. Bernado´, M. Blackledge, E. Mandelkow and D. I. Svergun, Biochemistry, 2008, 47, 10345–10353. |

| | | | | |
|---|---|---|---|---|
| Tau - K27 | 37 | 2 | SSPGSPGTPGSRSRTPSLPTPPTREPKKVAVVR TPPKSPSSAKSRLQTAPVPMPDLKNVKSKIGSTE NLKHQPGGGSVQIVYKPVDLSKVTSKCGSLGNI HHKPGGGQVEVKSEKLDFKDRVQSKIGSLDNIT HVPGGGNKKIETHKLTFRENAKAKTDHGAEIVY | E. Mylonas, A. Hascher, P. Bernado´, M. Blackledge, E. Mandelkow and D. I. Svergun, Biochemistry, 2008, 47, 10345–10353. |
| Tau - K17 | 36 | 2 | SSPGSPGTPGSRSRTPSLPTPPTREPKKVAVVR TPPKSPSSAKSRLQTAPVPMPDLKNVKSKIGSTE NLKHQPGGGSVQIVYKPVDLSKVTSKCGSLGNI HHKPGGGQVEVKSEKLDFKDRVQSKIGSLDNIT HVPGGGNKKIE | E. Mylonas, A. Hascher, P. Bernado´, M. Blackledge, E. Mandelkow and D. I. Svergun, Biochemistry, 2008, 47, 10345–10353. |
| Tau - K19 | 35 | 1 | QTAPVPMPDLKNVKSKIGSTENLKHQPGGGSVQ IVYKPVDLSKVTSKCGSLGNIHHKPGGGQVEVKS EKLDFKDRVQSKIGSLDNITHVPGGGNKKIE | E. Mylonas, A. Hascher, P. Bernado´, M. Blackledge, E. Mandelkow and D. I. Svergun, Biochemistry, 2008, 47, 10345–10353. |
| Tau - K44 | 52 | 2 | MAEPRQEFEVMEDHAGTYGLGDRKDQGGYTM HQDQEGDTDAGLKAEEAGIGDTPSLEDEAAGHV TQARMVSKSKDGTGSDDKKAKGADGKTKIATPR GAAPPGQKGQANATRIPAKTPPAPKTPPSSGEP PKSGDRSGYSSPGSPGTPGSRSRTPSLPTPPTR EPKKVAVVRTPPKSPSSAKSRLQTAPVPMPDLK NVKSKIGSTENLKHQPGGGKVQIVYKPVDLSKVT SKCGSLGNIHHKPGGGQVEVKSEKLDFKDRVQS KIGSLDNITHVPGGGNKKIE | E. Mylonas, A. Hascher, P. Bernado´, M. Blackledge, E. Mandelkow and D. I. Svergun, Biochemistry, 2008, 47, 10345–10353. |
| Tau - K10 | 40 | 1 | QTAPVPMPDLKNVKSKIGSTENLKHQPGGGSVQ IVYKPVDLSKVTSKCGSLGNIHHKPGGGQVEVKS EKLDFKDRVQSKIGSLDNITHVPGGGNKKIETHK LTFRENAKAKTDHGAEIVYKSPVVSGDTSPRHLS NVSSTGSIDMVDSPQLATLADEVSASLAKQGL | E. Mylonas, A. Hascher, P. Bernado´, M. Blackledge, E. Mandelkow and D. I. Svergun, Biochemistry, 2008, 47, 10345–10353. |
| Tau - K25 | 41 | 2 | MAEPRQEFEVMEDHAGTYGLGDRKDQGGYTM HQDQEGDTDAGLKAEEAGIGDTPSLEDEAAGHV TQARMVSKSKDGTGSDDKKAKGADGKTKIATPR GAAPPGQKGQANATRIPAKTPPAPKTPPSSGEP | E. Mylonas, A. Hascher, P. Bernado´, M. Blackledge, E. Mandelkow and D. I. Svergun, |

| | | | | |
|---|---|---|---|---|
| | | | PKSGDRSGYSSPGSPGTPGSRSRTPSLPTPPTR EPKKVAVVRTPPKSPSSAKSRL | Biochemistry, 2008, 47, 10345–10353. |
| Tau - K23 | 49 | 2 | MAEPRQEFEVMEDHAGTYGLGDRKDQGGYTM HQDQEGDTDAGLKAEEAGIGDTPSLEDEAAGHV TQARMVSKSKDGTGSDDKKAKGADGKTKIATPR GAAPPGQKGQANATRIPAKTPPAPKTPPSSGEP PKSGDRSGYSSPGSPGTPGSRSRTPSLPTPPTR EPKKVAVVRTPPKSPSSAKSRLKKIETHKLTFREN AKAKTDHGAEIVYKSPVVSGDTSPRHLSNVSST GSIDMVDSPQLATLADEVSASLAKQGL | E. Mylonas, A. Hascher, P. Bernado´, M. Blackledge, E. Mandelkow and D. I. Svergun, Biochemistry, 2008, 47, 10345–10353. |
| Tau - K32 AT8 AT100 | 41 | 3 | SEPGEPGEPGSRSREPELPTPPTREPKKVAVVR TPPKSPSSAKSRLQTAPVPMPDLKNVKSKIGSTE NLKHQPGGGKVQIINKKLDLSNVQSKCGSKDNIK HVPGGGSVQIVYKPVDLSKVTSKCGSLGNIHHK PGGGQVEVKSEKLDFKDRVQSKIGSLDNITHVP GGGNKKIETHKLTFRENAKAKTDHGAEIVY | E. Mylonas, A. Hascher, P. Bernado´, M. Blackledge, E. Mandelkow and D. I. Svergun, Biochemistry, 2008, 47, 10345–10353. |
| Tau - ht23 S214E | 54 | 3 | MAEPRQEFEVMEDHAGTYGLGDRKDQGGYTM HQDQEGDTDAGLKAEEAGIGDTPSLEDEAAGHV TQARMVSKSKDGTGSDDKKAKGADGKTKIATPR GAAPPGQKGQANATRIPAKTPPAPKTPPSSGEP PKSGDRSGYSSPGSPGTPGSRSRTPELPTPPTR EPKKVAVVRTPPKSPSSAKSRLQTAPVPMPDLK NVKSKIGSTENLKHQPGGGKVQIVYKPVDLSKVT SKCGSLGNIHHKPGGGQVEVKSEKLDFKDRVQS KIGSLDNITHVPGGGNKKIETHKLTFRENAKAKTD HGAEIVYKSPVVSGDTSPRHLSNVSSTGSIDMVD SPQLATLADEVSASLAKQGL | E. Mylonas, A. Hascher, P. Bernado´, M. Blackledge, E. Mandelkow and D. I. Svergun, Biochemistry, 2008, 47, 10345–10353. |
| Tau - ht23 AT8 AT100 | 52 | 3 | MAEPRQEFEVMEDHAGTYGLGDRKDQGGYTM HQDQEGDTDAGLKAEEAGIGDTPSLEDEAAGHV TQARMVSKSKDGTGSDDKKAKGADGKTKIATPR GAAPPGQKGQANATRIPAKTPPAPKTPPSSGEP PKSGDRSGYSEPGEPGEPGSRSREPELPTPPT REPKKVAVVRTPPKSPSSAKSRLQTAPVPMPDL KNVKSKIGSTENLKHQPGGGKVQIVYKPVDLSKV TSKCGSLGNIHHKPGGGQVEVKSEKLDFKDRVQ SKIGSLDNITHVPGGGNKKIETHKLTFRENAKAKT | E. Mylonas, A. Hascher, P. Bernado´, M. Blackledge, E. Mandelkow and D. I. Svergun, Biochemistry, 2008, 47, 10345–10353. |

| | | | | |
|---|---|---|---|---|
| | | | DHGAEIVYKSPVVSGDTSPRHLSNVSSTGSIDMV DSPQLATLADEVSASLAKQGL | |
| Tau - K18 P301L | 35 | 2 | QTAPVPMPDLKNVKSKIGSTENLKHQPGGGKVQ IINKKLDLSNVQSKCGSKDNIKHVLGGGSVQIVYK PVDLSKVTSKCGSLGNIHHKPGGGQVEVKSEKL DFKDRVQSKIGSLDNITHVPGGGNKKIE | E. Mylonas, A. Hascher, P. Bernado´, M. Blackledge, E. Mandelkow and D. I. Svergun, Biochemistry, 2008, 47, 10345–10353. |
| Tau - K18 ΔK280 | 79 | 10 | QTAPVPMPDLKNVKSKIGSTENLKHQPGGGKVQ IINKLDLSNVQSKCGSKDNIKHVLGGGSVQIVYKP VDLSKVTSKCGSLGNIHHKPGGGQVEVKSEKLD FKDRVQSKIGSLDNITHVPGGGNKKIE | E. Mylonas, A. Hascher, P. Bernado´, M. Blackledge, E. Mandelkow and D. I. Svergun, Biochemistry, 2008, 47, 10345–10353. |
| Tau - K18 ΔK280 I277P I308P | 35 | 2 | QTAPVPMPDLKNVKSKIGSTENLKHQPGGGKVQ PINKLDLSNVQSKCGSKDNIKHVLGGGSVQPVY KPVDLSKVTSKCGSLGNIHHKPGGGQVEVKSEK LDFKDRVQSKIGSLDNITHVPGGGNKKIE | E. Mylonas, A. Hascher, P. Bernado´, M. Blackledge, E. Mandelkow and D. I. Svergun, Biochemistry, 2008, 47, 10345–10353. |
| Histatin | 13.2 | 0.01 | DSHAKRHHGYKRKFHEKHHSHRGY | Cragnell, C., Durand, D., Cabane, B., and Skepö, M. (2016). Coarse-grained modeling of the intrinsically disordered protein Histatin 5 in solution: Monte Carlo simulations in combination with SAXS. Proteins 84, 777–791. |
| CortactinCR | 46.7 | | GPLGSGYGGKFGVEQDRMDKSAVGHEYQSKLS KHCSQVDSVRGFGGKFGVQMDRVDQSAVGFEY QGKTEKHASQKDYSSGFGGKYGVQADRVDKSA VGFDYQGKTEKHESQRDYSKGFGGKYGIDKDK VDKSAVGFEYQGKTEKHESQKDYVKGFGGKFG VQTDRQDKCALGWDHQEKLQLHESQKDYKTGF GGKFGVQSERQDSAAVGFDYKEKLAKHESQQD YSKGFGGKYGVQKDRMDKNASTFEDVTQVSSA YQKTVPVEAVTSKTSNIRANFENLAKEKEQEDRR | Li, X., Tao, Y., Murphy, J.W., Scherer, A.N., Lam, T.T., Marshall, A.G., Koleske, A.J., and Boggon, T.J. (2017). The repeat region of cortactin is intrinsically disordered in solution. Sci. Rep. 7, 16696. |

| | | | | |
|---|---|---|---|---|
| | | | KAEAERAQRMAKERQEQEEARRKLEEQARAKTQT | |
| Pertactin-NTD | 51.3 | 0.1 | DWNNQSIVKTGERQHGIHIQGSDPGGVRTASGTTIKVSGRQAQGILLENPAAELQFRNGSVTSSGQLSDDGIRRFLGTVTVKAGKLVADHATLANVGDTWDDDGIALYVAGEQAQASIADSTLQGAGGVQIERGANVTVQRSAIVDGGLHIGALQSLQPEDLPPSRVVLRDTNVTAVPASGAPAAVSVLGASELTLDGGHITGGRAAGVAAMQGAVVHLQRATIRRGEALAGGAVPGGAVPGGAVPGGFGPGGFGPVLDGWYGVDVSGSSVELAQSIVEAPELGAAIRVGRGARVTVPGGSLSAPHGNVIETGGARRFAPQAAPLSITLQAGAH | Riback, J.A., Bowman, M.A., Zmyslowski, A.M., Knoverek, C.R., Jumper, J.M., Hinshaw, J.R., Kaye, E.B., Freed, K.F., Clark, P.L., and Sosnick, T.R. (2017). Innovative scattering analysis shows that hydrophobic disordered proteins are expanded in water. Science 358, 238–241. |
| Reduced_Rnase H | 33.6 | 0.1 | KETAAAKFERQHMDSSTSAASSSNYCNQMMKSRNLTKDRCKPVNTFVHESLADVQAVCSQKNVACKNGQTNCYQSYSTMSITDCRETGSSKYPNCAYKTTQANKHIIVACEGNPYVPVHFDASV | Riback, J.A., Bowman, M.A., Zmyslowski, A.M., Knoverek, C.R., Jumper, J.M., Hinshaw, J.R., Kaye, E.B., Freed, K.F., Clark, P.L., and Sosnick, T.R. (2017). Innovative scattering analysis shows that hydrophobic disordered proteins are expanded in water. Science 358, 238–241. |
| Nup1573_frag | 24 | 5 | GCPSASPAFGANQTPTFGQSQGASQPNPPGFSISSSTALFPTGSQPAPPTFGTVSSSSQPPVFGQQPSQSAFGSTTPNA | Mercadante, D., Milles, S., Fuertes, G., Svergun, D.I., Lemke, E.A., and Gräter, F. (2015). Kirkwood-Buff Approach Rescues Overcollapse of a Disordered Protein in Canonical Protein Force Fields. J. Phys. Chem. B 119, 7975–7984. |
| LOX-PP | 37 | 0.4 | APPAAGQQQPPREPPAAPGAWRQQIQWENNGQVFSLLSLGSQYQPQRRRDPGAAVPGAANASAQQPRTPILLIRDNRTAAARTRTAGSSGVTAGRPRPTARHWFQAGYSTSRAREAGASRAENQTAPGEVPALSNLRPPSRVDGMVG | Vallet, S.D., Miele, A.E., Uciechowska-Kaczmarzyk, U., Liwo, A., Duclos, B., Samsonov, S.A., and Ricard-Blum, S. (2018). Insights into the structure and dynamics of lysyl |

| | | | | |
|---|---|---|---|---|
| | | | | oxidase propeptide, a flexible protein with numerous partners. Sci. Rep. 8, 11768. |
| H1_CTD | 25 | 0.2 | KGDEPKRSVAFKKTKKEVKKVATPKKAAKPKKAASKAPSKKPKATPVKKAKKKPAATPKKAKKPKVVKVKPVKASKPKKAKTVKPKAKSSAKRASKKK | Roque, A., Ponte, I., and Suau, P. (2007). Macromolecular crowding induces a molten globule state in the C-terminal domain of histone H1. Biophys. J. 93, 2170–2177. |
| p27_WT (v31) | 28.1 | 1.8 | GSHMKGACKVPAQESQDVSGSRPAAPLIGAPANSEDTHLVDPKTDPSDSQTGLAEQCAGIRKRPATDDSSTQNKRANRTEENVSDGSPNAGSVEQTPKKPGLRRRQT | Das, R.K., Huang, Y., Phillips, A.H., Kriwacki, R.W., and Pappu, R.V. (2016). Cryptic sequence features within the disordered protein p27Kip1 regulate cell cycle signaling. Proc. Natl. Acad. Sci. U. S. A. 113, 5616–5621. |
| p27_v14 | 29.4 | 1.3 | GSHMKGACKSSSPPSNDQGRPGDPKQVIDKTEVERTQDTSNIQETQSANNSGPDKPSRCDLAVSGVAAAALPAPGHANSTARDLTRDEEAGSVEQTPKKPGLRRRQT | Das, R.K., Huang, Y., Phillips, A.H., Kriwacki, R.W., and Pappu, R.V. (2016). Cryptic sequence features within the disordered protein p27Kip1 regulate cell cycle signaling. Proc. Natl. Acad. Sci. U. S. A. 113, 5616–5621. |
| p27_v15 | 29.2 | 1 | GSHMKGACIVANSPPDDVKSKEDVPQTDPRLTGGDRDNARASRTGNDPAGASTQSAEVACSNPILSTPDAQEKQAGTSNSKERPHEQLSAGSVEQTPKKPGLRRRQT | Das, R.K., Huang, Y., Phillips, A.H., Kriwacki, R.W., and Pappu, R.V. (2016). Cryptic sequence features within the disordered protein p27Kip1 regulate cell cycle signaling. Proc. Natl. Acad. Sci. U. S. A. 113, 5616–5621. |

| | | | | |
|---|---|---|---|---|
| p27_v44 | 24.9 | 1.3 | GSHMKGACRKPANAEADSSSCQNVPRGKSKQAPETPTGSPLGDATLNQVKPRRPSSASTNIGQLEDADEDDAEDHVGSAVTSQTIPNDRAGSVEQTPKKPGLRRRQT | Das, R.K., Huang, Y., Phillips, A.H., Kriwacki, R.W., and Pappu, R.V. (2016). Cryptic sequence features within the disordered protein p27Kip1 regulate cell cycle signaling. Proc. Natl. Acad. Sci. U. S. A. 113, 5616–5621. |
| p27_v56 | 23.3 | 1 | GSHMKGACGSSVLGTGNPRNQAHVSDTSLEEDDDEQDDSTPDEVSQACTIVASALDINAATPRSPKASPKRKRKRQSTAPAQGNEPPGNAGSVEQTPKKPGLRRRQT | Das, R.K., Huang, Y., Phillips, A.H., Kriwacki, R.W., and Pappu, R.V. (2016). Cryptic sequence features within the disordered protein p27Kip1 regulate cell cycle signaling. Proc. Natl. Acad. Sci. U. S. A. 113, 5616–5621. |
| p27_v78 | 22.1 | 0.3 | GSHMKGACALPSGVVPAEDDDDDEEEEDDQDPAQPQAVQGAAPSSGTNNSQPILPSIAVNSTTGPNSTAGKKKRKRRRTRHSNCATLSSAGSVEQTPKKPGLRRRQT | Das, R.K., Huang, Y., Phillips, A.H., Kriwacki, R.W., and Pappu, R.V. (2016). Cryptic sequence features within the disordered protein p27Kip1 regulate cell cycle signaling. Proc. Natl. Acad. Sci. U. S. A. 113, 5616–5621. |
| Ki-1/57 | 47 | 2 | PRRGEQQGWNDSRGPEGMLERAERRSYREYRPYETERQADFTAEKFPDEKPGDRFDRDRPLRGRGGPRGGMRGRGRGGPGNRVFDAFDQRGKREFERYGGNDKIAVRTEDNMGGCGVRTWGSGKDTSDVEPTAPMEEPTVVEESQGTPEEESPAKVPELEVEEETQVQEMTLDEWKNLQEQTRPKPEFNIRKPESTVPSKAVVIHKSKYRDDMVKDDYEDDSHVFRKPANDITSQLEINFGNLPRPGRGARGGTRGGRGRIRRAENYGPRAEVVMQDVAPNPDDPEDFPALS | Bressan, G.C., Silva, J.C., Borges, J.C., Dos Passos, D.O., Ramos, C.H.I., Torriani, I.L., and Kobarg, J. (2008). Human regulatory protein Ki-1/57 has characteristics of an intrinsically unstructured protein. J. Proteome Res. 7, 4465–4474. |
| CTCF-R domain (WT) | 32.5 | 1.8 | SAERRNSILTETLHRFSLEGDAPVSWTETKKQSFKQTGEFGEKRKNSILNPINSIRKFSIVQKTPLQMN | Marasini, C., Galeno, L., and Moran, O. (2013). A |

| | | | | |
|---|---|---|---|---|
| | | | GIEEDSDEPLERRLSLVPDSEQGEAILPRISVISTGPTLQARRRQSVLNLMTHSVNQGQNIHRKTTASTRKVSLAPQANLTELDIYSRRLSQETGLEISEEINEEDLKECFFDDME | SAXS-based ensemble model of the native and phosphorylated regulatory domain of the CFTR. Cell. Mol. Life Sci. 70, 923–933. |
| CTCF-R domain (phosphorylated) | 29.2 | 0.4 | SAERRNSILTETLHRFSLEGDAPVSWTETKKQSFKQTGEFGEKRKNSILNPINSIRKFSIVQKTPLQMNGIEEDSDEPLERRLSLVPDSEQGEAILPRISVISTGPTLQARRRQSVLNLMTHSVNQGQNIHRKTTASTRKVSLAPQANLTELDIYSRRLSQETGLEISEEINEEDLKECFFDDME | Marasini, C., Galeno, L., and Moran, O. (2013). A SAXS-based ensemble model of the native and phosphorylated regulatory domain of the CFTR. Cell. Mol. Life Sci. 70, 923–933. |
| hNHE1cdt | 37.5 | 0 | VPAHKLDSPTMSRARIGSDPLAYEPKEDLPVITIDPASPQSPESVDLVNEELKGKVLGLSRDPAKVAEEDEDDDGGIMMRSKETSSPGTDDVFTPAPSDSPSSQRIQRCLSDPGPHPEPGEGEPFFPKGQ | Kjaergaard, M., Nørholm, A.-B., Hendus-Altenburger, R., Pedersen, S.F., Poulsen, F.M., and Kragelund, B.B. (2010). Temperature-dependent structural changes in intrinsically disordered proteins: Formation of α-helices or loss of polyproline II? Protein Sci. 19, 1555–1564. |
| pMBP | 54 | 0 | ASQKRPSQRHGSKYLASASTMDHARHGFLPRHRDTGIDSLGRFFGADRGAPKRGSGKDGHHAARTTHYGSLPQKAQHGRPQDENPVVHFFKNIVTPRTPPPSQGKGRGLSLSRFSWGAEGQKPGFGYGGRAPDYKPAHKGLKGAQDAQGTLSKIFKLGGRDSRSGSPMARR | Majava, V., Wang, C., Myllykoski, M., Kangas, S.M., Kang, S.U., Hayashi, N., Baumgärtel, P., Heape, A.M., Lubec, G., and Kursula, P. (2010). Structural analysis of the complex between calmodulin and full-length myelin basic protein, an intrinsically disordered molecule. Amino Acids 39, 59–71. |
| HMPV | 27.4 | 0.5 | MSFPEGKDILFMGNEAAKLAEAFQKSLRKPSHKRSQSIIGEKVNTVSETLELPTISRPTKP | Renner, M., Paesen, G.C., Grison, C.M., Granier, S., Grimes, J.M., and Leyrat, C. |

| | | | | |
|---|---|---|---|---|
| | | | | (2017). Structural dissection of human metapneumovirus phosphoprotein using small angle x-ray scattering. Sci. Rep. 7, 14865. |
| redAFP | 22.2 | 0.1 | CKGADGAHGVNGCPGTAGAAGSVGGPGCDGGHGGNGGNGNPGCAGGVGGAGGASGGTGVGGRGGKGGSGTPKGADGAPGAP | Gates, Z.P., Baxa, M.C., Yu, W., Riback, J.A., Li, H., Roux, B., Kent, S.B.H., and Sosnick, T.R. (2017). Perplexing cooperative folding and stability of a low-sequence complexity, polyproline 2 protein lacking a hydrophobic core. Proc. Natl. Acad. Sci. U. S. A. 114, 2241–2246. |
| CSD1 (with overhang) | 35.4 | 0 | MAMITNSSSVPAESKSSKPSGKSDMDAALDDLIDTLGGPEETEEDNTTYTGPEVLDPMSSTYIEELGKREVTLPPKYRELLDKKEGIPVPPPDTSKPLGPDDAIDALSLDLTCSSPTADGKKTEKEKSTGEVLKAQSVGVIKSDPLESLN | Konno, T., Tanaka, N., Kataoka, M., Takano, E., and Maki, M. (1997). A circular dichroism study of preferential hydration and alcohol effects on a denatured protein, pig calpastatin domain I. Biochim. Biophys. Acta 1342, 73–82. |
| PAGE4_WT | 36.2 | 1.1 | MSARVRSRSRGRGDGQEAPDVVAFVAPGESQQEEPPTDNQDIEPGQEREGTPPIEERKVEGDCQEMDLEKTRSERGDGSDVKEKTPPNPKHAKTKEAGDGQP | Kulkarni, P., Jolly, M.K., Jia, D., Mooney, S.M., Bhargava, A., Kagohara, L.T., Chen, Y., Hao, P., He, Y., Veltri, R.W., et al. (2017). Phosphorylation-induced conformational dynamics in an intrinsically disordered protein and potential role in phenotypic heterogeneity. Proc. Natl. Acad. Sci. U. S. A. 114, E2644–E2653. |

| | | | | |
|---|---|---|---|---|
| PAGE4_WT_phosphorylated | 49.8 | 1.9 | MSARVRSRSRGRGDGQEAPDVVAFVAPGESQQEEPPTDNQDIEPGQEREGTPPIEERKVEGDCQEMDLEKTRSERGDGSDVKEKTPPNPKHAKTKEAGDGQP | Kulkarni, P., Jolly, M.K., Jia, D., Mooney, S.M., Bhargava, A., Kagohara, L.T., Chen, Y., Hao, P., He, Y., Veltri, R.W., et al. (2017). Phosphorylation-induced conformational dynamics in an intrinsically disordered protein and potential role in phenotypic heterogeneity. Proc. Natl. Acad. Sci. U. S. A. 114, E2644–E2653. |
| ERalpha-NTD | 31 | 0.2 | SNAMTMTLHTKASGMALLHQIQGNELEPLNRPQLKIPLERPLGEVYLDSSKPAVYNYPEGAAYEFNAAAAANAQVYGQTGLPYGPGSEAAAFGSNGLGGFPPLNSVSPSPLMLLHPPPQLSPFLQPHGQQVPYYLENEPSGYTVREAGPPAFYRPNSDNRRQGGRERLASTNDKGSMAMESAKETRY | Peng, Y., Cao, S., Kiselar, J., Xiao, X., Du, Z., Hsieh, A., Ko, S., Chen, Y., Agrawal, P., Zheng, W., Shi, W., Jiang, W., Yang, L., Chance, M. R., Surewicz, W. K., Buck, M., & Yang, S. (2019). A Metastable Contact and Structural Disorder in the Estrogen Receptor Transactivation Domain. Structure , 27(2), 229–240.e4. |
| A1-LCD-NLS | 27.6 | 0.16 | GSMASASSSQRGRSGSGNFGGGRGGGFGGNDNFGRGGNFSGRGGFGGSRGGGGYGGSGDGYNGFGNDGSNFGGGGSYNDFGNYNNQSSNFGPMKGGNFGGRSSGGSGGGGQYFAKPRNQGGYGGSSSSSSYGSGRRF | Bremer, A., Farag, M., Borcherds, W. M., Peran, I., Martin, E. W., Pappu, R. V., & Mittag, T. (2022). Deciphering how naturally occurring sequence features impact the phase behaviours of disordered prion-like domains. Nature Chemistry, 14(2), 196–207. |
| A1-LCD+NLS | 25.83 | 0.11 | GSMASASSSQRGRSGSGNFGGGRGGGFGGNDNFGRGGNFSGRGGFGGSRGGGGYGGSGDGYNGFGNDGSNFGGGGSYNDFGNYNNQSSNFGPMKGGNFGGRSSGPYGGGGQYFAKPRNQGGYGGSSSSSSYGSGRRF | Bremer, A., Farag, M., Borcherds, W. M., Peran, I., Martin, E. W., Pappu, R. V., & Mittag, T. (2022). Deciphering how naturally occurring |

| | | | | |
|---|---|---|---|---|
| | | | | sequence features impact the phase behaviours of disordered prion-like domains. Nature Chemistry, 14(2), 196–207. |
| A1-LCD-12F+12Y | 26.04 | 0.2 | GSMASASSSQRGRSGSGNYGGGRGGGYGGNDNYGRGGNYSGRGGYGGSRGGGGYGGSGDGYNGYGNDGSNYGGGGSYNDYGNYNNQSSNYGPMKGGNYGGRSSGGSGGGGQYYAKPRNQGGYGGSSSSSSYGSGRRY | Bremer, A., Farag, M., Borcherds, W. M., Peran, I., Martin, E. W., Pappu, R. V., & Mittag, T. (2022). Deciphering how naturally occurring sequence features impact the phase behaviours of disordered prion-like domains. Nature Chemistry, 14(2), 196–207. |
| A1-LCD+7F-7Y | 27.18 | 0.13 | GSMASASSSQRGRSGSGNFGGGRGGGFGGNDNFGRGGNFSGRGGFGGSRGGGGFGGSGDGFNGFGNDGSNFGGGGSFNDFGNFNNQSSNFGPMKGGNFGGRSSGGSGGGGQFFAKPRNQGGFGGSSSSSSFGSGRRF | Bremer, A., Farag, M., Borcherds, W. M., Peran, I., Martin, E. W., Pappu, R. V., & Mittag, T. (2022). Deciphering how naturally occurring sequence features impact the phase behaviours of disordered prion-like domains. Nature Chemistry, 14(2), 196–207. |
| A1-LCD-9F+6Y | 26.55 | 0.1 | GSMASASSSQRGRSGSGNFGGGRGGGYGGNDNYGRGGNYSGRGGFGGSRGGGGYGGSGDGYNGGGNDGSNYGGGGSYNDSGNYNNQSSNFGPMKGGNYGGRSSGGSGGGGQYGAKPRNQGGYGGSSSSSSYGSGRRY | Bremer, A., Farag, M., Borcherds, W. M., Peran, I., Martin, E. W., Pappu, R. V., & Mittag, T. (2022). Deciphering how naturally occurring sequence features impact the phase behaviours of disordered prion-like domains. Nature Chemistry, 14(2), 196–207. |
| A1-LCD-8F+4Y | 27.07 | 0.07 | GSMASASSSQRGRSGSGNFGGGRGGGYGGNDNGGRGGNYSGRGGFGGSRGGGGYGGSGDGYNGGGNDGSNYGGGGSYNDSGNYNNQSSNFGPMKGGNYGGRSSGGSGGGGQYGAKPRNQGGYGGSSSSSSYGSGRRF | Bremer, A., Farag, M., Borcherds, W. M., Peran, I., Martin, E. W., Pappu, R. V., & Mittag, T. (2022). Deciphering how naturally occurring sequence features impact the |

| | | | | |
|---|---|---|---|---|
| | | | | phase behaviours of disordered prion-like domains. Nature Chemistry, 14(2), 196–207. |
| A1-LCD-9F+3Y | 26.83 | 0.13 | GSMASASSSQRGRSGSGNFGGGRGGGYGGND NGGRGGNYSGRGGFGGSRGGGGYGGSGDGY NGGGNDGSNYGGGGSYNDSGNGNNQSSNFGP MKGGNYGGRSSGGSGGGGQYGAKPRNQGGY GGSSSSSSYGSGRRS | Bremer, A., Farag, M., Borcherds, W. M., Peran, I., Martin, E. W., Pappu, R. V., & Mittag, T. (2022). Deciphering how naturally occurring sequence features impact the phase behaviours of disordered prion-like domains. Nature Chemistry, 14(2), 196–207. |
| A1-LCD-10R | 26.71 | 0.07 | GSMASASSSQGGSSGSGNFGGGGGGFGGND NFGGGGNFSGSGGFGGSGGGGGYGGSGDGY NGFGNDGSNFGGGGSYNDFGNYNNQSSNFGP MKGGNFGGSSSGPYGGGGQYFAKPGNQGGYG GSSSSSSYGSGGGF | Bremer, A., Farag, M., Borcherds, W. M., Peran, I., Martin, E. W., Pappu, R. V., & Mittag, T. (2022). Deciphering how naturally occurring sequence features impact the phase behaviours of disordered prion-like domains. Nature Chemistry, 14(2), 196–207. |
| A1-LCD-6R | 25.73 | 0.09 | GSMASASSSQGGRSGSGNFGGGRGGGFGGND NFGGGGNFSGSGGFGGSRGGGGYGGSGDGY NGFGNDGSNFGGGGSYNDFGNYNNQSSNFGP MKGGNFGGSSSGPYGGGGQYFAKPGNQGGYG GSSSSSSYGSGGRF | Bremer, A., Farag, M., Borcherds, W. M., Peran, I., Martin, E. W., Pappu, R. V., & Mittag, T. (2022). Deciphering how naturally occurring sequence features impact the phase behaviours of disordered prion-like domains. Nature Chemistry, 14(2), 196–207. |
| A1-LCD+2R | 26.23 | 0.23 | GSMASASSSQRGRSGSGNFGGGRGGGFGGND NFGRGGNFSGRGGFGGSRGGGGYGGSGDGY NGFRNDGSNFGGGGRYNDFGNYNNQSSNFGP MKGGNFGGRSSGPYGGGGQYFAKPRNQGGYG GSSSSSSYGSGRRF | Bremer, A., Farag, M., Borcherds, W. M., Peran, I., Martin, E. W., Pappu, R. V., & Mittag, T. (2022). Deciphering how naturally occurring sequence features impact the phase behaviours of disordered |

| | | | | |
|---|---|---|---|---|
| | | | | prion-like domains. Nature Chemistry, 14(2), 196–207. |
| A1-LCD+7R | 27.09 | 0.07 | GSMASASSSQRGRSGRGNFGGGRGGGFGGNDNFGRGGNFSGRGGFGGSRGGGRYGGSGDRYNGFGNDGRNFGGGGSYNDFGNYNNQSSNFGPMKGGNFRGRSSGPYGRGGQYFAKPRNQGGYGGSSSSRSYGSGRRF | Bremer, A., Farag, M., Borcherds, W. M., Peran, I., Martin, E. W., Pappu, R. V., & Mittag, T. (2022). Deciphering how naturally occurring sequence features impact the phase behaviours of disordered prion-like domains. Nature Chemistry, 14(2), 196–207. |
| A1-LCD-3R+3K | 26.34 | 0.15 | GSMASASSSQRGKSGSGNFGGGRGGGFGGNDNFGRGGNFSGRGGFGGSKGGGGYGGSGDGYNGFGNDGSNFGGGGSYNDFGNYNNQSSNFGPMKGGNFGGRSSGGSGGGGQYFAKPRNQGGYGGSSSSSSYGSGRKF | Bremer, A., Farag, M., Borcherds, W. M., Peran, I., Martin, E. W., Pappu, R. V., & Mittag, T. (2022). Deciphering how naturally occurring sequence features impact the phase behaviours of disordered prion-like domains. Nature Chemistry, 14(2), 196–207. |
| A1-LCD-6R+6K | 27.87 | 0.08 | GSMASASSSQKGKSGSGNFGGGRGGGFGGNDNFGKGGNFSGRGGFGGSKGGGGYGGSGDGYNGFGNDGSNFGGGGSYNDFGNYNNQSSNFGPMKGGNFGGKSSGGSGGGGQYFAKPRNQGGYGGSSSSSSYGSGRKF | Bremer, A., Farag, M., Borcherds, W. M., Peran, I., Martin, E. W., Pappu, R. V., & Mittag, T. (2022). Deciphering how naturally occurring sequence features impact the phase behaviours of disordered prion-like domains. Nature Chemistry, 14(2), 196–207. |
| A1-LCD-10R+10K | 28.49 | 0.05 | GSMASASSSQKGKSGSGNFGGGKGGGFGGNDNFGKGGNFSGKGGFGGSKGGGGYGGSGDGYNGFGNDGSNFGGGGSYNDFGNYNNQSSNFGPMKGGNFGGKSSGGSGGGGQYFAKPKNQGGYGGSSSSSSYGSGKKF | Bremer, A., Farag, M., Borcherds, W. M., Peran, I., Martin, E. W., Pappu, R. V., & Mittag, T. (2022). Deciphering how naturally occurring sequence features impact the phase behaviours of disordered |

| | | | | |
|---|---|---|---|---|
| | | | | prion-like domains. Nature Chemistry, 14(2), 196–207. |
| A1-LCD-4D | 26.42 | 0.12 | GSMASASSSQRGRSGSGNFGGGRGGGFGGNGNFGRGGNFSGRGGFGGSRGGGGYGGSGGGYNGFGNSGSNFGGGGSYNGFGNYNNQSSNFGPMKGGNFGGRSSGPYGGGGQYFAKPRNQGGYGGSSSSSSYGSGRRF | Bremer, A., Farag, M., Borcherds, W. M., Peran, I., Martin, E. W., Pappu, R. V., & Mittag, T. (2022). Deciphering how naturally occurring sequence features impact the phase behaviours of disordered prion-like domains. Nature Chemistry, 14(2), 196–207. |
| A1-LCD+4D | 27.18 | 0.3 | GSMASASSSQRDRSGSGNFGGGRGGGFGGNDNFGRGGNFSGRGDFGGSRGGGGYGGSGDGYNGFGNDGSNFGGGGSYNDFGNYNNQSSNFGPMKGGNFGGRSSDPYGGGGQYFAKPRNQGGYGGSSSSSSYDSGRRF | Bremer, A., Farag, M., Borcherds, W. M., Peran, I., Martin, E. W., Pappu, R. V., & Mittag, T. (2022). Deciphering how naturally occurring sequence features impact the phase behaviours of disordered prion-like domains. Nature Chemistry, 14(2), 196–207. |
| A1-LCD+8D | 26.85 | 0.07 | GSMASASSSQRDRSGSGNFGGGRDGGFGGNDNFGRGDNFSGRGDFGGSRDGGGYGGSGDGYNGFGNDGSNFGGGGSYNDFGNYNNQSSNFGPMKGGNFGGRSSDPYGGGGQYFAKPRNQDGYGGSSSSSSYDSGRRF | Bremer, A., Farag, M., Borcherds, W. M., Peran, I., Martin, E. W., Pappu, R. V., & Mittag, T. (2022). Deciphering how naturally occurring sequence features impact the phase behaviours of disordered prion-like domains. Nature Chemistry, 14(2), 196–207. |
| A1-LCD+12D | 28.01 | 0.12 | GSMASADSSQRDRDDSGNFGDGRGGGFGGNDNFGRGGNFSDRGGFGGSRGDGGYGGDGDGYNGFGNDGSNFGGGGSYNDFGNYNNQSSNFDPMKGGNFGDRSSGPYDGGGQYFAKPRNQGGYGGSSSSSSYGSDRRF | Bremer, A., Farag, M., Borcherds, W. M., Peran, I., Martin, E. W., Pappu, R. V., & Mittag, T. (2022). Deciphering how naturally occurring sequence features impact the phase behaviours of disordered |

| | | | | |
|---|---|---|---|---|
| | | | | prion-like domains. Nature Chemistry, 14(2), 196–207. |
| A1-LCD+12E | 28.52 | 0.05 | GSMASAESSQREREESGNFGEGRGGGFGGND NFGRGGNFSERGGFGGSRGEGGYGGEGDGYN GFGNDGSNFGGGGSYNDFGNYNNQSSNFEPM KGGNFGERSSGPYEGGGQYFAKPRNQGGYGG SSSSSSYGSERRF | Bremer, A., Farag, M., Borcherds, W. M., Peran, I., Martin, E. W., Pappu, R. V., & Mittag, T. (2022). Deciphering how naturally occurring sequence features impact the phase behaviours of disordered prion-like domains. Nature Chemistry, 14(2), 196–207. |
| A1-LCD+7R+10D | 29.21 | 0.08 | GSMASADSSQRDRDGRGNFGDGRGGGFGGND NFGRGGNFSDRGGFGGSRGGGRYGGDGDRYN GFGNDGRNFGGGGSYNDFGNYNNQSSNFDPM KGGNFRDRSSGPYDRGGQYFAKPRNQGGYGG SSSSRSYGSDRRF | Bremer, A., Farag, M., Borcherds, W. M., Peran, I., Martin, E. W., Pappu, R. V., & Mittag, T. (2022). Deciphering how naturally occurring sequence features impact the phase behaviours of disordered prion-like domains. Nature Chemistry, 14(2), 196–207. |
| A1-LCD+7K+12D blocky | 25.62 | 0.14 | GSMASAKSSQRDRDDDGNFGKGRGGGFGGNK NFGRGGNFSKRGGFGGSRGKGKYGGKGDDYN GFGNDGDNFGGGGSYNDFGNYNNQSSNFDPM DGGNFDDRSSGPYDDGGQYFADPRNQGGYGG SSSSKSYGSKRRF | Bremer, A., Farag, M., Borcherds, W. M., Peran, I., Martin, E. W., Pappu, R. V., & Mittag, T. (2022). Deciphering how naturally occurring sequence features impact the phase behaviours of disordered prion-like domains. Nature Chemistry, 14(2), 196–207. |
| A1-LCD-12F+12Y 10R | 26.07 | 0.2 | GSMASASSSQGGSSGSGNYGGGGGGGYGGN DNYGGGGNYSGSGGYGGSGGGGGYGGSGDG YNGYGNDGSNYGGGGSYNDYGNYNNQSSNYG PMKGGNYGGSSSGPYGGGGQYYAKPGNQGGY GGSSSSSSYGSGGGY | Bremer, A., Farag, M., Borcherds, W. M., Peran, I., Martin, E. W., Pappu, R. V., & Mittag, T. (2022). Deciphering how naturally occurring sequence features impact the phase behaviours of disordered |

| | | | | |
|---|---|---|---|---|
| | | | | prion-like domains. Nature Chemistry, 14(2), 196–207. |
| A1-LCD10F+7R+12D | 28.6 | 0.04 | GSMASADSSQRDRDDRGNFGDGRGGGGGGNDNFGRGGNGSDRGGGGGSRGDGRYGGDGDRYNGGGNDGRNGGGGGSYNDGGNYNNQSSNGDPMKGGNGRDRSSGPYDRGGQYGAKPRNQGGYGGSSSSRSYGSDRRG | Bremer, A., Farag, M., Borcherds, W. M., Peran, I., Martin, E. W., Pappu, R. V., & Mittag, T. (2022). Deciphering how naturally occurring sequence features impact the phase behaviours of disordered prion-like domains. Nature Chemistry, 14(2), 196–207. |
| PNt | 51.1 | 0.13 | DWNNQSIVKTGERQHGIHIQGSDPGGVRTASGTTIKVSGRQAQGILLENPAAELQFRNGSVTSSGQLSDDGIRRFLGTVTVKAGKLVADHATLANVGDTWDDDGIALYVAGEQAQASIADSTLQGAGGVQIERGANVTVQRSAIVDGGLHIGALQSLQPEDLPPSRVVLRDTNVTAVPASGAPAAVSVLGASELTLDGGHITGGRAAGVAAMQGAVVHLQRATIRRGEALAGGAVPGGAVPGGAVPGGFGPGGFGPVLDGWYGVDVSGSSVELAQSIVEAPELGAAIRVGRGARVTVPGGSLSAPHGNVIETGGARRFAPQAAPLSITLQAGAH | Bowman, M. A., Riback, J. A., Rodriguez, A., Guo, H., Li, J., Sosnick, T. R., & Clark, P. L. (2020). Properties of protein unfolded states suggest broad selection for expanded conformational ensembles. Proceedings of the National Academy of Sciences, 117(38), 23356–23364. |
| Swap1 | 49.2 | 0.59 | DWNNQSIVKTGERQHGIHIQGSDPGGVRTASGTTIKVSGRQAQGILLENPAAELQFRNGSVTSSGQKSDDGIRRFLGTVTVLAGKLVADHATLANVGDTWDDDGIALYVAGEQAQASIADSTLQGAGGVQIERGANVTVQRSAIVLGGLHIGALQSLQPEDDPPSRVVLRDTNVTAVPASGAPAAVSVLGASLLTLDGGHITGGRAAGVAAMQGAVVHEQRATIRRGEALAGGAVPGGAVPGGAVPGGFGPGGFGPVLDGWYGVDVSGSSVELAQSIVEAPELGAAIRVGRGARVTVPGGSLSAPHGNVIETGGARRFAPQAAPLSITLQAGAH | Bowman, M. A., Riback, J. A., Rodriguez, A., Guo, H., Li, J., Sosnick, T. R., & Clark, P. L. (2020). Properties of protein unfolded states suggest broad selection for expanded conformational ensembles. Proceedings of the National Academy of Sciences, 117(38), 23356–23364. |
| Swap3 | 40.58 | 1.07 | DWNNQSIVKTGERQHGIHIQGSDPGGVRTASGTTIKVSGRQAQGILLENPAAELQFRNGSVTSSGQKSTDGTRRFLGDVIVKAGLLVADHATLANVGDTWDDDGIALYVAGEQAQASIADSTLQGAGGVQIERGANVDVLRLAIVDGGLHIGALQSQQPETSPPSRVVLRDTNVTAVPASGAPAAVSVQGASEQTLDGGAITG | Bowman, M. A., Riback, J. A., Rodriguez, A., Guo, H., Li, J., Sosnick, T. R., & Clark, P. L. (2020). Properties of protein unfolded states suggest broad selection for expanded |

| | | | | |
|---|---|---|---|---|
| | | | GRAAGVAAMLGHVVHLLRATIRRGEALAGGAVP GGAVPGGAVPGGFGPGGFGPVLDGWYGVDVS GSSVELAQSIVEAPELGAAIRVGRGARVTVPGGS LSAPHGNVIETGGARRFAPQAAPLSITLQAGAH | conformational ensembles. Proceedings of the National Academy of Sciences, 117(38), 23356–23364. |
| Swap4 | 53.37 | 0.17 | DWNNQSIVKTGERQHGIHIQGSDPGGVRTASGT TIKVSGRQAQGILLENPAAELQFRNGSVTSSGQL SFVGITRDLGRDTVKAGKLVADHATLANVGDTW DDDGIALYVAGEQAQASIADSTLQGAGGVQIERG ADVRVQREAIVDGGLHNGALQSLQPSILPPSTVV LRDTNVTAVPASGAPAAVLVSGASGLRLDGGHIH EGRAAGVAAMQGAVVTLQTATIRRGEALAGGAV PGGAVPGGAVPGGFGPGGFGPVLDGWYGVDV SGSSVELAQSIVEAPELGAAIRVGRGARVTVPGG SLSAPHGNVIETGGARRFAPQAAPLSITLQAGAH | Bowman, M. A., Riback, J. A., Rodriguez, A., Guo, H., Li, J., Sosnick, T. R., & Clark, P. L. (2020). Properties of protein unfolded states suggest broad selection for expanded conformational ensembles. Proceedings of the National Academy of Sciences, 117(38), 23356–23364. |
| Swap4.1 | 54.45 | 0.14 | DWNNQSIVKTGERQHGIHIQGSDPGGVRTASGT TIKVSGRQAQGILLENPAAELQFRNGSVTSSGQL SFVGITRRLGDDTVKAGKLVADHATLANVGDTW DDDGIALYVAGEQAQASIADSTLQGAGGVQIERG ADVEVQRRAIVDGGLHNGALQSLQPSILPPSTVV LRDTNVTAVPASGAPAAVLVSGASGLELDGGHIH RGRAAGVAAMQGAVVTLQTATIRRGEALAGGAV PGGAVPGGAVPGGFGPGGFGPVLDGWYGVDV SGSSVELAQSIVEAPELGAAIRVGRGARVTVPGG SLSAPHGNVIETGGARRFAPQAAPLSITLQAGAH | Bowman, M. A., Riback, J. A., Rodriguez, A., Guo, H., Li, J., Sosnick, T. R., & Clark, P. L. (2020). Properties of protein unfolded states suggest broad selection for expanded conformational ensembles. Proceedings of the National Academy of Sciences, 117(38), 23356–23364. |
| Swap5 | 48.71 | 0.34 | DWNNQSIVKTGERQHGIHIQGSDPGGVRTASGT TIKVSGRQAQGILLENPAAELQFRNGSVTSSGQL SDDGIEDFLGTVTVDAGELVADHATLANVGDTW DDDGIALYVAGEQAQASIADSTLQGAGGVQIEDG ANVTVQESAIVDGGLHIGALQSLQPRRLPPSRVV LRKTNVTAVPASGAPAAVSVLGASKLTLRGGHIT GGRAAGVAAMQGAVVHLQRATIRRGRALAGGAV PGGAVPGGAVPGGFGPGGFGPVLDGWYGVDV SGSSVELAQSIVEAPELGAAIRVGRGARVTVPGG SLSAPHGNVIETGGARRFAPQAAPLSITLQAGAH | Bowman, M. A., Riback, J. A., Rodriguez, A., Guo, H., Li, J., Sosnick, T. R., & Clark, P. L. (2020). Properties of protein unfolded states suggest broad selection for expanded conformational ensembles. Proceedings of the National Academy of Sciences, 117(38), 23356–23364. |
| Swap6 | 52.61 | 0.27 | DWNNQSIVKTGERQHGIHIQGSDPGGVRTASGT TIKVSGRQAQGILLENPAAELQFRNGSVTSSGQL SDRGIDRFLGTVTVEAGKLVADHATLANVGDTW | Bowman, M. A., Riback, J. A., Rodriguez, A., Guo, H., Li, J., Sosnick, T. R., & Clark, P. L. |

| | | | | |
|---|---|---|---|---|
| | | | DKDGIALYVAGRQAQASIADSTLQGAGGVQIREGANVTVQRSAIVDGGLHIGALQSLQPERLPPSDVVLRDTNVTAVPASGAPAAVSVLGASRLTLDGGHITGGDAAGVAAMQGAVVHLQRATIERGEALAGGAVPGGAVPGGAVPGGFGPGGFGPVLDGWYGVDVSGSSVELAQSIVEAPELGAAIRVGRGARVTVPGGSLSAPHGNVIETGGARRFAPQAAPLSITLQAGAH | (2020). Properties of protein unfolded states suggest broad selection for expanded conformational ensembles. Proceedings of the National Academy of Sciences, 117(38), 23356–23364. |
| sfAFP | 23.1 | 2 | CKGADGAHGVNGCPGTAGAAGSVGGPGCDGGHGGNGGNGNPGCAGGVGGAGGASGGTGVGGRGGKGGSGTPKGADGAPGAP | Gates ZP, Baxa MC, Yu W, Riback JA, Li H, Roux B, et al. Perplexing cooperative folding and stability of a low-sequence complexity, polyproline 2 protein lacking a hydrophobic core. Proc Natl Acad Sci U S A. 2017;114: 2241–2246. |
| FCP1 | 15.6 | 0.12 | ESSRESSNEDEGSSSEADEMAKALEAELNDLM | Gibbs, Eric B., and Scott A. Showalter. 2016. “Quantification of Compactness and Local Order in the Ensemble of the Intrinsically Disordered Protein FCP1.” The Journal of Physical Chemistry. B 120 (34): 8960–69. |
| RS-peptide | 12.62 | 0.07 | MYRSRSRSRSRSRSRSRS | **SAXS data – NMR data - Xiang, S., Gapsys, V., Kim, H.-Y., Bessonov, S., Hsiao, H.-H., Möhlmann, S., Klaukien, V., Ficner, R., Becker, S., Urlaub, H., Lührmann, R., de Groot, B., & Zweckstetter, M. (2013). Phosphorylation drives a dynamic switch in serine/arginine-rich proteins. Structure , 21(12), 2162–2174.** |

| | | | | |
|---|---|---|---|---|
| P1_100 | 29 | 0 | MAEEQARHVKNGLECIRALKAEPIGSLAIEEAMAAWSEISDNPGQERATCREEKAGSSGLSKPCLSAIGSTEGGAPRIRGQGPGESDDDAETLGIPPRNL | Naudi-Fabra, S., Tengo, M., Jensen, M. R., Blackledge, M., & Milles, S. (2021). Quantitative Description of Intrinsically Disordered Proteins Using Single-Molecule FRET, NMR, and SAXS. Journal of the American Chemical Society, 143(48), 20109–20121. |
| DSS1 | 25 | 0.1 | MSRAALPSLENLEDDDEFEDFATENWPMKDTELDTGDDTLWENNWDDEDIGDDDFSVQLQAELKKKGVAAC | Pesce, F., Newcombe, E. A., Seiffert, P., Tranchant, E. E., Olsen, J. G., Grace, C. R., Kragelund, B. B., & Lindorff-Larsen, K. (2022). Assessment of models for calculating the hydrodynamic radius of intrinsically disordered proteins. Biophysical Journal. https://doi.org/10.1016/j.bpj.2022.12.013 |
| GHR_ICD | 59.59 | 0.38 | SKQQRIKMLILPPVPVPKIKGIDPDLLKEGKLEEVNTILAIHDSYKPEFHSDDSWVEFIELDIDEPDEKTEESDTDRLLSSDHEKSHSNLGVKDGDSGRTSCCEPDILETDFNANDIHEGTSEVAQPQRLKGEADLLCLDQKNQNNSPYHDACPATQQPSVIQAEKNKPQPLPTEGAESTHQAAHIQLSNPSSLSNIDFYAQVSDITPAGSVVLSPGQKNKAGMSQCDMHPEMVSLCQENFLMDNAYFCEADAKKCIPVAPHIKVESHIQPSLNQEDIYITTESLTTAAGRPGTGEHVPGSEMPVPDYTSIHIVQSPQGLILNATALPLPDKEFLSSCGYVSTDQLNKIMP | Pesce, F., Newcombe, E. A., Seiffert, P., Tranchant, E. E., Olsen, J. G., Grace, C. R., Kragelund, B. B., & Lindorff-Larsen, K. (2022). Assessment of models for calculating the hydrodynamic radius of intrinsically disordered proteins. Biophysical Journal. https://doi.org/10.1016/j.bpj.2022.12.013 |
| NHE6cmdd | 32 | 0.2 | GPPLTTTLPACCGPIARCLTSPQAYENQEQLKDDDSDLILNDGDISLTYGDSTVNTEPATSSAPRRFMGNSSEDALDRELAFGDHELVIRGTRLVLPMDDSEPPLNLLDNTRHGPA | Pesce, F., Newcombe, E. A., Seiffert, P., Tranchant, E. E., Olsen, J. G., Grace, C. R., Kragelund, B. B., & Lindorff-Larsen, K. (2022). Assessment of models for |

| | | | | |
|---|---|---|---|---|
| | | | | calculating the hydrodynamic radius of intrinsically disordered proteins. Biophysical Journal. https://doi.org/10.1016/j.bpj.2022.12.013 |
| ANAC046 | 36 | 0.3 | NAPSTTITTTKQLSRIDSLDNIDHLLDFSSLPPLIDPGFLGQPGPSFSGARQQHDLKPVLHHPTTAPVDNTYLPTQALNFPYHSVHNSGSDFGYGAGSGNNNKGMIKLEHSLVSVSQETGLSSDVNTTATPEISSYPMMMNPAMMDGSKSACDGLDDLIFWEDLYTS | Pesce, F., Newcombe, E. A., Seiffert, P., Tranchant, E. E., Olsen, J. G., Grace, C. R., Kragelund, B. B., & Lindorff-Larsen, K. (2022). Assessment of models for calculating the hydrodynamic radius of intrinsically disordered proteins. Biophysical Journal. https://doi.org/10.1016/j.bpj.2022.12.013 |
| stath_NTD | 9.1 | 0.3 | DSSEEKFLRRIGRFG | Rieloff, E., & Skepö, M. (2020). Phosphorylation of a disordered peptide—Structural effects and force field inconsistencies. Journal of Chemical Theory and Computation. https://pubs.acs.org/doi/abs/10.1021/acs.jctc.9b01190 |
| A1_Aro_minus | 27.9 | 0.8 | GSMASASSSQRGRSGSGNSGGGRGGGFGGNDNFGRGGNSSGRGGFGGSRGGGGYGGSGDGYNGFGNDGSNSGGGGSSNDFGNYNNQSSNFGPMKGGNFGGRSSGGSGGGGQYSAKPRNQGGYGGSSSSSSSGSGRRF | Martin, E. W., Holehouse, A. S., Peran, I., Farag, M., Incicco, J. J., Bremer, A., Grace, C. R., Soranno, A., Pappu, R. V., & Mittag, T. (2020). Valence and patterning of aromatic residues determine the phase behavior of prion-like domains. Science, 367(6478), 694–699. |
| A1_Aro_minus_minus | 29.3 | 0.5 | GSMASASSSQRGRSGSGNSGGGRGGGFGGNDNSGRGGNSSGRGGFGGSRGGGGSGGSGDGYNGSGNDGSNSGGGGSSNDFGNSNNQSSNSGP | Martin, E. W., Holehouse, A. S., Peran, I., Farag, M., Incicco, J. J., Bremer, A., Grace, C. R., Soranno, A., Pappu, R. V., & |

| | | | | |
|---|---|---|---|---|
| | | | MKGGNFGGRSSGGSGGGGQYSAKPRNQGGSGGSSSSSSSGSGRRS | Mittag, T. (2020). Valence and patterning of aromatic residues determine the phase behavior of prion-like domains. Science, 367(6478), 694–699. |
| A1_Aro_plus | 24.2 | 1.5 | GSMAFASSFQRGRYGSGNFGGGRGGGFGGNDNFGRGGNFSGRGGFGGSRGGGGYGGSGDGYNGFGNDGSNFGGGGSYNDFGNYNNQSSNFGPMKGGNFGGRSSGGSYGGGQYFAKPRNQGGYGGSSFSSSYGSGRRF | Martin, E. W., Holehouse, A. S., Peran, I., Farag, M., Incicco, J. J., Bremer, A., Grace, C. R., Soranno, A., Pappu, R. V., & Mittag, T. (2020). Valence and patterning of aromatic residues determine the phase behavior of prion-like domains. Science, 367(6478), 694–699. |
| HeV_PNT3_CTD_200_254 | 28 | 0 | MSYYHHHHHHLESTSLYKKAGFTPTEEPPVIPEYYYGSGRRGDLSKSPPRGNVNLDSIKIYTSDDEDENQLEYEDEF | Nilsson, J. F., Baroudi, H., Gondelaud, F., Pesce, G., Bignon, C., Ptchelkine, D., Chamieh, J., Cottet, H., Kajava, A. V., & Longhi, S. (2022). Molecular Determinants of Fibrillation in a Viral Amyloidogenic Domain from Combined Biochemical and Biophysical Studies. International Journal of Molecular Sciences, 24(1). https://doi.org/10.3390/ijms24010399 |
| HeV_PNT3_200_310_YYY_AAA | 40 | 0 | MSYYHHHHHHLESTSLYKKAGFTPTEEPPVIPEAAAGSGRRGDLSKSPPRGNVNLDSIKIYTSDDEDENQLEYEDEFAKSSSEVVIDTTPEDNDSINQEEVVGDPSDQGLEHPFPLGKFPEKEETPDVRRKDS | Nilsson, J. F., Baroudi, H., Gondelaud, F., Pesce, G., Bignon, C., Ptchelkine, D., Chamieh, J., Cottet, H., Kajava, A. V., & Longhi, S. (2022). Molecular Determinants of Fibrillation in a Viral Amyloidogenic Domain from Combined Biochemical and Biophysical Studies. International Journal of |

| | | | | |
|---|---|---|---|---|
| | | | | Molecular Sciences, 24(1). https://doi.org/10.3390/ijms24010399 |
| HeV_PNT3_200_310_WT | 37 | 0 | MSYYHHHHHHLESTSLYKKAGSTPTEEPPVIPEYYYGSGRRGDLSKSPPRGNVNLDSIKIYTSDDEDENQLEYEDEFAKSSSEVVIDTTPEDNDSINQEEVVGDPSDQGLEHPFPLGKFPEKEETPDVRRKDS | Nilsson, J. F., Baroudi, H., Gondelaud, F., Pesce, G., Bignon, C., Ptchelkine, D., Chamieh, J., Cottet, H., Kajava, A. V., & Longhi, S. (2022). Molecular Determinants of Fibrillation in a Viral Amyloidogenic Domain from Combined Biochemical and Biophysical Studies. International Journal of Molecular Sciences, 24(1). https://doi.org/10.3390/ijms24010399 |
| NiV_PNT3_200_314_WT | 37 | 0 | MSYYHHHHHHLESTSLYKKAGFDPAKDSPVIAEHYYGLGVKEQNVGPQTSRNVNLDSIKLYTSDDEEADQLEFEDEFAGSSSEVIVGISPEDEEPSSVGGKPNESIGRTIEGQSIRDNLQAKDNKSTDVPGAGPKDS | Nilsson, J. F., Baroudi, H., Gondelaud, F., Pesce, G., Bignon, C., Ptchelkine, D., Chamieh, J., Cottet, H., Kajava, A. V., & Longhi, S. (2022). Molecular Determinants of Fibrillation in a Viral Amyloidogenic Domain from Combined Biochemical and Biophysical Studies. International Journal of Molecular Sciences, 24(1). https://doi.org/10.3390/ijms24010399 |
| red1_288_345 | 25 | 0 | GAMGISLPLLKQDDWLSSSKPFGSSTPNVVIEFDSDDDGDDFSNSKIEQSNLEKPPSNSENGGSHHHHHH | TBD |
| p150L_342_475 | 41 | 0 | MAERLGKQLKLRAEREEKEKLKEEAKRAKEEAKKKKEEEKELKEKERREKREKDEKEKAEKQRLKEERRKERQEALEAKLEEKRKKEEEKRLREEEKRIK | Gopinathan Nair, A., Rabas, N., Lejon, S., Homiski, C., Osborne, M. J., Cyr, N., |

| | | | | |
|---|---|---|---|---|
| | | | AEKAEITRFFQKPKTPQAPKTLAGSCGKFAPFEIKELEHHHHHH | Sverzhinsky, A., Melendy, T., Pascal, J. M., Laue, E. D., Borden, K. L. B., Omichinski, J. G., & Verreault, A. (2022). Unorthodox PCNA Binding by Chromatin Assembly Factor 1. International Journal of Molecular Sciences, 23(19). https://doi.org/10.3390/ijms231911099 |
| E1A_2022 | 36 | 0 | GSMSHFEPPTLHELYDLDVTAPEDPNEEAVSQIFPDSVMLAVQEGIDLLTFPPAPGSPEPPHLSRQPEQPEQRALGPVSMPNLVPEVIDLYCYEQLNPPSDDEDEEGEEFVLDY | González-Foutel, N. S., Glavina, J., Borcherds, W. M., Safranchik, M., Barrera-Vilarmau, S., Sagar, A., Estaña, A., Barozet, A., Garrone, N. A., Fernandez-Ballester, G., Blanes-Mira, C., Sánchez, I. E., de Prat-Gay, G., Cortés, J., Bernadó, P., Pappu, R. V., Holehouse, A. S., Daughdrill, G. W., & Chemes, L. B. (2022). Conformational buffering underlies functional selection in intrinsically disordered protein regions. Nature Structural & Molecular Biology, 29(8), 781–790. |
| RelA_TAD | 27 | 0 | MGSVPKPAPQPYTFPASLSTINFDEFSPMLLPSGQISNQALALAPSSAPVLAQTMVPSSAMVPLAQPPAPAPVLTPGPPQSLSAPVPKSTQAGEGTLSEALLHLQFDADEDLGALLGNSTDPGVFTDLASVDNSEFQQLLNQGVSMSHSTAEPMLMEYPEAITRLVTGSQRPPDPAPTPLGTSGLPNGLSGDEDFSSIADMDFSALLSQISSLEHHHHHH | Baughman, H. E. R., Narang, D., Chen, W., Villagrán Suárez, A. C., Lee, J., Bachochin, M. J., Gunther, T. R., Wolynes, P. G., & Komives, E. A. (2022). An intrinsically disordered transcription activation domain increases the DNA binding affinity and reduces the specificity of NFκB p50/RelA. The Journal of Biological Chemistry, 298(9), 102349. |

| | | | | |
|---|---|---|---|---|
| EIF_450_1_249 | 52 | 0 | GSMTDETAHPTQSASKQESAALKQTGDDQQESQQQRGYTNYNNGSNYTQKKPYNSNRPHQQRGGKFGPNRYNNRGNYNGGGSFRGGHMGANSSNVPWTGYYNNYPVYYQPQQMAAAGSAPANPIPVEEKSPVPTKIEITTKSGEHLDLKEQHKAKLQSQERSTVSPQPESKLKETSDSTSTSTPTPTPSTNDSKASSEENISEAEKTRRNFIEQVKLRKAALEKKRKEQLEGSSGNNNIPMKTTPENVEEK | Chaves-Arquero, B., Martínez-Lumbreras, S., Sibille, N., Camero, S., Bernadó, P., Jiménez, M. Á., Zorrilla, S., & Pérez-Cañadillas, J. M. (2022). eIF4G1 N-terminal intrinsically disordered domain is a multi-docking station for RNA, Pab1, Pub1, and self-assembly. Frontiers in Molecular Biosciences, 9, 986121. |
| TIF2_624_774 | 37 | 0 | ERADGQSRLHDSKGQTKLLQLLTTKSDQMEPSPLASSLSDTNKDSTGSLPGSGSTHGTSLKEKHKILHRLLQDSSSPVDLAKLTAEATGKDLSQESSSTAPGSEVTIKQEPVSPKKKENALLRYLLDKDDTKDIGLPEITPKLERLDSKT | Senicourt, L., le Maire, A., Allemand, F., Carvalho, J. E., Guee, L., Germain, P., Schubert, M., Bernadó, P., Bourguet, W., & Sibille, N. (2021). Structural insights into the interaction of the intrinsically disordered co-activator TIF2 with retinoic acid receptor heterodimer (RXR/RAR). Journal of Molecular Biology, 433(9), 166899. |
| IR_CTD | 38 | 0 | GPRRNQPAEQTTTTTTHTVVQQQTGGNTPAQGGTDATRAEDASLNRRDSQGSVASTHWSDSSSEVVNPYAEVGGARNSLSAHQPEEHIYDEVAADPGYSVIQNFSGSGPVTGRLIGTPGQGIQSTYALLANSGGLRLGMGGLTSGGESAVSSVNAAPTPGPVRFVWSHPQFEK | TBD |
| Tau_ht35_2022 | 46 | 0 | EPPKSGDRSGYSSPGSPGTPGSRSRTPSLPTPPTREPKKVAVVRTPPKSPSSAKSRLQTAPVPMPDLKNVKSKIGSTENLKHQPGGGKVQIINKKLDLSNVQSKCGSKDNIKHVPGGGSVQIVYKPVDLSKVTSKCGSLGNIHHKPGGGQVEVKSEKLDFKDRVQSKIGSLDNITHVPGGGNKKIETHKLTFRENAKAKTDH | Lyu, C., Da Vela, S., Al-Hilaly, Y., Marshall, K. E., Thorogate, R., Svergun, D., Serpell, L. C., Pastore, A., & Hanger, D. P. (2021). The Disease Associated Tau35 Fragment has an Increased Propensity to |

| | | | | |
|---|---|---|---|---|
| | | | GAEIVYKSPVVSGDTSPRHLSNVSSTGSIDMVDSPQLATLADEVSASLAKQGL | Aggregate Compared to Full-Length Tau. Frontiers in Molecular Biosciences, 8, 779240. |
| Tau_ht410_2N3R | 63 | 0 | MAEPRQEFEVMEDHAGTYGLGDRKDQGGYTMHQDQEGDTDAGLKESPLQTPTEDGSEEPGSETSDAKSTPTAEDVTAPLVDEGAPGKQAAAQPHTEIPEGTTAEEAGIGDTPSLEDEAAGHVTQARMVSKSKDGTGSDDKKAKGADGKTKIATPRGAAPPGQKGQANATRIPAKTPPAPKTPPSSGEPPKSGDRSGYSSPGSPGTPGSRSRTPSLPTPPTREPKKVAVVRTPPKSPSSAKSRLQTAPVPMPDLKNVKSKIGSTENLKHQPGGGKVQIVYKPVDLSKVTSKCGSLGNIHHKPGGGQVEVKSEKLDFKDRVQSKIGSLDNITHVPGGGNKKIETHKLTFRENAKAKTDHGAEIVYKSPVVSGDTSPRHLSNVSSTGSIDMVDSPQLATLADEVSASLAKQGL | Lyu, C., Da Vela, S., Al-Hilaly, Y., Marshall, K. E., Thorogate, R., Svergun, D., Serpell, L. C., Pastore, A., & Hanger, D. P. (2021). The Disease Associated Tau35 Fragment has an Increased Propensity to Aggregate Compared to Full-Length Tau. Frontiers in Molecular Biosciences, 8, 779240. |
| Tau_ht410_2N4R | 67 | 0 | MAEPRQEFEVMEDHAGTYGLGDRKDQGGYTMHQDQEGDTDAGLKESPLQTPTEDGSEEPGSETSDAKSTPTAEDVTAPLVDEGAPGKQAAAQPHTEIPEGTTAEEAGIGDTPSLEDEAAGHVTQARMVSKSKDGTGSDDKKAKGADGKTKIATPRGAAPPGQKGQANATRIPAKTPPAPKTPPSSGEPPKSGDRSGYSSPGSPGTPGSRSRTPSLPTPPTREPKKVAVVRTPPKSPSSAKSRLQTAPVPMPDLKNVKSKIGSTENLKHQPGGGKVQIINKKLDLSNVQSKCGSKDNIKHVPGGGSVQIVYKPVDLSKVTSKCGSLGNIHHKPGGGQVEVKSEKLDFKDRVQSKIGSLDNITHVPGGGNKKIETHKLTFRENAKAKTDHGAEIVYKSPVVSGDTSPRHLSNVSSTGSIDMVDSPQLATLADEVSASLAKQGL | Lyu, C., Da Vela, S., Al-Hilaly, Y., Marshall, K. E., Thorogate, R., Svergun, D., Serpell, L. C., Pastore, A., & Hanger, D. P. (2021). The Disease Associated Tau35 Fragment has an Increased Propensity to Aggregate Compared to Full-Length Tau. Frontiers in Molecular Biosciences, 8, 779240. |
| SMAD_linker | 29 | 0 | GPLPPVLVPRHTEILTELPPLDDYTHSIPENTNFPAGIEPQSNYIPETPPPGYISEDGETSDQQLNQSMDTGSPAELSPTTLSPVNHSLD | Gomes, T., Martin-Malpartida, P., Ruiz, L., Aragón, E., Cordeiro, T. N., & Macias, M. J. (2021). Conformational landscape of multidomain SMAD proteins. Computational |

| | | | | |
|---|---|---|---|---|
| | | | | and Structural Biotechnology Journal, 19, 5210–5224. |
| MenV_LBD | 25 | 0 | TTIKIMDPGVGDGATAAKSKRLFKEAPVVVSGPVIGDNPIVDADTIQLDELARPSLPKTKSQ | Webby, M. N., Herr, N., Bulloch, E. M. M., Schmitz, M., Keown, J. R., Goldstone, D. C., & Kingston, R. L. (2021). Structural Analysis of the Menangle Virus P Protein Reveals a Soft Boundary between Ordered and Disordered Regions. Viruses, 13(9). https://doi.org/10.3390/v13091737 |
| syndecan3_ED | 65 | 0 | MGSSHHHHHHSSGLVPRGSMAQRWRSENFERPVDLEGSGDDDSFPDDELDDLYSGSGSGYFEQESGIETAMETRFSPDVALAVSTTPAVLPTTNIQPVGTPFEELPSERPTLEPATSPLVVTEVPEEPSQRATTVSTTMETATTAATSTGDPTVATVPATVATATPSTPAAPPFTATTAVIRTTGVRRLLPLPLTTVATARATTPEAPSPPTTAAVLDTEAPTPRLVSTATSRPRALPRPATTQEPDIPERSTLPLGTTAPGPTEVAQTPTPETFLTTIRDEPEVPVSGGPSGDFELPEEETTQPDTANEVVAVGGAAAKASSPPGTLPKGARPGPGLLDNAIDSGSSAAQLPQKSILERKEVLVDYKDDDDK | Gondelaud, F., Bouakil, M., Le Fèvre, A., Miele, A. E., Chirot, F., Duclos, B., Liwo, A., & Ricard-Blum, S. (2021). Extended disorder at the cell surface: The conformational landscape of the ectodomains of syndecans. Matrix Biology Plus, 12, 100081. |
| syndecan4 | 42 | 0 | GSSHHHHHHSSGLVPRGSHMESIRETEVIDPQDLLEGRYFSGALPDDEDVVGPGQESDDFELSGSGDLDDLEDSMIGPEVVHPLVPLDNHIPERAGSGSQVPTEPKKLEENEVIPKRISPVEESEDVSNKVSMSSTVQGSNIFERTEVLAGCPEHDYKDDDDK | Gondelaud, F., Bouakil, M., Le Fèvre, A., Miele, A. E., Chirot, F., Duclos, B., Liwo, A., & Ricard-Blum, S. (2021). Extended disorder at the cell surface: The conformational landscape of the ectodomains of syndecans. Matrix Biology Plus, 12, 100081. |
| N_FATZ_1 | 35 | 0 | MAHHHHHHVDDDDKIMPLSGTPAPNKKRKSSKLIMELTGGGQESSGLNLGKKISVPRDVMLEELSLLTNRGSKMFKLRQMRVEKFIYENHPDVFSDSSMD | Sponga, A., Arolas, J. L., Schwarz, T. C., Jeffries, C. M., Rodriguez Chamorro, A., |

| | | | | |
|---|---|---|---|---|
| | | | HFQKFLPTVGGQLGTAGQGFSYSKSNGRGGSQAGGSGSAGQYGSDQQHHLGSGSGAGGTGGPAGQAGRGGAAGTAGVGETGSGDQAGGEAE | Kostan, J., Ghisleni, A., Drepper, F., Polyansky, A., De Almeida Ribeiro, E., Pedron, M., Zawadzka-Kazimierczuk, A., Mlynek, G., Peterbauer, T., Doto, P., Schreiner, C., Hollerl, E., Mateos, B., Geist, L., … Djinović-Carugo, K. (2021). Order from disorder in the sarcomere: FATZ forms a fuzzy but tight complex and phase-separated condensates with α-actinin. Science Advances, 7(22). https://doi.org/10.1126/sciadv.abg7653 |
| DeltaN_FATZ_1 | 39 | 0 | GPTVGGQLGTAGQGFSYSKSNGRGGSQAGGSGSAGQYGSDQQHHLGSGSGAGGTGGPAGQAGRGGAAGTAGVGETGSGDQAGGEGKHITVFKTYISPWERAMGVDPQQKMELGIDLLAYGAKAELPKYKSFNRTAMPYGGYEKASKRMTFQMPKFDLGPLLSEPLVLYNQNLSNRPSFNRTPIPWLSSGEPVDYNVDIGIPLDGETEEL | Sponga, A., Arolas, J. L., Schwarz, T. C., Jeffries, C. M., Rodriguez Chamorro, A., Kostan, J., Ghisleni, A., Drepper, F., Polyansky, A., De Almeida Ribeiro, E., Pedron, M., Zawadzka-Kazimierczuk, A., Mlynek, G., Peterbauer, T., Doto, P., Schreiner, C., Hollerl, E., Mateos, B., Geist, L., … Djinović-Carugo, K. (2021). Order from disorder in the sarcomere: FATZ forms a fuzzy but tight complex and phase-separated condensates with α-actinin. Science Advances, 7(22). https://doi.org/10.1126/sciadv.abg7653 |
| histatin_2021 | 15 | 0 | DSHAKRHHGYKRKFHEKHHSHRGY | Sagar, A., Jeffries, C. M., Petoukhov, M. V., Svergun, D. I., & Bernadó, P. (2021). Comment on the Optimal Parameters to Derive Intrinsically Disordered Protein |

| | | | | |
|---|---|---|---|---|
| | | | | Conformational Ensembles from Small-Angle X-ray Scattering Data Using the Ensemble Optimization Method. Journal of Chemical Theory and Computation, 17(4), 2014–2021. |
| synthELP | 66 | 0 | GGVPGAIPGGVPGGVFYPGAGLGALGGGALGPGGKPLKPVPGGLAGAGLGAGLGAFPAVTFPGALVPGGVADAAAAYKAAKAGAGLGGVPGVGGLGVSAGAVVPQPGAGVKPGKVPGVGLPGVYPGGVLPGARFPGVGVLPGVPTGAGVKPKAPGVGGAFAGIPGVGPFGGPQPGVPLGYPIKAPKLPGGYGLPYTTGKLPYGYGPGGVAGAAGKAGYPTGTGVGPQAAAAAAAKAAAKFGAGAAGVLPGVGGAGVPGVPGAIPGIGGIAGVGTPAAAAAAAAAAKAAKYGAAAGLVPGGPGFGPGVVGVPGAGVPGVGVPGAGIPVVPGAGIPGAAVPGVVSPEAAAKAAAKAAKYGARPGVGVGGIPTYGVGAGGFPGFGVGVGGIPGVAGVPGVGGVPGVGGVPGVGISPEAQAAAAAKAAKYGVGTPAAAAAKAAAKAAQFGLVPGVGVAPGVGVAPGVGVAPGVGLAPGVGVAPGVGVAPGVGVAPGIGPGGVAAAAKSAAKVAAKAQLRAAAGLGAGIPGLGVGVGVPGLGVGAGVPGLGVGAGVPGFGAVPGALAAAKAAKYGAAVPGVLGGLGALGGVGIPGGVVGAGPAAAAAAAKAAAKAAQFGLVGAAGLGGLGVGGLGVPGVGGLGGIPPAAAAKAAKYGAAGLGGVLGGAGQFPLGGVAARPGFGLSPIFPGGACLGKACGRKRK | Lockhart-Cairns, M. P., Newandee, H., Thomson, J., Weiss, A. S., Baldock, C., & Tarakanova, A. (2020). Transglutaminase-mediated cross-linking of tropoelastin to fibrillin stabilises the elastin precursor prior to elastic fibre assembly. Journal of Molecular Biology, 432(21), 5736–5751. |
| UL11 | 24 | 0 | MGLSFSGTRPCCCRNNVLITDDGEVVSLTAHDFDVVDIESEEEGNFYVPPDMRGVTRAPGRQRLRSSDPPSRHTHRRTPGGACPATQFPPPMSDSEWSHPQFEK | Metrick, C. M., Koenigsberg, A. L., & Heldwein, E. E. (2020). Conserved Outer Tegument Component UL11 from Herpes Simplex Virus 1 Is an Intrinsically Disordered, RNA-Binding Protein. mBio, 11(3). https://doi.org/10.1128/mBio.00810-20 |

| | | | | |
|---|---|---|---|---|
| GON7_NTD | 31 | 0 | MGHHHHHHENLYFQGELLGEYVGQEGKPQKLRVSCEAPGDGDPFQGLLSGVAQMKDMVTELFDPLVQGEVQHRVAAAPDEDLDGDDEDDAEDENNIDNRTNFDGPSAKRPKTPS | Arrondel, C., Missoury, S., Snoek, R., Patat, J., Menara, G., Collinet, B., Liger, D., Durand, D., Gribouval, O., Boyer, O., Buscara, L., Martin, G., Machuca, E., Nevo, F., Lescop, E., Braun, D. A., Boschat, A.-C., Sanquer, S., Guerrera, I. C., … Mollet, G. (2019). Defects in t6A tRNA modification due to GON7 and YRDC mutations lead to Galloway-Mowat syndrome. Nature Communications, 10(1), 3967. |
| Bmal1_CTD_P624A | 28 | 0 | GPDASSPGGKKILNGGTPDIPSTGLLPGQAQETPGYPYSDSSSILGENPHIGIDMIDNDQGSSSPSNDEAAMAVIMSLLEADAGLGGPVDFSDLPWAL | Garg, A., Orru, R., Ye, W., Distler, U., Chojnacki, J. E., Köhn, M., Tenzer, S., Sönnichsen, C., & Wolf, E. (2019). Structural and mechanistic insights into the interaction of the circadian transcription factor BMAL1 with the KIX domain of the CREB-binding protein. The Journal of Biological Chemistry, 294(45), 16604–16619. |
| NID_2059_2325 | 47 | 0 | GPHMQVPRTHRLITLADHICQIITQDFARNQVPSQASTSTFQTSPSALSSTPVRTKTSSRYSPESQSQTVLHPRPGPRVSPENLVDKSRGSRPGKSPERSHIPSEPYEPISPPQGPAVHEKQDSMLLLSQRGVDPAEQRSDSRSPGSISYLPSFFTKLESTSPMVKSKKQEIFRKLNSSGGGDSDMAAAQPGTEIFNLPAVTTSGAVSSRSHSFADPASNLGLEDIIRKALMGSFDDKVEDHGVVMSHPVGIMPGSASTSVVTSSEARRDE | Cordeiro, T. N., Sibille, N., Germain, P., Barthe, P., Boulahtouf, A., Allemand, F., Bailly, R., Vivat, V., Ebel, C., Barducci, A., Bourguet, W., le Maire, A., & Bernadó, P. (2019). Interplay of protein disorder in retinoic acid receptor heterodimer and its corepressor regulates gene expression. Structure , 27(8), 1270–1285.e6. |

| | | | | |
|---|---|---|---|---|
| MAP2c | 67 | 0 | MADERKDEGKAPHWTSASLTEAAAHPHSPEMK DQGGSGEGLSRSANGFPYREEEEGAFGEHGSQ GTYSDTKENGINGELTSADRETAEEVSARIVQVV TAEAVAVLKGEQEKEAQHKDQPAALPLAAEETVN LPPSPPPSPASEQTAALEEATSGESAQAPSAFKQ AKDKVTDGITKSPEKRSSLPRPSSILPPRRGVSG DREENSFSLNSSISSARRTTRSEPIRRAGKSGTS TPTTPGSTAITPGTPPSYSSRTPGTPGTPSYPRT PGTPKSGILVPSEKKVAIIRTPPKSPATPKQLRLIN QPLPDLKNVKSKIGSTDNIKYQPKGGQVQIVTKKI DLSHVTSKCGSLKNIRHRPGGGRVKIESVKLDFK EKAQAKVGSLDNAHHVPGGGNVKIDSQKLNFRE HAKARVDHGAEIITQSPSRSSVASPRRLSNVSSS GSINLLESPQLATLAEDVTAALAKQGL | Melková, K., Zapletal, V., Jansen, S., Nomilner, E., Zachrdla, M., Hritz, J., Nováček, J., Zweckstetter, M., Jensen, M. R., Blackledge, M., & Žídek, L. (2018). Functionally specific binding regions of microtubule-associated protein 2c exhibit distinct conformations and dynamics. The Journal of Biological Chemistry, 293(34), 13297–13309. |
| TRF2_BR | 17 | 0 | GPPGSMAGGGGSSDGSGRAAGRRASRSSGRA RRGRHEPGLGGPAERGAG | Necasová, I., Janoušková, E., Klumpler, T., & Hofr, C. (2017). Basic domain of telomere guardian TRF2 reduces D-loop unwinding whereas Rap1 restores it. Nucleic Acids Research, 45(21), 12170–12180. |
